# The ancestral animal genetic toolkit revealed by diverse choanoflagellate transcriptomes

**DOI:** 10.1101/211789

**Authors:** Daniel J. Richter, Parinaz Fozouni, Michael B. Eisen, Nicole King

## Abstract

The changes in gene content that preceded the origin of animals can be reconstructed by comparison with their sister group, the choanoflagellates. However, only two choanoflagellate genomes are currently available, providing poor coverage of their diversity. We sequenced transcriptomes of 19 additional choanoflagellate species to produce a comprehensive reconstruction of the gains and losses that shaped the ancestral animal gene repertoire. We find roughly 1,700 gene families with origins on the animal stem lineage, of which only a core set of 36 are conserved across animals. We find more than 350 gene families that were previously thought to be animal-specific actually evolved before the animal-choanoflagellate divergence, including Notch and Delta, Toll-like receptors, and glycosaminoglycan hydrolases that regulate animal extracellular matrix (ECM). In the choanoflagellate *Salpingoeca helianthica*, we show that a glycosaminoglycan hydrolase modulates rosette colony size, suggesting a link between ECM regulation and morphogenesis in choanoflagellates and animals.

**Data Availability:** Raw sequencing reads: NCBI BioProject PRJNA419411 (19 choanoflagellate transcriptomes), PRJNA420352 (*S. rosetta* polyA selection test)

Transcriptome assemblies, annotations, and gene families: https://dx.doi.org/10.6084/m9.figshare.5686984

Protocols: https://dx.doi.org/10.17504/protocols.io.kwscxee

## Introduction

The biology of the first animal, the “Urmetazoan,” has fascinated and confounded biologists for more than a century (Dujardin, 1841; James-Clark, 1867; Haeckel, 1869, 1873, 1874; Kent, 1880; Leadbeater and McCready, 2000). What features defined the biology of the Urmetazoan, and which of those features represent animal innovations? Despite the fact that the first animals originated over 600 million years ago (Douzery et al., 2004; Hedges et al., 2004; Peterson et al., 2004; Narbonne, 2005; Knoll, 2011) and left no fossil evidence, features of their genomes can be reconstructed through phylogenetically-informed comparisons among extant animals, their closest living relatives, the choanoflagellates, and other closely related lineages (King, 2004; Rokas, 2008; Richter and King, 2013; Grau-Bové et al., 2017; Sebé-Pedrós et al., 2017).

Although over 1,000 genomes of animals have been sequenced (NCBI Resource Coordinators, 2017), only two choanoflagellate genomes have been previously published. These two choanoflagellates are the strictly unicellular *Monosiga brevicollis* and the emerging model choanoflagellate *Salpingoeca rosetta*, which differentiates into a number of cellular states, ranging from single cells to multicellular rosette colonies (Fairclough et al., 2010; Dayel et al., 2011). The *M. brevicollis* and *S. rosetta* genomes revealed that many genes critical for animal biology, including p53, Myc, cadherins, C-type lectins, and diverse tyrosine kinases, evolved before the divergence of animals and choanoflagellates (King et al., 2008; Fairclough et al., 2013), whereas many other genes essential for animal biology, including collagen, and components of the Wnt, Notch/Delta, Hedgehog, TGF-β, and innate immune pathways (e.g., Toll-like receptors) have not been detected in choanoflagellates, and therefore have been considered textbook examples of animal innovations.

Nonetheless, *M. brevicollis* and *S. rosetta* are relatively closely related (Carr et al., 2017), leaving the bulk of choanoflagellate diversity unexplored. Moreover, both species have demonstrably experienced gene loss, as some genes conserved between animals and non-choanoflagellates are apparently missing from *M. brevicollis* and *S. rosetta*. Examples include RNAi pathway components, which are present across eukaryotes (Shabalina and Koonin, 2008), and the cell adhesion protein β-integrin and T-box and Runx transcription factor families, which have been detected in the filasterean *Capsaspora owczarzaki* (Sebé-Pedrós and Ruiz-Trillo, 2010; Sebé-Pedrós et al., 2010, 2011, 2013a; Ferrer-Bonet and Ruiz-Trillo, 2017). Gene loss can lead to false negatives during ancestral genome reconstruction, and the phenomenon in choanoflagellates parallels that of animals, where two species selected for early genome projects, *Drosophila melanogaster* and *Caenorhabditis elegans*, were later found to have lost numerous genes (e.g., Hedgehog and NF-*κ*B in *C. elegans* and fibrillar collagens in both *C. elegans* and *D. melanogaster)* that are critical for animal development and otherwise conserved across animal diversity (C. elegans Sequencing Consortium, 1998; Aspöck et al., 1999; Gilmore, 1999; Rubin et al., 2000).

To counteract the impact of gene loss in *M. brevicollis* and *S. rosetta*, and gain a more complete picture of the Urmetazoan gene catalog, we analyzed the protein coding genes of 19 previously unsequenced species of choanoflagellates representing each major known lineage (Carr et al., 2017). By comparing their gene catalogs with those of diverse animals and other phylogenetically relevant lineages, we have greatly expanded and refined our understanding of the genomic heritage of animals. This more comprehensive data set revealed that ~372 gene families that were previously thought to be animal-specific actually evolved prior to the divergence of choanoflagellates and animals, including gene families required for animal development (e.g., Notch/Delta), immunity (e.g., Toll-like receptors) and extracellular matrix biology (glycosaminoglycan hydrolase). We find that an additional ~1,718 gene families evolved along the animal stem lineage, many of which likely underpin unique aspects of animal biology. Interestingly, most of these animal-specific genes were subsequently lost from one or more species, leaving only 36 core animal-specific genes that were conserved in all animals within our data set. Finally, we demonstrate that a glycosaminoglycan hydrolase involved in remodeling animal extracellular matrix is conserved among choanoflagellates and animals and regulates rosette colony morphogenesis in the choanoflagellate *Salpingoeca helianthica*.

## Results

### The phylogenetic distribution of animal and choanoflagellate gene families

To reconstruct the genomic landscape of animal evolution, we first cataloged the protein coding potential of nineteen diverse choanoflagellate species by sequencing and assembling their transcriptomes (Figure 1, Supp. Figure 1, Supp. Table 1). Because most of these species were previously little-studied in the laboratory, two important stages of this project were the establishment of growth conditions optimized for each species and the development of improved protocols for isolating choanoflagellate mRNA for cDNA library construction and sequencing (Supp. Table 1, Methods).

**Figure 1.**
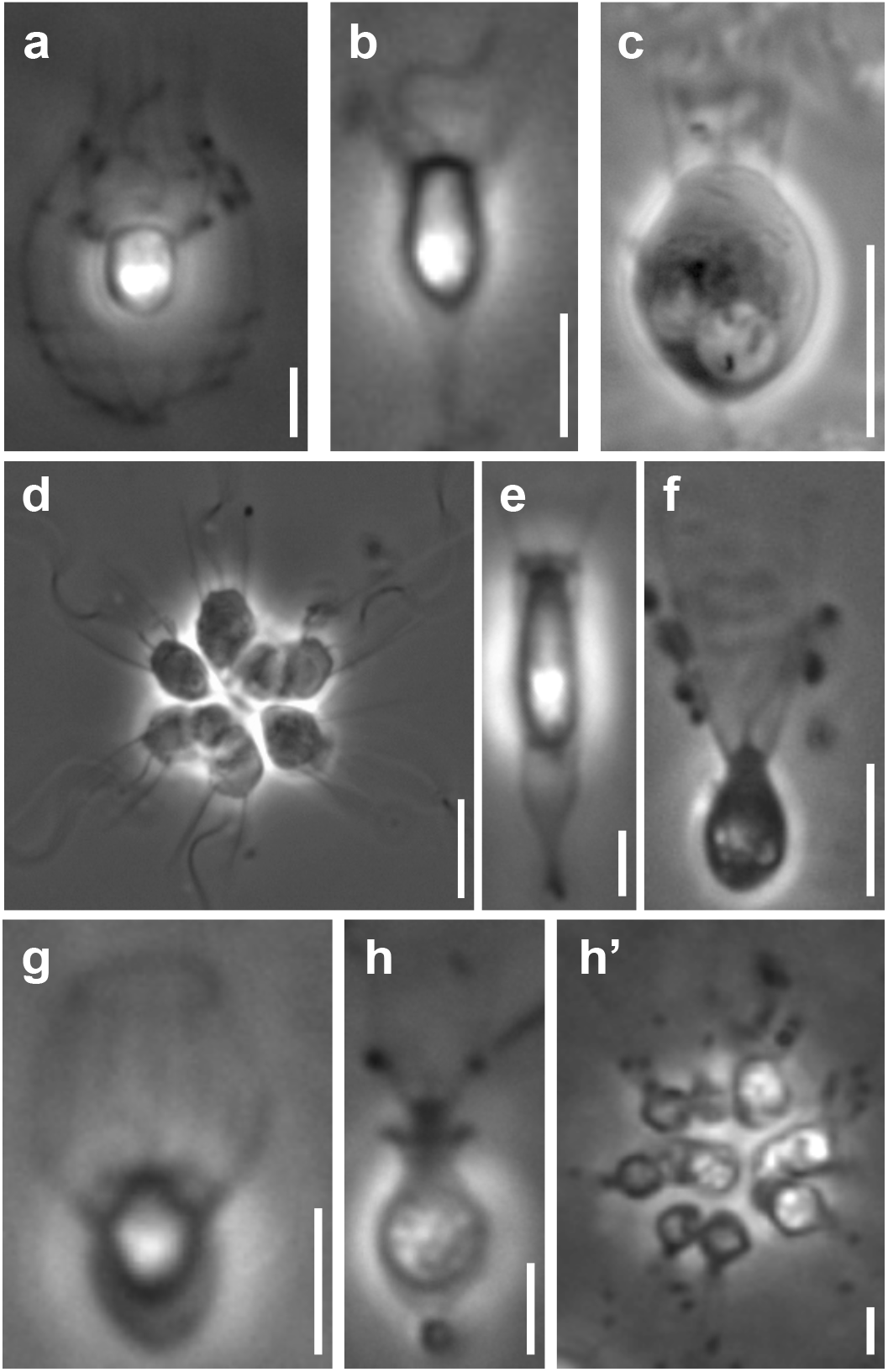
Representative choanoflagellates analyzed in this study. Choanoflagellates have diverse morphologies, including single cells, multicellular colonies, and the production in some lineages of extracellular structures. (a) *Diaphanoeca grandis*, within a silica-based extracellular structure called a ‘lorica’. (b) *Acanthoeca spectabilis*, within lorica. (c) *Codosiga hollandica*, with a basal organic stalk. (d) A rosette colony of *Salpingoeca rosetta*; image courtesy of Mark Dayel. (e) *Salpingoeca dolichothecata*, within an organic extracellular structure called a ‘theca’. (f) *Mylnosiga fluctuans*, with no extracellular structure. (g) *Didymoeca costata*, within lorica. (h) *Salpingoeca helianthica*, within theca. (h’) A rosette colony of *S. helianthica*. All scale bars represent 5 *μ*m. Prey bacteria are visible in most panels as small black dots.

Using multiple independent metrics, we found that the new choanoflagellate transcriptomes approximate the completeness of choanoflagellate genomes for the purposes of cataloging protein-coding genes (Supp. Figure 2a). For example, the genomes of *M. brevicollis* and *S. rosetta* contain 83% and 89%, respectively, of a benchmark set of conserved eukaryotic genes [BUSCO; (Simão et al., 2015)], whereas each of the new choanoflagellate transcriptomes contains between 88 - 96% (Supp. Table 2). In addition, our analyses of conserved proteins encoded by each of the choanoflagellate gene catalogs revealed that the average phylogenetic distance between pairs of choanoflagellates is slightly larger than the phylogenetic distance between the mouse *Mus musculus* and the sponge *Amphimedon queenslandica*, indicating that the phylogenetic diversity of choanoflagellates is comparable to that of animals (Supp. Figure 3).

We next compared the choanoflagellate gene catalogs against those of diverse animals and phylogenetically relevant outgroups (Supp. Table 3) to identify gene families and determine the ancestry of genes present in animals. Two features that distinguish our analyses from prior reconstructions of ancestral animal gene content are (1) the additional breadth and depth provided by 19 phylogenetically diverse and newly-sequenced choanoflagellates and (2) a probabilistic and phylogenetically-informed approach designed to avoid the artificial inflation of ancestral gene content resulting from methods that rely on binary decisions for gene family presence or absence in a species (Supp. Figure 4, Methods).

By grouping gene families by their phylogenetic distribution on a heat map, we were able to visualize and infer their evolutionary history, as well as their presence or absence in each species analyzed (Figure 2, Supp. Figure 5, Supp. Figure 6, Supp. Table 4, Supp. Table 5). Several notable observations emerge from this visualization. First, the origins of animals, choanoflagellates, and choanozoans [the monophyletic group composed of animals and choanoflagellates (Brunet and King, 2017)] were each accompanied by the evolution of distinct sets of gene families (i.e., synapomorphies), some of which likely underpin their unique biological features. Second, the amount of gene family evolution that occurred on the animal and choanoflagellate stem lineages is roughly equivalent, indicating that the specific functions of novel gene families, not their quantity, was critical to the divergence of animals from their sister group. Finally, although different sets of gene families can reliably be inferred to have been present in the last common ancestor of each group, gene family loss was rampant during animal and choanoflagellate diversification. [After these analyses were complete, several additional genomes from early-branching holozoans and animals became available. Incorporating them *post hoc* into the heat map did not substantially change any of the above observations (Supp. Figure 7; Methods).]

**Figure 2.**
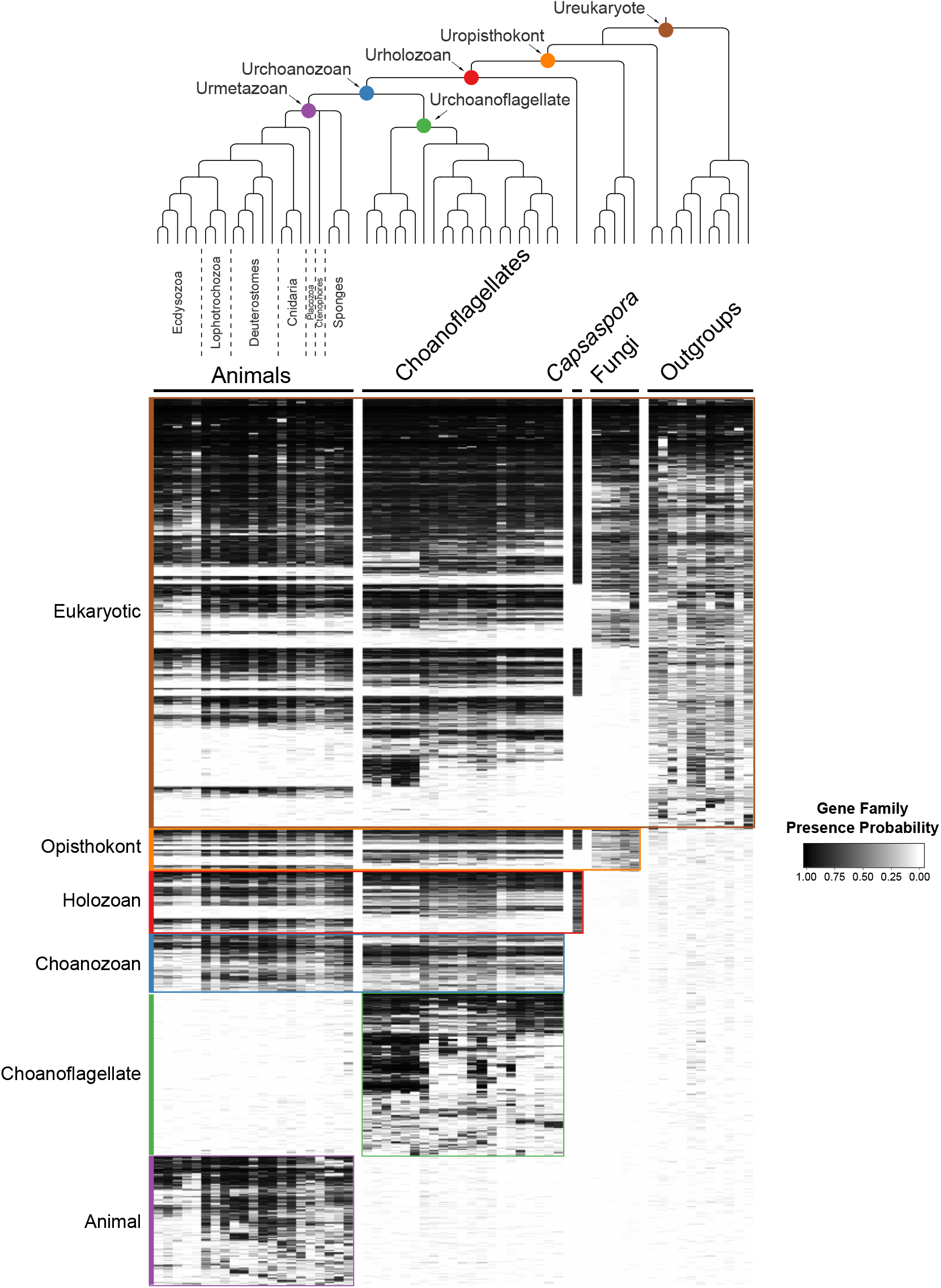
The evolution of gene families in animals, choanoflagellates and their eukaryotic relatives. Top, the phylogenetic relationships among the species whose gene contents were compared. Each colored node represents the last common ancestor of a group of species. Bottom, a heat map of the 13,358 orthologous gene families inferred to have been present in at least one of six nodes representing common ancestors of interest: Ureukaryote, Uropisthokont, Urholozoan, Urchoanozoan, Urchoanoflagellate, and Urmetazoan (the full heat map for all gene families is shown in Supp. Figure 6). Each row represents a gene family. Gene families are sorted by their presence in each group of species, indicated by colored bars and boxes (eukaryotes, opisthokonts, holozoans, choanozoans, choanoflagellates and animals) and subsequently clustered within groups by uncentered Pearson correlation.

### Differential retention and loss of ancestral gene families in extant animals and choanoflagellates

While the phenomenon of gene loss has been well documented in the evolution of animals and other eukaryotes (Wolf and Koonin, 2013; Albalat and Cañestro, 2016; O’Malley et al., 2016), it has been unclear which extant animals come closest to representing the gene content of the Urmetazoan. Using the Urchoanozoan and Urmetazoan gene family catalogs reconstructed in this study, we ranked extant species based on their conservation of ancestral gene families (Figure 3, Supp. Figure 8). Compared with other animals in our study, the cephalochordate *Branchiostoma floridae* retains the most gene families that evolved along the animal stem lineage and also the most gene families with pre-choanozoan ancestry [extending prior observations that *B. floridae* preserved a comparatively large portion of the gene content of the last common ancestor of chordates (Louis et al., 2012)]. Among the non-bilaterian animal lineages, the cnidarian *Nematostella vectensis* most completely retains the Urmetazoan genetic toolkit [consistent with previous findings of conservation between *N. vectensis* and bilaterians (Putnam et al., 2007; Sullivan and Finnerty, 2007)], followed by the sponge *Oscarella pearsei*. Importantly, *B. floridae, N. vectensis*, and *O. pearsei* each retain different subsets of the Urmetazoan gene catalog, as only two thirds (67%) of the genes retained in at least one of the species are found in all three species. In contrast, the more rapidly evolving ecdysozoans *C. elegans, Pristionchus pacificus* and *Tetranychus urticae*, as well as the ctenophore *Mnemiopsis leidyi*, retain the fewest ancestral gene families, suggesting widespread gene family loss in these lineages, although the draft nature of some of their genome assemblies and high rates of sequence evolution may artificially inflate counts of missing genes.

**Figure 3.**
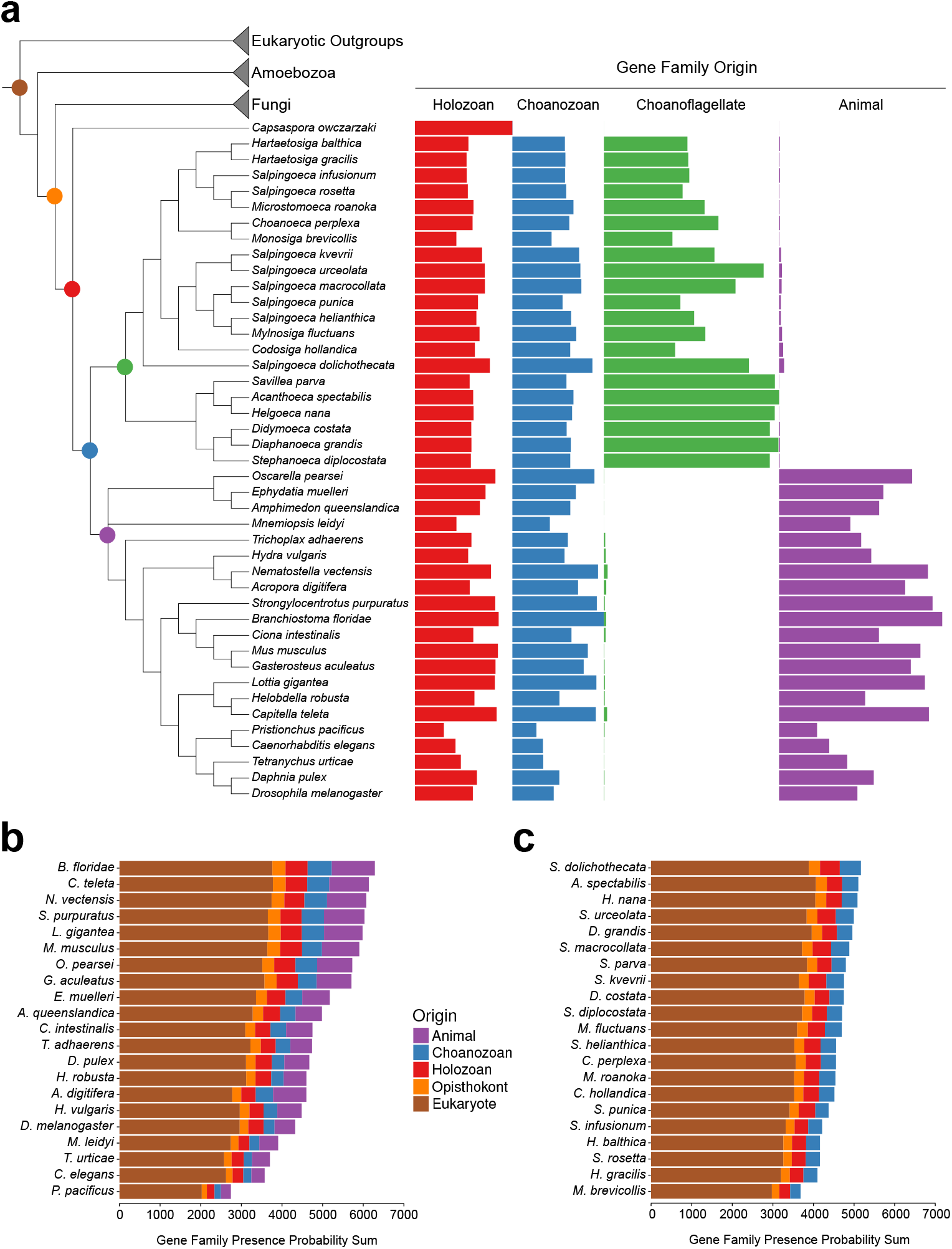
Gene family retention in animals, choanoflagellates, and *Capsaspora owczarzaki*. (a) Phylogenetic tree with gene family retention. Gene families are divided by their origin in the last common ancestor in different groups: holozoan, choanozoan, choanoflagellate, or animal. Colors correspond to nodes indicated in the phylogenetic tree. Bars represent the sum of presence probabilities for gene families with each origin. [Note that a small sum of probability is assigned to choanoflagellates for animal-specific gene families, and vice versa. This represents the sum of probabilities for the gene families that did not meet the 0.1 average presence probability threshold, but still had some residual probability between 0 and 0.1 (see Methods)]. (b-c) Ranked order of gene family retention for (b) animals and (c) choanoflagellates, similar to (a), but with the addition of gene families originating in the last common ancestor of Opisthokonts and of eukaryotes. Gene families originating within choanoflagellates are not included, in order to focus only on those potentially shared with animals.

Of the 21 choanoflagellates in our analysis, *Salpingoeca dolichothecata* [which is not closely related to *Salpingoeca rosetta* (Carr et al., 2017)] retains the most choanozoan-specific gene families, and therefore may be relatively more informative for comparative genomic studies of animal origins than are the two choanoflagellate species with previously-sequenced genomes, *M. brevicollis* and *S. rosetta*, which retain among the fewest ancestral gene families of any choanoflagellate (Figure 3, Supp. Figure 8). Several key gene families that were previously thought to be absent from choanoflagellates are conserved in *S. dolichothecata* and other choanoflagellates: the ancient ribonucleases Argonaute and Dicer, which are required for RNAi across eukaryotes (Jinek and Doudna, 2009), and holozoan gene families previously found in *C. owczarzaki* that are important for the regulation of animal development, including the transcription factors Churchill and Runx (Sebé-Pedrós et al., 2011) and a diagnostic domain for integrin β (Sebé-Pedrós et al., 2010) (Supp. Figure 9, Supp. Figure 10, Supp. Table 6, Methods). These findings echo the more general observation that the criteria used to select species for genome sequencing, often based on the ease with which they can be cultivated in the laboratory, also frequently select for those with streamlined genomes [e.g., (Gu et al., 2005)].

### Animal-specific gene families: innovation and loss

Gene families that originated on the stem lineage leading to animals are more likely to function in pathways or processes that distinguish animals from other eukaryotic groups. We identified ~1,718 such animal-specific gene families (Supp. Table 5), which were enriched for functions in cell communication, gene expression regulation, and cell adhesion (all three p < 10^−8^; Methods, Supp. Figure 11, Supp. Table 7), three processes that have previously been hypothesized to have been required for the evolution of animal multicellularity (King, 2004; Rokas, 2008; de Mendoza et al., 2013; Babonis and Martindale, 2017; Sebé-Pedrós et al., 2017). Animal-specific gene families include well-known developmental receptors, signaling proteins and transcription factors such as TGF-β, Hedgehog, Pax and Sox [consistent with previous reports (Srivastava et al., 2010; Riesgo et al., 2014)]. Notably, we detected many animal-specific gene families with no known function; 203 gene families (12% of total) lack any Pfam domain, and a further 50 (3%) are annotated only with Pfam domains of unknown function. The biochemical activities of these uncharacterized animal-specific gene families remain to be discovered, along with their roles in animal development and evolution.

Of the ~1,718 gene families that originated on the animal stem lineage, we found only 36 that were subsequently retained in all 21 animal genomes we analyzed (Table 1), leaving ~1,682 gene families that, despite being specific to animals, were lost in one or more extant lineages. We thus asked whether these 36 core animal-specific gene families might participate in pathways critical for animal biology. Indeed, this set of genes includes seven from the Wnt pathway (including Frizzled, Dishevelled, TCF/LEF and β-catenin), five involved in cell-cell adhesion (including integrin α, laminin, and vinculin), and other well-known animal gene families such as JNK, caspases, and metabotropic glutamate receptors. [Recent studies in myxozoans, a parasitic lineage of cnidarians, and in glass sponges, which develop into syncytial larvae and adults, revealed the absence of some Wnt components, indicating that even among the 36 core animal-specific genes, those involved in developmental patterning appear to be dispensable in animals with dramatically derived body plans (Chang et al., 2015; Schenkelaars et al., 2017).] The 36 core animal gene families also include several that are less well characterized or whose specific contributions to animal origins and animal biology are not immediately obvious, such as two subunits of the transcription-regulating Mediator complex (Malik and Roeder, 2010) and the ubiquitination-associated BTB/POZ domain-containing protein 9 (DeAndrade et al., 2012). For comparison, choanoflagellates have a similarly small set of 57 gene families (out of ~2,005 choanoflagellate-specific gene families) that are conserved in all 21 choanoflagellate species that we sampled; 30% of these gene families encode Pfam domains related to kinase signaling (including protein kinases, phosphatases and adapters; Supp. Table 8).

**Table 1.** Core animal-specific gene families that are present in all animals in this study. Representative gene names and annotations are based on a consensus from each gene family (Methods). Gene families are ordered by pathway/function. *: missing in myxozoans, a lineage of parasitic cnidarians (Chang et al., 2015), ** missing in myxozoans and in glass sponges (Schenkelaars et al., 2017), two animal lineages with dramatically derived body plans.

| **Table 1. Core animal-specific gene families that are present in all animals in this study.** |  |  |
| --- | --- | --- |
| Representative gene names and annotations are based on a consensus from each gene family (Methods). Gene families are ordered by pathway/function. * : missing in myxozoans, a lineage of parasitic cnidarians (Chang et al., 2015), ** missing in myxozoans and in glass sponges (Schenkelaars et al., 2017), two animal lineages with dramatically derived body plans. |  |  |
| **Gene Family ID** | **Representative Gene Name(s)** | **Pathway/Function** |
| 3849 | T-box transcription factor TBX 2/3 | gene regulation |
| 5290 | mediator subunit | gene regulation |
| 6241 | mediator subunit | gene regulation |
| 6675 | nuclear factor 1 A/B | gene regulation |
| 8693 | interleukin enhancer-binding factor 2 | gene regulation |
| 5720 | lethal(2) giant larvae | gene regulation |
| 7805 | MEX3 B/C | gene regulation |
| 441 | low-density lipoprotein receptor (LRP) 1/2/4/5/6 | Wnt* |
| 3570 | frizzled 1/2/5/7/8 | Wnt* |
| 4891 | catenin delta | Wnt |
| 5637 | transcription factor 7 (TCF/LEF) | Wnt |
| 6000 | transcription factor COE 1-4 | Wnt |
| 6254 | dishevelled 1-3 | Wnt** |
| 6532 | catenin beta | Wnt |
| 804 | hemicentin/obscurin/titin | cell-cell adhesion |
| 4442 | integrin alpha 2/5/8 | cell-cell adhesion |
| 6831 | fermitin 1-3 | cell-cell adhesion |
| 7164 | laminin gamma 1-3 | cell-cell adhesion |
| 8024 | vinculin | cell-cell adhesion |
| 476 | metabotropic glutamate receptor 1-8 | synapse / vesicle |
| 4993 | calcium-dependent secretion activator | synapse / vesicle |
| 7929 | receptor-type tyrosine-protein phosphatase-like N | synapse / vesicle |
| 6010 | MAPK 8-10 | MAPK / JNK |
| 8406 | kinase suppressor of Ras 2 | MAPK / JNK |
| 2737 | caspase 3/7/9 | apoptosis |
| 5675 | calumenin | calcium ion |
| 6842 | cyclin T 1/2 | cell cycle |
| 621 | dystonin/desmoplakin/plectin | cytoskeleton |
| 7916 | phosphorylase b kinase | glycan |
| 6512 | heparan-sulfate 6-O-sulfotransferase 1-3 | heparan sulfate |
| 7146 | inositol monophosphatase 3 | inositol |
| 4051 | tetraspanin 5/14/17/33 | metalloprotease |
| 6163 | protein kinase C iota/zeta | PI3K |
| 6251 | small G protein signaling modulator 1-2 | Rab GTP |
| 6366 | MAP kinase-activating death domain protein 9-11 | TNF |
| 8587 | BTB/POZ domain-containing protein 9 | ubiquitin |

While novel features of animal biology might have evolved with the emergence of new gene families, the loss of ancient genes also influenced animal origins. We detected ~1,645 gene families that evolved prior to the choanoflagellate-animal divergence that were retained in choanoflagellates and lost entirely from animals. These include gene families in pathways necessary for the biosynthesis of the amino acids leucine, isoleucine, valine, methionine, histidine, lysine and threonine (Supp. Figure 12, Supp. Table 9). The shikimic acid pathway, which is required for the synthesis of the aromatic amino acids tryptophan and phenylalanine, and other aromatic compounds, was also lost along the animal stem lineage [although subsequently regained in cnidarians through horizontal gene transfer from bacteria (Fitzgerald and Szmant, 1997; Starcevic et al., 2008)]. We thus demonstrate that components of the biosynthesis pathways for nine amino acids that are essential in animals (Payne and Loomis, 2006; Guedes et al., 2011) were lost on the animal stem lineage, and not prior to the divergence of choanoflagellates and animals. The SLN1 two-component osmosensing system, which has been shown in fungi to regulate acclimation to changes in environmental salinity (Posas et al., 1996), is also conserved in choanoflagellates but absent in animals. Together, the ensemble of animal gene family losses reflects the substantial changes in metabolism and ecology that likely occurred during early animal evolution.

### Choanozoan-specific gene families: innovation and loss

In addition to the set of gene families that evolved on the animal stem lineage, those that originated on the holozoan and choanozoan stem lineages also contributed to the genomic heritage of animals. Our increased sampling of choanoflagellate diversity allowed us to ask whether gene families previously thought to have been animal innovations, due to their absence from *M. brevicollis*, *S. rosetta* and other outgroups, may in fact have evolved before the divergence of animals and choanoflagellates. We found that ~372 gene families previously thought to be restricted to animals have homologs in one or more of the 19 newly sequenced choanoflagellates (Supp. Table 5; see Supp. Table 10 for a list of pathways with components gained or lost on the Choanozoan stem lineage).

Within this set of genes are the Notch receptor and its ligand Delta/Serrate/Jagged (hereafter Delta), which interact to regulate proliferation, differentiation and apoptosis during animal developmental patterning (Artavanis-Tsakonas, 1999). Intact homologs of these important signaling proteins have never previously been detected in non-animals, although some of their constituent protein domains were previously found in *M. brevicollis, S. rosetta* and *C. owczarzaki* (King et al., 2008; Suga et al., 2013). In our expanded choanoflagellate data set, we detected a clear Notch homolog in *Mylnosiga fluctuans* with the prototypical EGF (epidermal growth factor), Notch, transmembrane and Ank (ankyrin) domains in the canonical order, while five other choanoflagellates contain a subset of the typical protein domains of Notch proteins (Supp. Figure 10, Supp. Figure 13). Similarly, the choanoflagellate species S. *dolichothecata* expresses a protein containing both of the diagnostic domains of animal Delta (MNNL and DSL, both of which were previously thought to be animal-specific) and four other choanoflagellate species contain one of the two diagnostic domains of Delta, but not both. Moreover, the Notch/Delta downstream transcriptional regulator Suppressor of Hairless is present in all choanoflagellate species analyzed, and therefore was likely present in the Urchoanozoan. The distributions of Notch and Delta in choanoflagellates suggest that they were present in the Urchoanozoan and subsequently lost from most (but not all) choanoflagellates, although it is formally possible that they evolved convergently through shuffling of the same protein domains in animals and in choanoflagellates. A similar portrait emerges for the cadherins Flamingo and Protocadherin (Chae et al., 1999; Usui et al., 1999; Frank and Kemler, 2002) that were previously thought to be animal-specific, but are found in a subset of choanoflagellates within our data set (Supp. Figure 10, Methods).

We also found evidence that numerous gene families and pathways that originated in animals arose through shuffling of more ancient protein domains and genes that were already present in the Urchoanozoan (Ekman et al., 2007; King et al., 2008; Grau-Bové et al., 2017). For example, the new choanoflagellate gene catalogs confirm that several signature animal signaling pathways, such as Hedgehog, Wnt, JNK, JAK-STAT, and Hippo, are composed of a mixture of gene families that were present in the Urchoanozoan and others that evolved later on the animal stem lineage or within animals (Snell et al., 2006; Adamska et al., 2007; Hausmann et al., 2009; Richards and Degnan, 2009; Srivastava et al., 2010; Sebé-Pedrós et al., 2012; Babonis and Martindale, 2017) (Supp. Table 9). For another animal signaling pathway, TGF-β, the critical ligand, receptor and transcription factor gene families are composed of animal-specific domain architectures (Heldin et al., 1997; Munger et al., 1997), although all three contain constituent protein domains that evolved on the choanozoan stem lineage (Supp. Figure 14).

### The pre-animal origins of the animal innate immunity pathway

One surprise from our analyses was the existence in choanoflagellates of genes required for innate immunity in animals. Although innate immunity is a feature of both animal and plant biology, the receptors and pathways used by these evolutionarily distant organisms are thought to have evolved independently (Ausubel, 2005). The animal immune response is initiated, in part, when potential pathogens stimulate (either directly or indirectly) the Toll-like receptors (TLRs), which have previously been detected only in animals (Leulier and Lemaitre, 2008). Importantly, although TLRs are found in nearly all bilaterians and cnidarians, they are absent from placozoans, ctenophores, and sponges [proteins with similar, but incomplete domain architectures have been detected in sponges (Miller et al., 2007; Riesgo et al., 2014)] and were therefore thought to have evolved after the origin of animals.

We found that 14 of 21 sequenced choanoflagellates encode clear homologs of animal TLRs (Figure 4a-c), implying that TLRs first evolved on the Urchoanozoan stem lineage (Methods). Animal and choanoflagellate TLRs are composed of an N-terminal signal peptide, multiple leucine rich repeat (LRR) domains, a transmembrane domain, and an intracellular Toll/interleukin 1 receptor/resistance (TIR) domain. In the canonical TLR signaling pathway, the interaction of the intracellular TIR domain of Toll-like receptors with TIR domains on adapter proteins (e.g., MyD88) initiates one of a number of potential downstream signaling cascades that ultimately lead to activation of the NF-*κ*B transcription factor (Janeway and Medzhitov, 2002).

**Figure 4.**
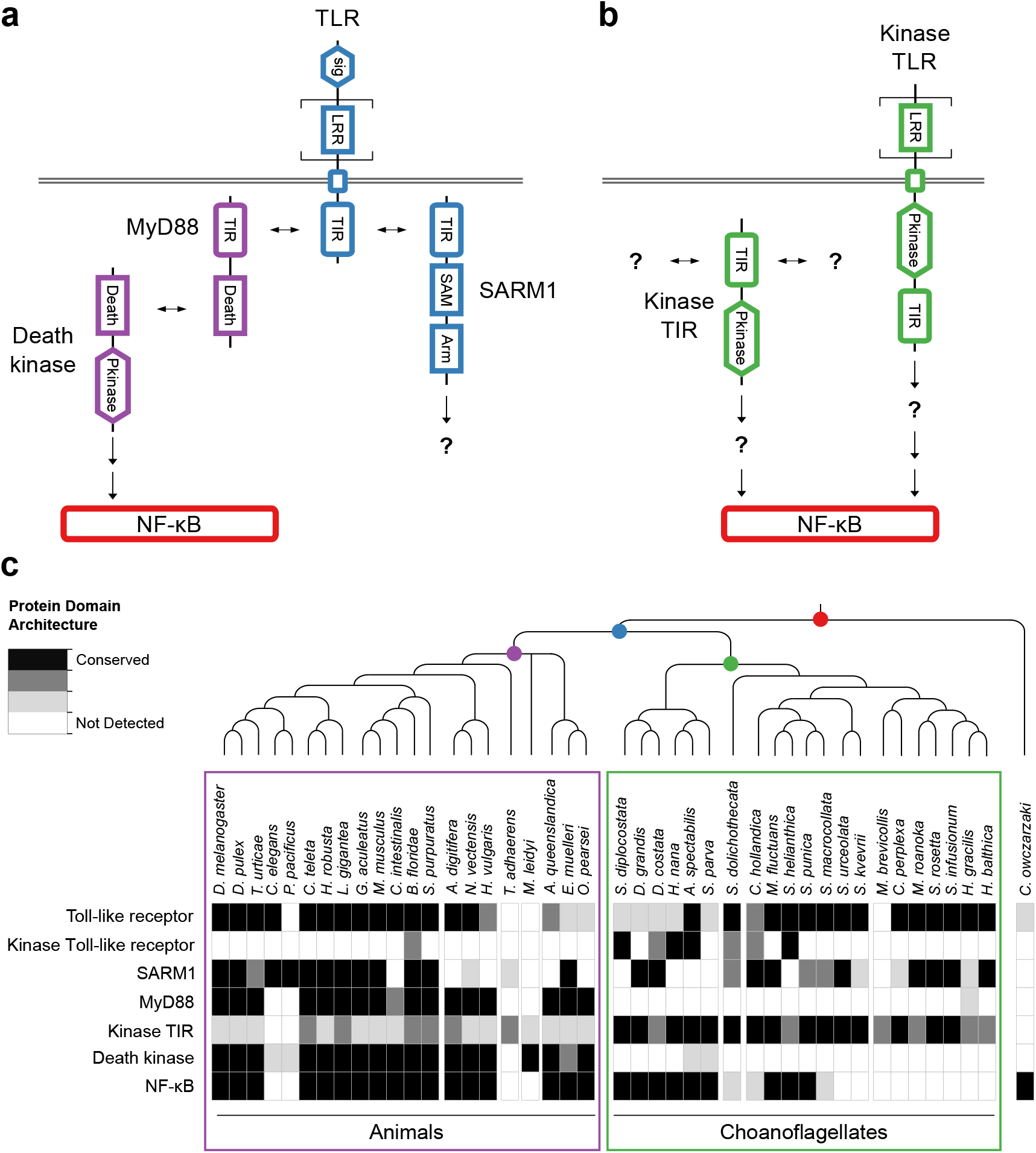
Evolution of the TLR signaling pathway. Components of the canonical TLR pathway (a) and potential choanoflagellate Kinase TLR signaling (b), with their canonical domain architectures and colored by their inferred ancestral origin (blue = choanozoan ancestry, purple = metazoan ancestry, green = choanoflagellate ancestry, and red = holozoan ancestry). Question marks denote steps of the signaling pathway and/or interaction partners that are hypothesized, but untested. (c) Presence of receptors, adapters, kinases and the transcription factor NF-*x*B in animals, choanoflagellates and *C. owczarzaki*.

To investigate whether the Urchoanozoan TLR might have activated a downstream signaling pathway that resembled the canonical TLR pathway in animals, we searched for TLR adapters, downstream kinases and NF-*κ*B in choanoflagellates (Figure 4c, Supp. Table 6). While many choanoflagellates encode NF-*x*B, we found no evidence for two critical Death domain-containing proteins involved in the TLR-dependent activation of NF-*κ*B: the adapter protein MyD88 (Wiens et al., 2005; Gauthier et al., 2010) and the downstream kinase IRAK (Song et al., 2012). However, we did detect two new classes of choanoflagellate-specific proteins that pair kinase domains directly with LRR and/or TIR domains, seemingly bypassing the need to recruit kinases in multi-protein signaling complexes: TLR-like proteins with an intracellular kinase domain positioned between the transmembrane domain and TIR domain (which we provisionally term “Kinase TLRs”) and proteins encoding TIR and kinase domains, but lacking a transmembrane domain (which we provisionally term “Kinase TIRs”). In addition, we detected homologs of the TIR-containing adapter SARM1, a multi-functional protein that can trigger both NF-*x*B-dependent and NF-*κ*B-independent responses (Couillault et al., 2004; Sethman and Hawiger, 2013; Liu et al., 2014). Choanoflagellate SARM1 homologs contain a conserved glutamic acid residue that is necessary for SARM1 NADase activity in animals (Essuman et al., 2017) (Supp. Figure 15). Finally, although we did not detect most animal cytosolic innate immune sensors in choanoflagellates, including the LRR-containing NLR family, ALRs, MAVS, MDA5 and RIG-I, we did find evidence for both cGAS and STING in choanoflagellates [as previously reported in *M. brevicollis* (Wu et al., 2014)] (Methods). Thus, critical components of the animal innate immune pathway, including both extracellular and intracellular pattern sensing receptors, predate animal origins.

### Chondroitin sulfate cleavage promotes rosette colony growth in S. helianthica

Glycosaminoglycans (GAGs) are specialized carbohydrate polymers that form an important part of the animal extracellular matrix (ECM), the scaffold among neighboring cells that is involved in nearly every aspect of animal biology (Hay, 1991; Adams and Watt, 1993; Sorokin, 2010). The ability to produce GAGs was until recently thought to be an innovation of animals. The biosynthetic pathway for the GAG chondroitin sulfate is found across animal diversity, including in early-branching lineages such as sponges, and was recently detected in *S. rosetta*, extending the evolutionary history of GAGs to the stem-choanozoan era (Woznica et al., 2017). However, relatively little is known about the evolutionary history of enzymes that cleave GAGs and thereby contribute to the regulation of animal ECM.

There are two major classes of GAG-cleaving enzymes that, based on their phylogenetic distributions and mechanisms of action, likely evolved independently (Stern and Jedrzejas, 2006). The GAG hydrolases, which promote, for example, ECM cleavage during sperm/egg fusion and venom-induced ECM digestion (Cherr et al., 2001; Girish and Kemparaju, 2007), were previously thought to be animal-specific. GAG lyases, on the other hand, have previously been found almost exclusively in bacteria and a small number of fungi (Hynes and Walton, 2000), where they likely allow these organisms to digest animal (and potentially choanoflagellate) ECM, either as a food source or to facilitate pathogenesis and other eukaryote-microbe interactions.

We detected homologs of animal GAG hydrolases in at least 12 of the 21 choanoflagellate species included in this study (Figure 5a, Supp. Figure 16). Therefore, GAG hydrolases are not animal-specific, but instead evolved earlier, in a similar time frame to the chondroitin sulfate biosynthetic pathway. Interestingly, at least 18 of the 21 choanoflagellates in this study also encode a GAG lyase, which in animals are only found in sponges and cnidarians, suggesting that a GAG lyase was present early in eukaryotic evolution, conserved in the Urchoanozoan and Urmetazoan, and subsequently lost from most animals.

**Figure 5.**
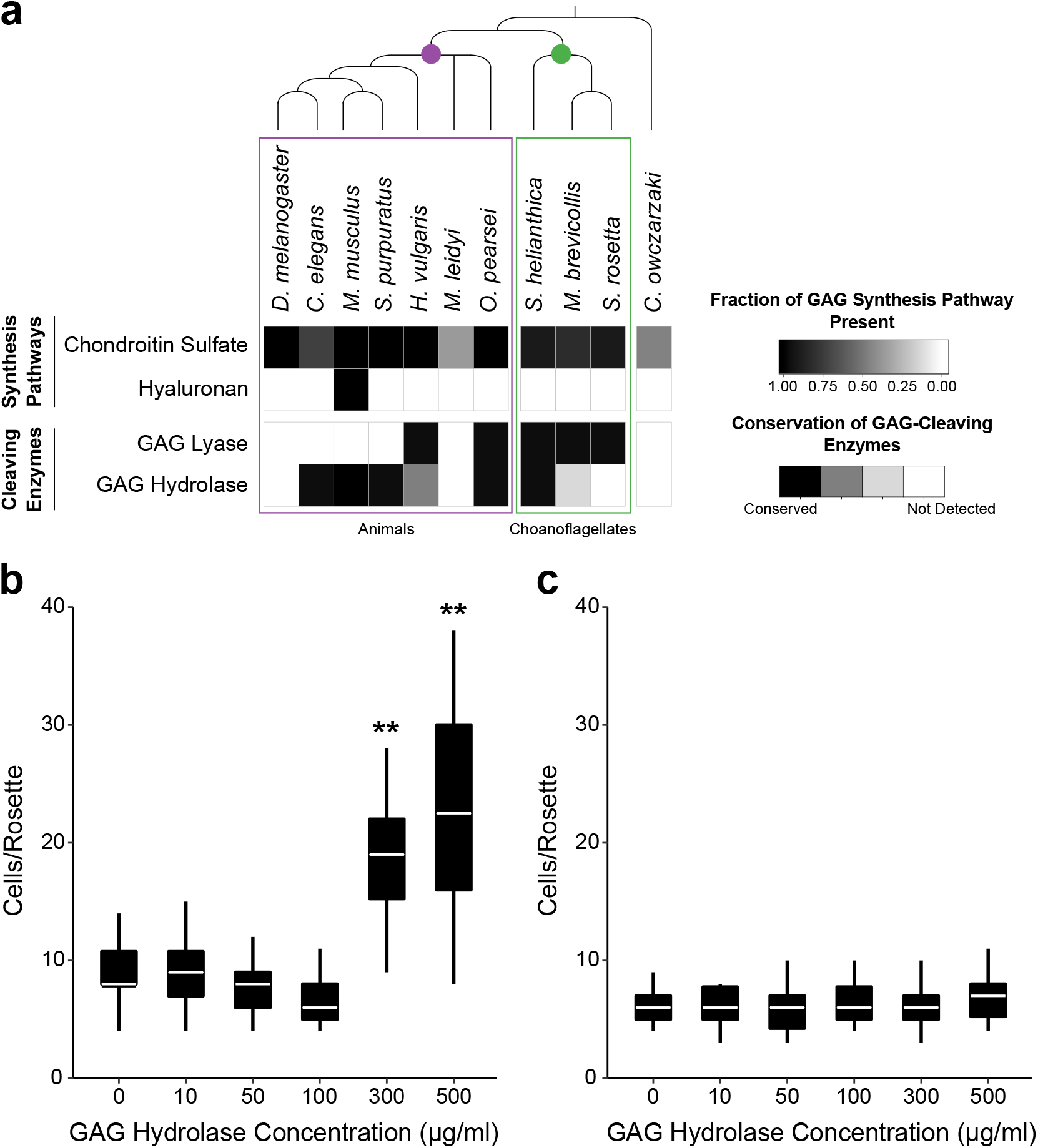
Choanoflagellates encode the genes for glycosaminoglycan (GAG) synthesis and cleavage, and treatment with a GAG hydrolase promotes rosette colony growth in the choanoflagellate *Salpingoeca helianthica*. (a) Presence/absence of GAG biosynthesis genes [adapted from (Woznica et al., 2017)] and GAG-cleaving enzymes. *S. helianthica* expresses a homolog of metazoan GAG hydrolase, whereas *Salpingoeca rosetta*, another colony-forming choanoflagellate, does not. (b) The number of cells per rosette in *S. helianthica* increases following treatment with bovine GAG hydrolase. (c) GAG hydrolase has no detectable effect on the number of cells per rosette in *S. rosetta*. ** p < 10^−8^ different from untreated control, Wilcoxon rank sum test. N = 30 colonies per treatment. The white bar represents the median, the boxes represent the 25th to 75th percentiles, and the whiskers extend to the largest and smallest numbers of cells per colony (but no further than 1.5 times the distance between the 25th and 75th percentiles).

Both *S. rosetta* and another rosette colony-forming choanoflagellate, *S. helianthica*, produce an ECM visible by electron microscopy [Supp. Figure 17a-b; (Dayel et al., 2011)] and express the genes necessary to synthesize chondroitin sulfate, but lack the genes to synthesize other GAGs such as hyaluronan (Figure 5a, Supp. Table 11). *S. helianthica* expresses both a GAG hydrolase and a GAG lyase, whereas *S. rosetta* encodes only GAG lyases, raising the possibility that they use different mechanisms for regulating chondroitin sulfate-bearing proteoglycans in their ECM.

Because ECM is necessary for multicellularity in animals (Hay, 1991; Adams and Watt, 1993; Abedin and King, 2010) and has been implicated in choanoflagellate multicellularity (Levin et al., 2014) we tested how treatment of *S. helianthica* and *S. rosetta* with GAG hydrolase would influence rosette morphology. Treatment of *S. helianthica* with 500 *μ*g/ml of a well-characterized commercially available bovine GAG hydrolase significantly increased the number of cells per rosette from ~9 cells/rosette to ~23 cells/rosette (Figure 5b, Supp. Figure 17). In contrast, *S. rosetta* rosette size showed no response to bovine GAG hydrolase (Figure 5c, Supp. Figure 17). We infer that animal GAG hydrolases influence *S. helianthica* colony biology by cleaving chondroitin sulfate molecules in the ECM. Given that *S. rosetta* produces chondroitin sulfate, but does not produce an endogenous GAG hydrolase, its insensitivity to a heterologous GAG hydrolase hints at the potential specificity of chondroitin sulfate cleavage in rosette morphogenesis.

## Discussion

Our increased sampling of choanoflagellates reveals how Urmetazoan gene content evolved as a mosaic of old, new, rearranged, and repurposed protein domains, genes and pathways. We have identified roughly 8,200 gene families that were present in the Urmetazoan [consistent with recent findings (Simakov and Kawashima, 2017)], about 80% of which were also present in the Urchoanozoan and the remainder of which evolved on the animal stem lineage (Supp. Table 5). The patchwork ancestry of the Urmetazoan genome is illustrated by the fact that many gene families responsible for animal development, immunity and multicellular organization evolved through shuffling of protein domains that first originated in the choanozoan stem lineage together with ancient or animal-specific domains (e.g. the TGF-β ligand and receptor; Supp. Figure 10, Supp. Figure 14). In addition, other gene families found in the Urchoanozoan were subsequently combined into new pathways in the animal stem lineage along with newly evolved genes (e.g., the TLR and Hedgehog pathways; Figure 4, Supp. Table 9). On the other hand, the history of the Urmetazoan genome is not simply one of innovation and co-option, as roughly 1,650 Urchoanozoan genes were lost, including genes for the synthesis of nine essential amino acids (Supp. Figure 12, Supp. Table 9) (Payne and Loomis, 2006; Guedes et al., 2011; Erives and Fassler, 2015).

One of the most critical events for animal origins was the transition to multicellularity. Multicellularity evolved repeatedly across eukaryotic diversity, with animals, plants, and fungi each representing the results of independent transitions to multicellularity (Buss, 1987; Bonner, 1998; King, 2004). While it has long been thought that the special case of animal multicellularity evolved after the divergence of the animal and choanoflagellate stem lineages, there are some hints that the Urchoanozoan might have been capable of multicellular development (Carr et al., 2008a, 2017, Fairclough et al., 2010, 2013). Many choanoflagellate species form multicellular colonies (Leadbeater, 2015; Carr et al., 2017). This occurs in *S. rosetta*, and likely other choanoflagellates, through a developmental process akin to zygotic cleavage in animals (Fairclough et al., 2010; Dayel et al., 2011). Multicellular integrity in animals is stabilized by a shared ECM that contains proteoglycans decorated with the GAG chondroitin sulfate (Hay, 1991); similarly, choanoflagellates synthesize chondroitin sulfate (Woznica et al., 2017) and their multicellular rosettes may be stabilized via the ECM (Levin et al., 2014). In animals, ECM remodeling, including through the cleavage of GAG proteoglycans, is required for proper development and regeneration following injury (Busch and Silver, 2007; Giger et al., 2010; Alilain et al., 2011). We show here that GAG hydrolases used by animals to remodel their proteoglycan-rich ECM are also produced by diverse choanoflagellates. Although the functions of endogenous GAG hydrolases in choanoflagellates have yet to be investigated, treatment of the choanoflagellate *S. helianthica* with an animal GAG hydrolase leads to a significant increase in rosette colony size. It is possible that the *S. helianthica* GAG hydrolase regulates ECM availability, and thus rosette size, by modulating ECM structure, the dysregulation of which has been implicated in pathogenesis in animals (Bonnans et al., 2014). In contrast, in the choanoflagellate *S. rosetta*, which does not encode an endogenous GAG hydrolase but does possess a GAG lyase, animal GAG hydrolases have no effect on rosette size, although bacterially-produced GAG lyases have been shown to induce sexual reproduction in *S. rosetta* (Woznica et al., 2017).

The origin of multicellularity in animals provided novel niches for bacteria to exploit, requiring the evolution of new mechanisms for mediating interactions with pathogenic and commensal bacteria. Nonetheless, the roots of animal innate immunity clearly predate animal origins. We have found that choanoflagellates express TLRs, transmembrane receptors that trigger the animal innate immune response, as well as its canonical downstream signaling target, NF-*κ*B, suggesting that both existed in the in the Urchoanozoan. Like modern choanoflagellates and sponges, the Urchoanozoan likely preyed upon bacteria (McFall-Ngai et al., 2013; Richter and King, 2013), and bacterial cues can induce life history transitions in choanoflagellates (Alegado et al., 2012; Woznica et al., 2016, 2017), although the mechanisms by which choanoflagellates capture bacteria and sense bacterial cues are unknown. We hypothesize that the core TLR/NF-*κ*B pathway functioned in prey sensing, immunity, or more complex processes in the Urchoanozoan that subsequently formed the basis of a self-defense system in animals. Because critical pathway members linking TLR and NF-*κ*B appear to be animal innovations (e.g., MyD88), the animal signaling pathway may have evolved to diversify downstream signaling processes to tailor responses in a multicellular context. This pathway diversification may have included the evolution of roles in development, as TLRs have been implicated in both NF-*κ*B-dependent and NF-*x*B-independent developmental signaling (in addition to their functions in immunity) in bilaterians and in the cnidarian *N. vectensis* (Leulier and Lemaitre, 2008; Brennan et al., 2017). The uncharacterized choanoflagellate-specific Kinase TLRs and Kinase TIRs may function as part of a streamlined signaling pathway that mediates responses to extracellular cues, including bacteria, although further research will be required to test this hypothesis.

Our study provides an exhaustive and detailed view of the changes in gene content that laid the foundation for animal origins. Changes in how Urmetazoan gene families were regulated also likely played a role in animal evolution, as aspects of animal phosphoproteome remodeling are shared with *C. owczarzaki* (Sebé-Pedrós et al., 2016b), and features of animal gene co-regulation and alternative splicing (but not animal promoter types and enhancers) have been found in *C. owczarzaki* (Sebé-Pedrós et al., 2013b, 2016a) and in the holozoan *Creolimax fragrantissima* (de Mendoza et al., 2015). In comparison, within other lineages that independently evolved multicellularity based on distinct genomic foundations, other types of changes in gene family content gave rise to different multicellular forms. In the social amoeba *Dictyostelium discoideum*, many novel gene families are involved in extracellular sensing (Glöckner et al., 2016), similar to the marked increase in signal transduction gene families found in the multicellular brown alga *Ectocarpus siliculosus* (Cock et al., 2010), whereas gene innovations in multicellular ascomycete fungi are enriched for functions related to endomembrane organelles (Nguyen et al., 2017). In contrast, the gene complement of the multicellular green alga *Volvox carteri* is largely distinguished from its unicellular relative *Chlamydomonas reinhardtii* by expansions or contractions within gene families, rather than the evolution of new families (Prochnik et al., 2010). Our work identifies the specific gene families whose invention or co-option distinguished the Urmetazoan from all other organisms and therefore contributed to the birth of unique mechanisms for regulating development, homeostasis and immunity in animals.

## Materials and Methods

The sections on Quality trimming, Error correction, *De novo* transcriptome assembly, Identification and removal of cross-contamination, Prediction of amino acid sequences from assembled transcripts and elimination of redundant transcripts, and Measurement of expression levels and elimination of noise transcripts were described in (Peña et al., 2016). They are repeated here for convenience and clarity (with modifications to the text but not to the underlying methods).

### Origin of cultures

We acquired 18 of 20 cultures used for transcriptome sequencing from external sources (Supp. Table 1). We isolated the remaining two cultures, *Acanthoeca spectabilis* (Virginia) ATCC PRA-387 and *Codosiga hollandica* ATCC PRA-388. (A. *spectabilis* (Virginia) is a different isolate from A. *spectabilis* ATCC PRA-103, which was originally collected in Australia.) We collected the water sample from which *A. spectabilis* (Virginia) was isolated on December 19, 2007 near Hog Island, Virginia (GPS coordinates: 37.472502, −75.816018) and we isolated *C. hollandica* from a sample collected on June 25, 2008 from Madeira, Portugal (GPS coordinates: 32.762222, −17.125833). C. *hollandica* was formally described in Carr *et al.*, 2017 (Carr et al., 2017).

We isolated choanoflagellates with a micromanipulator and a manual microinjector (PatchMan NP 2 and CellTram vario 5176 (Eppendorf, Hamburg, Germany) for A. *spectabilis*, and XenoWorks Micromanipulator and Analog Microinjector (Sutter Instrument, Novato, California, United States) for *C. hollandica)*. We pulled glass needles used for isolation from 1 mm diameter borosilicate glass (GB100-10, Science Products GmbH, Hofheim, Germany) using a Flaming/Brown needle puller (P-87, Sutter Instrument) with the following program: heat = 820, pull = 50, velocity = 140, time = 44. We polished and sterilized needles by passing them briefly through a low flame. We used a separate needle for each attempted isolation, transferring single cells into separate culture flasks containing appropriate growth medium (see Supp. Table 1). In order to reduce the possibility of contamination during the isolation procedure, we generated sterile air flow across the microscope and micromanipulator apparatus using a HEPA-type air purifier (HAP412BN, Holmes, Boca Raton, Florida, United States).

One culture we obtained from ATCC, *Salpingoeca infusionum*, was contaminated by an unidentified biflagellated unicellular eukaryote. To remove the contaminant from the culture, we counted then diluted cells into separate wells of two 24-well plates. After 7 days of growth, we found 4 of 48 wells containing only *S. infusionum* and bacteria, 1 well containing only the contaminant and bacteria, and the remaining 43 wells containing only bacteria. We selected one of the four wells containing only *S. infusionum* for use in transcriptome sequencing.

### Antibiotic treatment and optimization of culture conditions

Choanoflagellates are co-isolated with diverse bacterial prey, which serve as a food source. In order to limit bacterial diversity to the species that led to optimal choanoflagellate growth, we treated each culture with a panel of ten different antibiotics (Supp. Table 12). We obtained all antibiotics from Thermo Fisher Scientific (Waltham, Massachusetts, United States) with the exception of erythromycin, gentamicin, ofloxacin, and polymyxin B (Sigma-Aldrich, St. Louis, Missouri, United States). We sterilized antibiotic solutions before use by filtration through a 0.22 *μ*m syringe filter (Thermo Fisher Scientific) in a laminar flow hood. We initially treated each culture with all ten antibiotics. We selected initial treatments that decreased bacterial density and diversity, and then repeatedly diluted the cultures into fresh medium with the same antibiotic until no further change in bacterial density or diversity was observed. We then re-treated each of these cultures with an additional test of all ten antibiotics, as their modified bacterial communities often responded differently from their initial communities. We repeated successive rounds of treatment until no further reduction in bacterial density or diversity was observed (Supp. Table 1).

We tested a range of concentrations of different temperatures and growth media (Supp. Table 13) in order to maximize choanoflagellate cell density, with three types of nutrient sources: quinoa grains, proteose peptone/yeast extract, and cereal grass. We used filtered water (Milli-Q, Millipore, Burlington, Massachusetts) when preparing all solutions. For marine species, we used 32.9 grams per liter of artificial seawater (Tropic Marin, Montague, Massachusetts, United States). We used proteose peptone (Sigma-Aldrich Chemical) at a final concentration of 0.002% (w/v) and yeast extract (Becton Dickinson Biosciences, San Jose, California, United States) at a final concentration of 0.0004% (w/v). To prepare cereal grass media (also known as chlorophyll alfalfa, Basic Science Supplies, St. Augustine, Florida, United States), we added it to autoclaved water and allowed it to steep until cool. Once cool, we removed large particles of cereal grass by repeated vacuum filtration through a ceramic Buchner funnel with a double layer of Grade 1 cellulose filter paper (Whatman, GE Healthcare Life Sciences, Marlborough, Massachusetts, United States). We autoclaved organic quinoa grains and added them to the medium after filtration, with roughly 2 grains added per 25 cm^2^ of culture vessel surface area. We measured final nutrient content of each type of medium by Flow Injection Analysis at the University of California, Santa Barbara Marine Science Institute (Supp. Table 3). We tested buffered medium for two freshwater species that experienced lowered pH during growth, *C. hollandica* (pH 5.5) and *Salpingoeca punica* (pH 5), using 50 mM HEPES (Thermo Fisher Scientific) adjusted to a pH of 7. We sterilized all media with a 0.22 *μ*m vacuum filter (Steritop, Millipore) in a laminar flow hood prior to use.

We selected final culture conditions that maximized choanoflagellate density and variety of cell types present, as we hypothesized that different cell types, each potentially expressing different subsets of genes, would yield the greatest diversity of transcripts for sequencing. We defined five generic choanoflagellate cell types: ‘attached’: attached to the bottom of the culture vessel or to a piece of floating debris, either directly, within a theca, within a lorica, or on a stalk; ‘slow swimmer’: a typical swimming cell; ‘fast swimmer’: a cell with reduced collar length swimming at higher speed; ‘free-swimming colonial’: in a colony swimming in the water column; ‘attached colonial’: in a colony attached to a stalk; ‘passively suspended’: suspended in the water column, either naked or within a lorica. See (Dayel et al., 2011; Carr et al., 2017) for further information on choanoflagellate cell types and (Leadbeater, 2015; Richter and Nitsche, 2016) for descriptions of extracellular structures (thecae, loricae, etc.).

### Growth of cultures in large batches in preparation for RNA isolation

We routinely grew choanoflagellates in 25 cm^2^ angled neck cell culture flasks with 0.2 *μ*m vented caps (Corning Life Sciences, Corning, New York, United States) containing 25 ml of medium. We performed all cell culture work in a laminar flow hood. To reduce the possibility of cross-contamination among samples in the hood, we dispensed media into growth vessels prior to the introduction of cultures, we only worked with a single culture at a time, and we cleaned thoroughly with 70% ethanol before and after introducing cultures. To grow and collect large amounts of cells for RNA preparation, we used different growth vessels, volumes, growth durations and centrifugation times as appropriate to each culture (Supp. Table 1). Vessels included long neck Pyrex glass culture flasks (Corning), 150 mm plastic tissue culture dishes (Becton Dickinson), and 75 cm^2^ angled neck cell culture flasks with 0.2 *μ*m vented caps (Corning).

We harvested cultures depending on the cell types present (Supp. Table 1). For cultures with 5 percent or fewer attached cells, we collected liquid by pipetting without scraping (to reduce the number of bacteria collected). For cultures containing between 5 and 90 percent attached cells, we harvested single plates by pipetting after scraping cells from the bottom of the plate. For cultures with 90 percent or greater attached cells, we combined multiple plates as follows: we removed and discarded the liquid from the first plate by pipetting, added 50 ml of either artificial sea water or filtered water, as appropriate, scraped cells from the plate, removed the liquid from the second plate, transferred the liquid from the first to the second plate, and repeated the procedure on subsequent plates. For cultures containing quinoa grains or large bacterial debris, we filtered with a 40 *μ*m cell strainer (Fisher) after collection.

We pelleted cells in 50 ml conical tubes at 3220 x g in a refrigerated centrifuge at 4° C, removed the first 47.5 ml of supernatant by pipetting, and the last 2.5 ml by aspiration. When we harvested more than 50 ml for a culture, we spun in separate tubes, removed all but 2.5 ml of supernatant, resuspended, combined into a single 50 ml conical tube, and repeated the centrifugation as above. We flash froze pellets in liquid nitrogen and stored them at −80° C. To reduce the possibility of cross-contamination, we harvested and centrifuged each culture separately, we used disposable plastic pipette tubes, conical tubes, and cell scrapers, and we cleaned all other material (bench tops, pipettes, etc.) with ELIMINase (Decon Laboratories, King of Prussia, Pennsylvania, United States) between cultures.

### RNA isolation

We isolated total RNA from cell pellets with the RNAqueous kit (Ambion, Thermo Fisher Scientific). We modified the manufacturer’s protocol to double the amount of lysis buffer, in order to increase RNA yield and decrease degradation. We performed both optional steps after adding lysis buffer: we spun for 3 minutes at top speed at 1° C and passed the supernatant through a 25 gauge syringe needle several times. We used the minimum suggested volumes in the two elution steps (40 *μ*l and 10 *μ*l). We measured RNA concentration using a NanoDrop ND-1000 spectrophotometer (Thermo Fisher Scientific).

For all species except *C. hollandica*, we immediately proceeded to digest genomic DNA using the TURBO DNA-free kit (Ambion, Thermo Fisher Scientific), following the manufacturer’s protocol with a 30 minute incubation. After digestion, we removed DNase with DNase Inactivation Reagent for all species except *S. punica*, whose RNA extract was incompatible with the reagent. We instead removed DNase from *S. punica* by extracting with two volumes of pH 8 phenol:chloroform:isoamyl alcohol, removing residual phenol with two volumes of chloroform:isoamyl alcohol, and precipitating with 0.3 M sodium acetate pH 5.2 (all three from Sigma-Aldrich), 25 *μ*g of GlycoBlue (Thermo Fisher Scientific) as a carrier and two volumes of 100% ethanol. We washed the pellet in 70% ethanol and resuspended in 50 *μ*l of nuclease-free water. For *Didymoeca costata*, after DNase removal with the Inactivation Reagent, the RNA still appeared to be slightly contaminated with protein, so we performed a pH 8 phenol:chloroform extraction to remove it. For *C. hollandica*, we observed significant total RNA degradation in the presence of DNase buffer. Instead, to remove genomic DNA we performed three successive rounds of extraction with pH 4.5 phenol:chloroform:isoamyl alcohol (Sigma-Aldrich), followed by the chloroform:isoamyl and precipitation steps described above. To reduce the possibility of cross-contamination among samples, we performed RNA isolation and DNase digestion for a single culture at a time, we used disposable materials when possible, and we cleaned all other materials (bench tops, centrifuges, dry baths, pipettes, etc.) thoroughly with ELIMINase before use.

We evaluated total RNA on Bioanalyzer 2100 RNA Pico chips (Agilent Technologies, Santa Clara, California, United States) with four criteria to be considered high-quality: (1) four distinct ribosomal RNA peaks (16S and 23S for bacteria, 18S and 28S for choanoflagellates), (2) low signal in all other regions, as a non-ribosomal signal is evidence of degradation, (3) at least a 1:1 ratio of 28S ribosomal area to 18S ribosomal area, since 28S ribosomal RNA is likely to degrade more easily than is 18S ribosomal RNA, and (4) an RNA Integrity Number (RIN) of 7 or greater (Schroeder et al., 2006). (We note that the Bioanalyzer software could not calculate RIN for several cultures.) If we were not able to obtain high-quality total RNA after the first attempt for any culture, we repeated cell growth, centrifugation and total RNA isolation up to a maximum of 5 times, and selected the best available total RNA sample to use for transcriptome sequencing. We produced a rough estimate of the amount of choanoflagellate total RNA present in each sample by calculating the ratio of choanoflagellate to bacterial ribosomal RNA peaks (18S vs. 16S and 28S vs. 23S) and multiplying the resulting fraction by the total amount of RNA present in the sample (Supp. Table 1).

### Test of polyA selection to separate choanoflagellate from bacterial RNA

The standard library preparation protocol for Illumina mRNA sequencing used poly-dT beads to separate polyadenylated mRNA from other types of non-polyadenylated RNA such as rRNA and tRNA. For choanoflagellates, the bead selection step also serves to separate choanoflagellate mRNA from bacterial RNA (which is not polyadenylated). Because the amount of bacterial RNA isolated from a culture often exceeded the amount of choanoflagellate RNA by one to several orders of magnitude, we reasoned that the standard bead selection might not be sufficient. We tested this hypothesis on *S. rosetta* Px1, a culture grown with a single species of bacterium, *Algoriphagus machipongonensis*. Because both species have sequenced genomes (Alegado et al., 2011; Fairclough et al., 2013), we could identify the origin of sequenced RNA using a straightforward read mapping procedure. We cultivated *S. rosetta* Px1 cells as described previously (Fairclough et al., 2010). We scraped and harvested 50 ml of culture after three days of growth in a 150 ml tissue culture dish. We performed centrifugation (with a 10 minute spin), RNA isolation, DNase digestion and total RNA quality assessment as described above.

We compared the standard Illumina TruSeq v2 mRNA preparation protocol, which performs two rounds of polyA selection with a single set of poly-dT-coated beads, against a modified protocol that repeats the polyA selection steps, for a total of four rounds of polyA selection with two sets of beads. For all other aspects of library preparation, we followed the manufacturer’s protocol. We quantified libraries by qPCR (Kapa Biosystems, Sigma-Aldrich) and sequenced them on a HiSeq 2000 machine (Illumina, San Diego, California, United States) at the Vincent J. Coates Genomics Sequencing Laboratory at the California Institute for Quantitative Biosciences (Berkeley, California, United States).

We generated 16,970,914 single-end 50 bp reads for the library prepared with two rounds of polyA selection, and 17,182,953 for the four-round library. We mapped reads to the *S. rosetta* and A. *machipongonensis* genomes using BWA version 0.6.1 (Li and Durbin, 2009) and SAMtools version 0.1.18 (Li et al., 2009) with default parameter values. We counted reads mapping to *S. rosetta* ribosomal loci on supercontig 1.8 (5S: positions 1900295-1900408, 18S: 1914756-1916850 and 28S: 1917502-1923701). The number of reads mapping to the non-ribosomal portion of the *S. rosetta* genome did not differ substantially between the two data sets: 12,737,031 reads mapped for the two-round data, and 12,585,647 for the four-round data. Similarly, 10,509,262 reads from the two-round data mapped to *S. rosetta* transcripts and 10,181,522 for the four-round data. We also asked whether additional rounds of polyA selection would cause increased RNA breakage due to pipetting or heating during the selection process [e.g., (Kingston, 2001)], leading to lower coverage of the 5’ ends of transcripts (because the poly-dT sequence binds to the 3’ end of RNA molecules). We estimated the loss of 5’ transcript ends due to shear to affect less than roughly 5% of transcripts (Supp. Figure 2a).

The four-round method removed roughly an order of magnitude more non-polyadenylated RNA than the two-round method (Supp. Figure 2b). We observed that the four-round data set had a slightly lower overall read quality. To address this, we tested whether a difference in read quality between the two data sets could account for the difference in read mapping by randomly resampling the two-round data set to contain the same number of either Phred-like quality 20 or Phred-like quality 30 bases as the four-round data set, but neither resampling affected the results. We also tested whether transcript assembly quality would suffer in the four-round data set by assembling both data sets *de novo* with Trinity release 2012-03-17 (Grabherr et al., 2011) with default parameter values and mapping the resulting contigs to the *S. rosetta* genome using BLAT version 34 (Kent, 2002) with default parameter values, but we observed no substantial difference between the two data sets.

Given the superior ability of four rounds of polyA selection to remove contaminating bacterial RNA with little to no loss of transcript coverage, we adopted this methodology for subsequent transcriptome sequencing. The raw sequence reads for this experiment are available at the NCBI Short Read Archive with the BioProject identifier PRJNA420352.

### Library preparation and mRNA sequencing

We began the Illumina TruSeq v2 mRNA library preparation protocol with approximately 2 *μ*g of total RNA per culture, if available (Supp. Table 1). We performed four rounds of polyA selection (instead of the standard two) and introduced two further modifications to the standard protocol: first, we repeated the Agencourt AMPure XP (Beckman Coulter, Indianapolis, Indiana, United States) bead clean-up step to enhance adapter removal, and second, we used 1.5 *μ*l less volume in all bead elution steps, in order to reduce loss during the protocol. We prepared libraries from 5 RNA samples at a time, and the libraries were later multiplexed into groups of 6 or 7 per sequencing lane. To allow us to detect evidence of potential cross-contamination during either process, we ensured that the groupings for sample preparation and sequencing were distinct (Supp. Table 1).

We estimated library concentration using the Qubit dsDNA HS Assay (Thermo Fisher Scientific) and determined quality and fragment size distribution with a Bioanalyzer 2100 High Sensitivity DNA chip (Agilent). We quantified libraries by qPCR (Kapa Biosystems, Sigma-Aldrich) and sequenced them on an Illumina HiSeq 2000 at the Vincent J. Coates Genomics Sequencing Laboratory at the California Institute for Quantitative Biosciences (Berkeley, California, United States). One group of libraries was sequenced twice (consisting of A. *spectabilis, Diaphanoeca grandis, Helgoeca nana, S. helianthica, S. infusionum* and *Salpingoeca urceolata*) due to a drop-off in quality after base 50 on the forward read of sequencing pairs; quality scores up to base 50 on the forward read and on reverse reads were not affected. The second, repeat sequencing run did not experience this issue. We incorporated both sequencing runs for affected libraries into subsequent analyses (including Quality trimming and Error correction, see below). We produced between 23 million and 61 million paired-end 100 bp sequencing reads per library (Supp. Table 1). Raw sequence reads are available at the NCBI Short Read Archive with the BioProject identifier PRJNA419411 (accession numbers for each species are listed in Supp. Table 1).

### Quality trimming

We trimmed sequence reads using Trimmomatic version 0.30 (Lohse et al., 2012) with two separate filters: (1) removal of TruSeq adapter sequence and (2) trimming very low quality bases from the ends of each read. To implement these filters, we ran Trimmomatic in three phases. In the first phase, we clipped palindromic adapters using the directive ILLUMINACLIP:2:40:15 and discarded resulting reads shorter than 25 bases with MINLEN:25. This resulted in two data sets: one containing reads whose mate pair remained in the set, and the other composed of reads whose pair was removed due to adapter contamination. In the second phrase, we clipped simple adapters from the remaining paired data set using the directive ILLUMINACLIP:2:40:15, imposed a minimum Phred-like quality cutoff of 5 on the first ten and last ten bases using LEADING:5 and TRAILING:5, subjected the read to a minimum sliding window quality using SLIDINGWINDOW:8:5 and discarded resulting reads shorter than 25 bases with MINLEN:25. The third phase operated on the remaining unpaired reads from the first phase, and implemented the same directives as the second phase. We used a permissive minimum quality of 5 in order to remove very low quality bases, as these might interfere with read error correction in the subsequent processing step. We discarded reads less than 25 in length because they were shorter than the k-mer size of the Trinity assembler (see *De novo* transcriptome assembly below). In all adapter clipping operations, we used sequences appropriate to the index used for multiplexed sequencing. The number of sequence reads and total bases remaining after trimming for each library are given in Supp. Table 1.

### Error correction

We performed error correction on trimmed reads using Reptile v1.1 (Yang et al., 2010) following the authors’ instructions, with the modifications described below. We began by using the ‘fastq-converter.pl’ script to convert from FASTQ and to discard reads with more than 1 ambiguous character “N” in any window of 13 bases. For reads with one “N”, we chose the character “a” as the substitute for “N”, as all of the characters in our input reads were in upper case (A, C, G, or T); thus, we could later recognize “N” bases converted in this step. Next, we tuned parameters using the ‘seq-analy’ utility following the authors’ instructions, in four steps: (1) Running ‘seq-analy’ with default settings. (2) Adjusting the input settings to ‘seq-analy’ using the results from step 1. For all species, we set MaxBadQPerKmer to 8 and KmerLen to 25 (to match the k-mer length used in Trinity). (3) Re-running ‘seq-analy’ using the adjusted input settings. (4) Creating the input settings to ‘Reptile’ based on the output of step 3. We set KmerLen to 13 and step to 12 for all species. The values of QThreshold, T_expGoodCnt, T_card and Qlb differed by species (Supp. Table 1). All other parameters were left at their defaults to run Reptile.

We noticed that the locations within reads of errors identified by Reptile fell into two general classes: sporadic errors not located adjacent to any other error, and clustered errors, in which several adjacent bases within the same k-mer window were identified as errors. In some extreme cases, every single base within a sequence read was identified as a target for error correction; we observed this phenomenon in the set of read corrections for every species. We reasoned that this was an unintended consequence of the iteration-to-exhaustion approach (step 2d) of the Reptile algorithm. Therefore, we designed a method to correct sporadic errors, but not clustered errors. For each species, we began by grouping each read according to the total number of errors identified. Within each group, we built a distribution of the number of adjacent errors within the same k-mer window. For sporadic errors, this number should be close to 0, but for clustered errors, the number could be up to the k-mer size minus one. There was a clear pattern within each of these distributions, with some errors identified with no neighbors (sporadic errors), a smaller number identified with 1 neighbor, and an increasing number beginning at 2 or more neighbors (clustered errors). We used these empirical distributions to set the maximum allowable amount of neighboring errors within a k-mer window as the count just prior to the beginning of the secondary increase within each distribution. For example, for *D. grandis*, in the case of the group of reads containing 4 total identified errors, there were 316,103 errors with no neighbors within the same k-mer, 197,411 with one neighbor, 156,043 with 2 neighbors, and 353,639 with 3 neighbors (that is, all 4 errors were within the same k-mer window). Thus, for the group of reads containing 4 total identified errors in *D. grandis*, we only corrected errors with up to 2 neighboring errors in the same k-mer window. After running Reptile error correction of sequence reads and quality files subject to these cutoffs, we performed a final step of restoring ambiguous bases converted by ‘fastq-converter.pl’ (from “N” to “a”) that were not subsequently corrected by Reptile back to their original value of “N”.

### De novo *transcriptome assembly*

We performed *de novo* transcriptome assembly on trimmed, corrected sequence reads and quality files with Trinity release 2013-02-25 (Grabherr et al., 2011) with ‘--min_contig_length’ set to 150 and all other parameters at their default values. We chose a minimum contig length of 150 (rather than the default of 200) so as not to exclude assembly fragments that might encode predicted proteins with lengths between 50 and 66 amino acids, because some domains in the Pfam database are as short as 50 amino acids. Because none of the species we sequenced had an available genome assembly, we did not know whether transcripts might be encoded in overlapping positions within the genome. To test this possibility, we repeated each Trinity assembly with the addition of the ‘--jaccard-clip’ option and compared the estimated number of fusion transcripts predicted by Transdecoder release 2012-08-15 (Haas et al., 2013). We found essentially no difference in the number of predicted fusion transcripts between the original and ‘--jaccard-clip’ assemblies, and so we continued with the original assemblies. Assembly statistics are reported in Supp. Table 1.

### Identification and removal of cross-contamination

Cross-contamination within a multiplexed Illumina sequencing lane is estimated to cause incorrect assignment of roughly 0.5% of index pairs (Kircher et al., 2012). We designed a procedure to eliminate transcriptome assembly contigs resulting from incorrect index assignments. We ran blastn version 2.2.26 (Altschul, 1997) with a maximum E value of 1 x 10^−10^ to query contigs from each species against all other species. Because of the evolutionary distances among the choanoflagellates we sequenced (Supp. Figure 3), most contigs from one species had no matches in any other species. Within the contigs that did have cross-species matches (Supp. Figure 18a), we observed a large number that were identical or nearly-identical, which were likely cross-contaminants, and another set of matches distributed around roughly 80% identity, likely representing highly conserved genes. The two cases were separated at roughly 96% identity. After exploring the distribution of match lengths in a similar manner (Supp. Figure 18b), we considered matches at 96% or greater identity of at least 90 bases in length to be cross-contaminated.

Next, we identified the sources of cross-contaminated contigs by comparing the number of reads mapping from both species for each match. We first masked contigs with Tandem Repeats Finder version 4.04 (Benson, 1999), with the following parameter values: match = 2, mismatch = 7, indel penalty = 7, match probability = 80, mismatch probability = 10, min score = 30, max period = 24. We next mapped reads to masked contigs using the Burroughs-Wheeler Aligner, BWA, version 0.7.5a (Li and Durbin, 2009) and SAMtools version 0.1.18 (Li et al., 2009). We ran BWA ‘aln’ with the ‘-n 200’ option to allow up to 200 equally best hits to be reported, and all other parameter values left at their defaults. Based on the distribution of read mapping ratios between the pair of species matching for each cross-contaminated contig (Supp. Figure 18c), we retained only contigs for the species in a pair with 10 times or more reads mapping, and discarded all other contigs, with one exception: if a contig had at least 10,000 reads mapping, we did not discard it, regardless of read mapping ratio. We observed that many contigs encoding conserved genes (for example, α-tubulin and elongation factor 1α) also tended to be the most highly expressed, and thus the read mapping ratio was often close to 1 for these contigs. We identified as cross-contaminated and removed between 1.7% and 8.8% of contigs for each species (Supp. Table 1). We note that our procedure would also be expected to discard sequences from any bacterial species that were present in two different choanoflagellate cultures. For a more detailed examination of the cross-contamination removal process, see (Richter, 2013).

### Prediction of amino acid sequences from assembled transcripts and elimination of redundant transcripts

We predicted proteins from decontaminated contigs with Transdecoder release 2012-08-15 (Haas et al., 2013) with a minimum protein sequence length of 50. We noticed that many of the proteins originating from different contigs within a species were completely identical along their entire length. Furthermore, we also observed many contigs whose predicted proteins were a strict subset of the predicted proteins from another contig. For example, contig 1 might encode predicted proteins A and B, and contig 2 might encode two predicted proteins exactly matching A and B, and a third predicted protein C. We removed both types of redundancy (exact matches and subsets) from the set of predicted proteins, and we also removed the contigs from which they were predicted (Supp. Table 1).

### Measurement of expression levels and elimination of noise transcripts

To estimate expression levels, we mapped sequence reads to decontaminated, non-redundant, Tandem Repeats-masked contigs using the Burroughs-Wheeler Aligner, BWA, version 0.7.5a (Li and Durbin, 2009). We ran BWA ‘mem’ with the ‘-a’ option to report all equally best hits, with all other parameter values left at their defaults. We converted BWA output to BAM format using SAMtools version 0.1.18 (Li et al., 2009) and ran eXpress version 1.4.0 (Roberts and Pachter, 2012) with default parameter values in order to produce estimated expression levels, in fragments per kilobase per million reads (FPKM). The distribution of FPKM values (Supp. Figure 18d) had a peak near 1, with steep decreases in the number of contigs at lower values. Therefore, we chose an extremely conservative noise threshold two orders of magnitude below the peak, at FPKM 0.01, and discarded contigs (and their associated predicted proteins) below this value (Supp. Table 1).

The final sets of contigs are available as Supp. File 1, and the proteins as Supp. File 2. FPKM values for contigs are given in Supp. File 3.

### Completeness of predicted protein sets

To estimate the completeness of the predicted protein set for each species, we searched our data for conserved eukaryotic proteins with BUSCO version 3.0.2 (Simão et al., 2015). We used default parameter values and the 303 BUSCOs from the ‘eukaryota_odb9’ set, and performed searches with HMMER version 3.1b2 (Eddy, 2011). We note that each final transcriptome assembly contained between 18,816 and 61,053 proteins per species (Supp. Table 1), markedly more than the 9,196 and 11,629 genes predicted, respectively, from the assembled genomes of *M. brevicollis* (King et al., 2008) and *S. rosetta* (Fairclough et al., 2013). The relatively higher protein counts predicted from choanoflagellate transcriptomes likely represent an overestimate resulting from the inherent complexities in assembling unique contig sequences from short read mRNA sequencing data in the absence of a reference genome (Grabherr et al., 2011), including the fact that sequence reads from different splice variants may have assembled into separate contigs while being encoded by the same gene. Because our goal was to reconstruct the full diversity of genes in the Urchoanozoan and Urmetazoan, the tendency of transcriptomes to yield overestimates of gene numbers was not a significant concern.

### Construction of gene families and their probabilities of presence

To identify gene families, we chose a set of representative animals and outgroup species (Supp. Table 3). We used the 19 choanoflagellates we sequenced and the complete predicted protein sets from the *M. brevicollis* (King et al., 2008) and *S. rosetta* genomes (Fairclough et al., 2013). As an internal control, we had sequenced two independent isolates of *Stephanoeca diplocostata*, whose predicted proteins we compared using CD-HIT version 4.5.4 (Li and Godzik, 2006) with default parameter values. We found that the two *S. diplocostata* isolates contained essentially equivalent predicted protein sets, so we used only the Australian isolate for constructing orthologous groups. We compared the 21 choanoflagellate species to 21 representative animals with genome-scale sequence data available, with an emphasis on early-branching non-bilaterians: sponges, ctenophores, *T. adhaerens* and cnidarians. We included 17 outgroups with sequenced genomes in our analysis: *C. owczarzaki*, a holozoan representative of the closest relatives of animals and choanoflagellates, five fungi chosen to represent fungal diversity, two amoebozoa, and 9 species representing all other major eukaryotic lineages. The predicted proteins of the chytrid *Homolaphlyctis polyrhiza* were released in two partially redundant sets, which we combined using CD-HIT version 4.5.4 with default parameter values.

We applied OrthoMCL version 2.0.3 (Li, 2003) to construct gene families (orthologous groups of proteins) using recommended parameter values. As recommended in the OrthoMCL documentation, we performed an all versus all sequence homology search using blastp version 2.2.26 (Altschul, 1997) with an expectation value of 1 x 10^−5^, and we determined orthologous groups with MCL-edge version 12-068 (Enright, 2002; van Dongen and Abreu-Goodger, 2012) using an inflation parameter of 1.5.

Although most genes in most gene families were likely to be orthologous, no existing algorithm can yet perfectly separate orthologs from paralogs and spurious BLAST hits (Altenhoff et al., 2016). Approaches based on binary presence/absence calls and parsimony, developed for morphological characters, may be vulnerable to yielding hyperinflated estimates of the gene content of the Urchoanozoan, since any gene family containing at least one protein from animals and one from choanoflagellates would be inferred to have been present in the Urchoanozoan. Indeed, we observed a number of gene families containing many animals but only one or two choanoflagellates, and vice versa (Supp. Figure 4a). If these gene families represented true orthologous groups and were therefore present in the ancestral Choanozoan, they would require several independent loss events within one of the two groups; we reasoned that it was more parsimonious that some proportion of these genes evolved in only one of the two groups and that the isolated BLAST hits from the other group represented false positives. To address this problem, we produced membership probabilities for each protein within each gene family, based on its average BLAST E value to all other proteins within the cluster (the absence of a hit was treated as the maximum possible (i.e., least probable) E value). A true ortholog should have low E value hits to nearly all other members of its gene family, whereas a false ortholog should have higher E value hits to only one or a few members of the gene family (and therefore a very high average E value). We chose the lowest average gene family E value for each species within each gene family and rescaled them using the empirical cumulative density function (van der Vaart, 1998, p. 265) of all E values from the initial all versus all homology search, ordering them from highest to lowest, to produce a probability between 0 and 1 (Supp. Figure 4b). A protein with low E value BLAST hits to all other members of its gene family had a probability close to 1, and a protein with high E value BLAST hits to only a few members of its gene family had a probability close to 0 (Supp. Figure 4c-d). We found that most proteins fell close to one of these extremes, and that the probabilities clearly distinguished between hits between proteins within the same gene family versus hits between proteins in two different gene families (Supp. Figure 4e). In addition, proteins within gene families identified above containing many choanoflagellates and many animals had probabilities closer to 1, whereas proteins within gene families with few choanoflagellates and many animals contained both low probabilities, which are likely to be false orthologs, and high probabilities likely representing true orthologs (Supp. Figure 4f).

Protein sequences for gene families are available as Supp. File 4, and presence probabilities by species are listed as part of Supp. Table 5.

### Inference of gene family origins and heat map

For each gene family, we calculated the sum of membership probabilities for species from each of the five major groups in this study (choanoflagellates, animals, *C. owczarzaki*, fungi, and other eukaryotes). Based on the resulting distribution (Supp. Figure 5a), for a gene family to be considered present in a group, we required this sum to equal or surpass 10% of the number of species in the group. For choanoflagellates and animals, each of which have 21 species in our data set, this equates roughly to a gene family being represented at high probability in more than two species. Next, we developed a set of parsimony-based rules to determine the origin of each gene family (Supp. Table 4). If a gene family was present within two groups, we considered it to have been present in their last common ancestor. For gene families containing proteins from species in only one group, we considered it to have been present in the last common ancestor of that group only if it was present within two or more separate sub-groups (Supp. Figure 5b). For example, within animals, we defined three sub-groups: sponges *(O. pearsei, Ephydatia muelleri*, A. *queenslandica)*, ctenophores *(M. leidyi)*, and later-branching animals *(T. adhaerens*, cnidarians, bilaterians). An animal-specific gene family present in at least two of these three sub-groups would be considered to have been present in the Urmetazoan (and thus to have evolved on the animal stem lineage). Inferred group presences for each gene family are available in Supp. File 5.

In the text, we precede estimated gene family counts with a tilde. We selected a conservative probability threshold of 10% in order to minimize the number of gene families that might erroneously be assigned as specific to a given group (e.g., animals or choanozoans). This choice may have resulted in some gene families that are truly specific to a group instead being incorrectly assigned as shared with another group. As a consequence, counts of gene families originating in different groups should represent relatively conservative estimates.

To produce a heat map for visual display of gene families, we ordered them by their patterns of presence within the five major species groups. Within a given pattern (for example, absent in outgroups and fungi but present in *C. owczarzaki*, animals and choanoflagellates) we clustered gene families by Pearson correlation using Cluster 3.0 (de Hoon et al., 2004), with all other parameter values set to their defaults. We visualized heat maps using Java TreeView version 1.1.6r4 (Saldanha, 2004) and color palettes from ColorBrewer (Harrower and Brewer, 2003). For display purposes, we restricted Figure 2 to gene families inferred to have been present in at least one of six nodes representing last common ancestors of interest: Ureukaryote, Uropisthokont, Urholozoan, Urchoanozoan, Urchoanoflagellate, and Urmetazoan. The full heat map for all gene families with representatives from at least two species is shown in Supp. Figure 6. Gene families present, gained and loss at each ancestral node are listed in Supp. File 6.

### Test of gene family presence in recently released genomes

Because new genome sequences are continuously becoming available, we tested whether the ancestral gene content reconstruction we performed would be influenced by the addition of new genome-scale data from two key sets of species: early-branching animals and early-branching holozoans. We chose species with high-quality publicly available protein catalogs that would maximize the phylogenetic diversity added to our data set. For early-branching animals, we selected two sponges, the demosponge *Tethya wilhelma* (Francis et al., 2017) and the calcareous sponge *Sycon ciliatum* (Fortunato et al., 2014), and one ctenophore, *Pleurobrachia bachei* (Moroz et al., 2014). For early-branching holozoans, we selected the teretosporeans *C. fragrantissima, Ichthyophonus hoferi, Chromosphaera perkinsii* and *Corallochytrium limacisporum* (Grau-Bové et al., 2017). Because we planned to implement a best reciprocal BLAST approach, which might be confounded by paralogs present within any of the species, we began by removing redundancy separately for each species using CD-HIT version 4.6 (Li and Godzik, 2006) with default parameter values. We then performed best reciprocal blastp from the protein catalog of each additional species versus all 59 species included in our original analysis (Supp. Table 3), with the same maximum E value (1 x 10^−5^). We retained only top reciprocal blastp hits. We next calculated gene family presence probabilities for each additional species, using a slightly modified procedure. Because we retained only the top hit for each additional species protein to each original species, an additional species protein could only match to a single representative per species within each gene family, although the family might contain multiple representatives per species. Thus, instead of calculating average E value from each additional species protein to all members of each gene family (as we did for the original set), we calculated the average of best hits to each species within the family. Next, because each protein from an additional species might hit multiple different gene families, we chose the single gene family match with the lowest average E value. Finally, for each gene family and each additional species, we selected the lowest average E value of any protein in the species to the gene family and translated those to probabilities using the same empirical cumulative density function as for the original analysis (Supp. Figure 4b).

To visualize whether the additional species might substantially impact our inferences of ancestral gene content, we inserted their presence probabilities into the original heat map of Figure 2 without reordering gene families (Supp. Figure 7). The additional sponges display similar patterns of gene family presence probability to the original sponge species in the analysis, as does the additional ctenophore in comparison to the original ctenophore. Within the early-branching holozoans, *C. perkinsii* appears to show slight evidence for the presence of animal-specific gene families, but generally with low probabilities, roughly comparable to the choanoflagellate *S. dolichothecata*. Thus, we estimate that a very small proportion of animal-specific gene families would instead be classified as originating in Holozoa with the addition of early-branching holozoans. In addition, since we only included one representative early-branching holozoan (*C. owczarzaki*) in our original analysis, a subset of gene families originally classified as originating in Choanozoa would instead be assigned to Holozoa with the additional species; as such, we did not strongly emphasize the distinction between choanozoan-origin and holozoan-origin gene families in our results. Furthermore, as described below (Protein domains and evidence for gene presence based on domain architecture), the additional species would have a negligable effect on our inferences of gene family presence based on protein domain architecture.

### Species tree and phylogenetic diversity

Since the focus of this study was not phylogenetic tree construction, and because the topic has been thoroughly addressed elsewhere, we selected species trees from prior publications rather than building one based on our data. Furthermore, because our major findings were based on comparisons among groups whose phylogenetic relationships are well established (animals, choanoflagellates, *C. owczarzaki*, fungi and other eukaryotes), differences in tree topology within these groups should be of minor importance. For choanoflagellates, we used the species tree from (Carr et al., 2017), for animals (Philippe et al., 2009) and for other eukaryotes (Burki et al., 2016). Because the branching order of early animals is still under active debate, we depicted the relationships among early-branching animals as a polytomy (King and Rokas, 2017). We visualized the resulting tree with Archaeopteryx version 0.9813 (Han and Zmasek, 2009).

To measure phylogenetic diversity, we selected a set of 49 gene families for which each species had exactly one protein representative and no more than 5 species were missing (or roughly 10% of the 59 species in our data set). We aligned the protein sequences in each gene family separately using MAFFT version 7.130b (Katoh and Standley, 2013) with the parameters ‘--maxiterate 1000 --localpair’. We performed two rounds of alignment trimming with trimAl version 1.2rev59 (Capella-Gutierrez et al., 2009): in the first round, we used the parameter ‘-automated1’, and we supplied the output of the first round to the second round, with the parameter ‘-gt 0.5’. We constructed phylogenetic trees from the two-round trimmed alignments separately for each gene family with RAxML version 8.2.0 (Stamatakis, 2014), with the options ‘-m PROTGAMMALGF’ and ‘-f a -N 100’. We measured cophenetic distances between pairs of species on the resulting trees using the ape package version 4.1 (Paradis et al., 2004) and plotted pairwise distances averaged across all 49 gene families using the ggplot2 package version 2.2.1 (Wickham, 2009), both with R version 3.4.1 (R Core Team, 2017) in the RStudio development platform version 1.0.143 (RStudio Team, 2016).

### Group-specific core gene families found in all extant members

To identify animal-specific or choanoflagellate-specific gene families that were also present in all species within either group, we required every species in the group to pass the 10% minimum probability criterion. These core gene families are subject to several potential technical artifacts. First, an incomplete genome or transcriptome assembly could result in a species appearing to lack a gene family. Second, gene families containing repeated or repetitive protein domains (e.g. EGF or Ankyrin) might cause inappropriate inferences of sequence homology in our BLAST-based approach. Third, a gene family which duplicated on the lineage leading to the last common ancestor of a group could produce two gene families, among which paralogs are incorrectly partitioned, resulting in one or more species appearing to lack one of the two families. Thus, the lists of core gene families should not necessarily be considered as exhaustive, especially for serially duplicated or repeat-containing gene families.

We annotated animal-specific pan-animal gene families (Table 1) by selecting consensus features and functions in UniProt (Uniprot Consortium, 2017), FlyBase (Gramates et al., 2017) and WormBase (Lee et al., 2017). Notably, BTB/POZ domain-containing protein 9, whose function is relatively poorly characterized in comparison to other core animal-specific gene families, contains the BTB Pfam domain, which was identified as part of an expanded repertoire of ubiquitin-related domain architectures in animals (Grau-Bové et al., 2015).

### Gene family retention

To determine retention of ancestral gene families within extant species, we began with the sets inferred to have evolved on the stem lineages leading to the Ureukaryote, Uropisthokont, Urholozoan, Urchoanozoan, Urchoanoflagellate, and Urmetazoan (Supp. Table 4). For each set of gene families partitioned by ancestral origin, we summed the membership probabilities for each species. Because we applied a 10% membership probability threshold to consider a gene family to be present within group of species (see above, Inference of gene family origins and heat map), a species in one group might have a small residual sum of membership probabilities for gene families assigned to another group. As an example, some choanoflagellates may display a small amount of retention of animal-specific gene families, which represents the sum of non-zero membership probabilities that did not reach the 10% threshold.

To test whether *B. floridae, N. vectensis*, and *O. pearsei* retained the same gene families, we applied the 10% presence probability threshold within each species. We found that there were 7,863 Urmetazoan gene families retained in at least one of the three species and 5,282 retained in all three (67%).

Some animal species with draft genomes, for example *P. pacificus*, were among those that retained the fewest ancestral gene families. However, the lack of gene predictions resulting from an incomplete genome assembly is likely not as strong as the signal produced by gene loss. In the example case of *P. pacificus*, its sister species *C. elegans*, which has a finished genome, retains the second fewest gene families. We produced phylogenetic tree-based visualizations using the Interactive Tree of Life (iTOL) web site (Letunic and Bork, 2016). We produced bar charts using the ggplot2 package version 2.2.1 (Wickham, 2009) with R version 3.4.1 (R Core Team, 2017) in the RStudio development platform version 1.0.143 (RStudio Team, 2016).

### Gene ontology and pathway analysis

We annotated gene family function using the PANTHER Classification System (Thomas, 2003). We used the PANTHER HMM library version 7.2 (Mi et al., 2010) and the PANTHER HMM Scoring tool version 1.03 (Thomas et al., 2006) with default parameter values and the recommended expectation value cutoff of 10^−23^. We used PANTHER-provided files to associate PANTHER HMM hits with Gene Ontology (GO) terms (Ashburner et al., 2000). Annotations for both sets of terms, for all input proteins (including those not placed into gene families) are available in Supp. File 7. Annotations by gene family are listed in Supp. File 8.

To measure GO term enrichment, we used Ontologizer version 2.1 (Robinson et al., 2004; Grossmann et al., 2007) with the Parent-Child-Union method of comparison, a Bonferroni correction for multiple testing, and an adjusted p-value cutoff of 0.05. We accepted gene family GO annotations present in at least half of the proteins in the family. We assessed the enrichment of animal-specific gene families by comparing against the set of gene families present in the Ureukaryote. We considered the Ureukaryote set to be the most appropriate comparison since gene families that evolved in Holozoa, Choanozoa or Opisthokonts are themselves enriched for GO terms associated with animal gene families (Supp. Fig. 6).

We compared pathways present in the Urholozoan, Urchoanozoan and Urmetazoan using MAPLE version 2.3.0 (Takami et al., 2016). As input, we provided protein sequences only for species descending from each ancestral node. We supplied the following parameters to MAPLE: NCBI BLAST; single-directional best hit (bi-directional best hit would not have been appropriate, since our input database consisted of gene families containing closely-related proteins from multiple species); KEGG version 20161101; and organism list “ea” (all eukaryotes in KEGG). We compared completeness based on the WC (whole community) module completion ratio. We listed modules which differed by 25% or greater in completeness between the Urchoanozoan and Urmetazoan in Supp. Table 9, and between the Urholozoan and Urchoanozoan in Supp. Table 10. To focus on amino acid biosynthesis pathways (Supp. Figure 12), we exported KEGG results from MAPLE for the Urchoanozoan and Urmetazoan gene sets and compared them using the KEGG Mapper Reconstruct Pathway tool, pathway ID 01230 (Kanehisa et al., 2012).

### Evidence for gene presence based on protein domain architecture

We predicted protein domains with the Pfam version 27.0 database and pfam_scan.pl revision 2010-06-08 (Punta et al., 2012), which used hmmscan from the HMMER 3.0 package (Eddy, 2011). We performed all Pfam searches with default parameter values. We predicted signal peptides and transmembrane domains using Phobius version 1.01 (Käll et al., 2004) with default parameter values. Domains for all input proteins (including those not placed into gene families) are listed in Supp. File 7. Domains by gene family are shown in Supp. File 8.

To determine which animal-specific gene families lacked a Pfam domain of known function, we calculated the proportion of proteins in each animal gene family that were annotated with a given Pfam domain (Supp. Figure 19). We accepted Pfam domains present in at least 10% of proteins in a gene family. Domains of unknown function in the Pfam database had names beginning with “DUF”. To ensure that these domain names were not assigned a function in a more recent version of the Pfam database published after our initial annotation, we checked against Pfam version 31.0 and considered all domains whose names no longer began with “DUF” as having an assigned function.

We established a set of criteria for the presence of genes of interest, based on domain content and OrthoMCL gene families (Supp. Table 6). In all cases, ‘strong’ evidence for conservation consisted of the canonical domain architecture (a diagnostic domain or set of domains in a specific order). Because Pfam domains are constructed from a set of aligned protein sequences selected from sequenced species, they are often strongly enriched for animals (and, in particular, for animal models such as *D. melanogaster* and *C. elegans*) and therefore may be biased against detecting instances of protein domains with more remote homology. To address this concern, for some gene families, we considered the presence of a protein in the same gene family as another protein with ‘strong’ evidence to constitute potential ‘moderate’ or ‘weak’ evidence. In addition, we considered previous reports to be ‘strong’ evidence in two cases: the presence of a canonical TLR in *N. vectensis* (Miller et al., 2007; Sullivan et al., 2007), and the presence of NF-*x*B in *C. owczarzaki* (Sebé-Pedrós et al., 2011).

Genome or transcriptome data for 9 additional early-branching holozoans became available since our initial analyses of protein domain architecture (Torruella et al., 2015; Grau-Bové et al., 2017). To test whether the genes and protein domains of interest that we inferred to have originated in the Urchoanozoan did not in fact originate in the Urholozoan, we interrogated the domain content of the 9 new species. In all cases except one, there were no proteins in any of the nine species with domain architectures that would qualify as ‘strong’ evidence to change our inference of ancestral origin (i.e., where there was not already ‘strong’ evidence in *C. owczarzaki)*. The exception was the Glyco_hydro_56 domain in the filasterean *Ministeria vibrans*, which would be considered strong evidence for the presence of GAG hydrolase.

### Notch and Delta

For Notch, we defined ‘strong’ evidence for conserved domain architecture to be one or more EGF repeats, a Notch domain, a transmembrane domain and an Ank domain, in that order. This domain architecture was unique to animals and choanoflagellates in our data set. Many, but not all, Notch genes in animals also contain NOD and NODP protein domains. However, we did not use these as evidence, because numerous animal Notch genes do not encode these domains, including some bilaterians (Gazave et al., 2009) (for example, the primate *Macaca mulatta* (UniProt ID F7HCR6), the tapeworm *Hymenolepis microstoma* (A0A068XGW6), and the nematode *Ascaris suum* (U1NAR2)). In our data, the NOD and NODP domains were not present in any sponge or choanoflagellate species, although two of the three sponges *(E. muelleri* and A. *queenslandica)* encoded proteins with EGF, Notch, transmembrane and Ank domains, and a previous study found that A. *queenslandica* Notch and Delta were both expressed in in patterns consistent with other animals (Richards et al., 2008). We hypothesize that the NOD and NODP domains were not detected in either choanoflagellates or sponges because the NOD and NODP Pfam models were constructed from sequence alignments that included proteins mostly from bilaterians.

For Delta, we defined ‘strong’ evidence as the presence of both an MNNL (N terminus of Notch ligand) and a DSL domain. Both of these domains were considered to be animal-specific prior to this study (although we note that the DSL domain is found in two recently sequenced holozoans, the ichthyosporeans *C. fragrantissima* and *Pirum gemmata* (Grau-Bové et al., 2017)). The choanoflagellate *S. dolichothecata* expressed a protein encoding both the MNNL and DSL domains (m.249548), but lacking the transmembrane domain typically found in Delta. However, the contig encoding this protein was only partially assembled, and the presence of other partially assembled contigs in *S. dolichothecata* encoding combinations of DSL, transmembrane and EGF domains increases our confidence that *S. dolichothecata* encodes a *bona fide* Delta. (Predicted proteins encoding MNNL and DSL, but not a transmembrane domain, were also present in some animals in our data set; for example, *Capitella teleta* and *Strongylocentrotus purpuratus*.)

### TGF-β signaling pathway

The canonical TGF-β signaling ligand (e.g., Activins or BMPs) is characterized by the TGFb_propeptide (LAP) and TGF_beta domains (Munger et al., 1997), the type I TGF-β receptor by the combination of a TGF_beta_GS and a kinase domain, and the transcriptional activator SMAD by the MH1 and MH2 domains (Heldin et al., 1997). Although none of these combinations was present in choanoflagellates, we detected the individual TGFb_propeptide, TGF_beta_GS, MH1 and MH2 domains encoded in separate proteins (Supp. Figure 14). None of these four domains is found in eukaryotes outside Choanozoa (kinases are ancient domains, and the TGF_beta domain is found only in animals).

### Toll-like receptor signaling and innate immunity

For our analysis of Toll-like receptors, we did not distinguish among members of the LRR and TIR families (whose names in Pfam take the form of LRR_X, where X is a number, or TIR/TIR_2), because we found that different animal TLRs could contain domain combinations of each type. For example, *M. musculus* TLR11 (UniProt Q6R5P0, NCBI GI 45429992) contains LRR_8 and TIR_2 domains, whereas *M. musculus* TLR6 (UniProt Q9EPW9, NCBI GI 157057101) contains LRR_4 and TIR domains. We also did not attempt to differentiate among single vs. multiple cysteine cluster LRR domains (Imler, 2003), because we found that TLRs containing a multiple cysteine cluster LRRNT Pfam domain as the final LRR prior to the transmembrane region were restricted to *D. melanogaster* within our data set.

Although we did not find strong evidence for canonical animal TLR domain architecture in sponges, *M. leidyi* and *T. adhaerens*, we did detect proteins in sponges and cnidarians matching the domain content of animal interleukin receptors, which can also signal via MyD88 (O’Neill, 2008): an extracellular immunoglobulin (Ig) domain, a transmembrane domain, and an intracellular TIR domain (as found previously for the sponge A. *queenslandica* (Gauthier et al., 2010)). This architecture was not present in choanoflagellates nor any other non-animal in our data set. We note that, in our analyses, the proteins previously identified as TLR2 homologs in A. *queenslandica, O. carmela* and other sponges (Riesgo et al., 2014) appear to have domain architectures matching interleukin receptors or other proteins, but not TLRs.

For genes in the canonical *M. musculus* TLR/ NF-*κ*B signaling pathway, we collected information on domain architectures in the Pfam 31.0 database (which did not include the choanoflagellates sequenced in this study) and on the species present in the same gene family as the *M. musculus* protein (Supp. Table 14). With the exception of adapters and kinases containing the Death domain, there were no architectures that were both specific to animals and present across animal diversity, two features required of diagnostic architectures for a gene family. Similarly, we did not identify any gene families restricted to holozoans (as would be expected for genes participating in the TLR/ NF-*κ*B pathway) and whose choanoflagellate members we could unequivocally identify as orthologous to the *M. musculus* proteins in the family. Thus, we restricted our current analyses to TLRs, adapters containing TIR or Death domains, and NF-*x*B. Future analyses more focused on detecting individual signaling components [e.g., (Gilmore and Wolenski, 2012)] are likely to further elucidate their evolutionary histories.

We built an alignment for all proteins in the gene family containing SARM1 (gene family 6840), with the addition of the human sequence (UniProt Q6SZW1), since previous work has focused on the human SARM1 protein. We aligned proteins with MAFFT 7.130b (Katoh and Standley, 2013) with the parameters ‘--maxiterate 1000 --localpair’ for high accuracy alignments. We visualized the alignment with JalView 2 (Waterhouse et al., 2009). In addition to SARM1 and Kinase TIRs, choanoflagellates also encoded a diversity of protein domains associated with intracellular TIR domain-containing proteins (lacking a transmembrane domain). Domains found associated with TIR in multiple choanoflagellate species included ankyrin repeat (Ank) and armadillo (Arm) protein-protein interaction domains and Src homology 2 (SH2) kinase signal transduction domains.

In addition to TLRs, we also searched the choanoflagellates in our data set for cytosolic immune sensing genes. Several classes of nucleotide-binding domain and leucine-rich repeat (NLR) proteins have previously been detected in the *A. queenslandica* genome, suggesting that NLRs were present in the Urmetazoan (Yuen et al., 2014). The architecture of NLRs consists of a protein interaction domain (either Death or CARD), a nucleotide binding domain (NACHT) and multiple LRRs. Although we detected proteins encoding both NACHT and LRR domains in multiple choanoflagellates, neither the Death or CARD domain was present in any choanoflagellate we studied. The CARD domain in animal NLRs either directly or indirectly mediates activation of caspases (von Moltke et al., 2013), which are among the 36 core animal-specific gene families that we detected, suggesting that both NLRs and associated caspase downstream signaling activity may be animal innovations. Plants possess an analogous intracellular sensing pathway, NBS-LRR, which, like the TLR pathway, is thought to have evolved independently (Urbach and Ausubel, 2017). Diagnostic domains were absent in choanoflagellates for four additional cytosolic sensors: three containing CARD domains [mitochondrial antiviral signaling protein (MAVS), melanoma differentiation-associated protein 5 (MDA5) and retinoic acid inducible gene 1 (RIG-I)] and one containing HIN and PYRIN domains [absent in melanoma 2 (AIM2)]. However, we did find evidence for two other sensors previously reported in *M. brevicollis* (Wu et al., 2014): STING is present in both *Salpingoeca macrocollata* and *M. brevicollis*, based on the presence of the diagnostic Pfam domain TMEM173, and cGAS, identified by the Pfam domain Mab-21, is present in *S. macrocollata*, *M. brevicollis* and *M. fluctuans*.

### Glycosaminoglycan (GAG) synthesis genes

We identified GAG synthesis genes in *S. helianthica* using the list of human genes from Data S3 in (Woznica et al., 2017). We first standardized the identifiers of the human sequences to those in NCBI release GRCh38 of the human genome. Next, we ran blastp version 2.2.26 (Altschul, 1997) against *S. helianthica* with a maximum E value of 10^−5^ and selected the matching *S. helianthica* protein with the lowest E value. If a single *S. helianthica* protein hit two human proteins, we selected the lowest E value hit (and used the next best hit, if any, to the other human protein). In three cases, there was a *S. helianthica* BLAST hit for a human protein that had no homologs in *D. melanogaster, C. elegans, Hydra vulgaris (magnipapillata)*, or A. *queenslandica* in Woznica *et al*. We investigated these hits further by running blastp of the *S. helianthica* sequence against the NCBI nr database. In each case, we found that the NCBI nr BLAST hit did not match the function of the original human query protein. For the human dermatan sulfate synthesis gene D4ST (NP_569735.1), the *S. helianthica* top hit m. 244703 matched the chondroitin synthesis gene C4ST in the nr database; thus, we considered m.244703 to be a homolog of C4ST instead of D4ST. For the human hyaluronan synthesis genes HAS1 and HAS2 (NP_001514.2 and NP_005319.1), the *S. helianthica* top hits m.93480 and m.227356 both matched to chitin synthesis genes in the nr database, and not to hyaluronan synthesis genes; we thus discarded both of these hits. We note that neither *S. rosetta* nor *S. helianthica* had a match to the C6ST gene from the chondroitin sulfate synthesis pathway. However, *S. rosetta* was shown experimentally to produce chondroitin sulfate in Woznica *et al.;* it is possible that another gene present in both species, such as C4,6ST, may substitute for the function of C6ST.

To add *M. musculus, M. leidyi* and *O. pearsei* to the table of GAG synthesis genes, we used the same blastp-based process as for *S. helianthica. M. leidyi* and *O. pearsei* both had hits to the D4ST gene. In both cases, the best blastp matches of these proteins to the NCBI nr database were D4ST genes in other animals, indicating that both species are likely to encode this gene, and that it may have been present in the Urmetazoan and lost in *H. vulgaris, D. melanogaster* and *C. elegans*.

Four of the GAG synthesis genes (XYLT1, B4GT7, B3GT6 and B3GA3) participate in the synthesis of a linker region for both chondroitin sulfate and dermatan sulfate, but not hyaluronan (Kjellén and Lindahl, 1991; Mizumoto et al., 2013). When calculating the completeness of the chondroitin sulfate synthesis pathway, we included these four linker synthesis genes together with the seven chondroitin sulfate synthesis genes.

### GAG cleavage genes: hydrolases and lyases

The hydrolase and lyase GAG cleavage enzymes are frequently referred to as hyaluronidase and chondroitinase, respectively, although nearly all enzymes of both classes have some level of activity on both substrates (Yamagata et al., 1968; Stern and Jedrzejas, 2006; Kaneiwa et al., 2008). We therefore use the term GAG hydrolase in lieu of hyaluronidase, and GAG lyase in lieu of chondroitinase.

To build phylogenetic trees for GAG hydrolase and GAG lyase, we downloaded the sequences used to build the ‘Full’ alignments for their diagnostic domains from the Pfam 31.0 database (Finn et al., 2016): Glyco_hydro_56 for GAG hydrolase and Lyase_8 for GAG lyase. We appended sequences from gene families in our data set with at least one member encoding a diagnostic domain. We next ran CD-HIT version 4.6 (Li and Godzik, 2006), with default parameter values, for both genes in order to remove redundancy introduced by the additional sequences (because the species in our data set overlapped with those that were used to build the Pfam models). We aligned each gene separately using MAFFT version 7.130b (Katoh and Standley, 2013) with the parameters ‘--maxiterate 1000 --localpair’ for high accuracy alignments. After examining the alignments, we used trimAl version 1.2rev59 (Capella-Gutierrez et al., 2009) to trim GAG lyase with the option ‘-gt 0.8’ and GAG hydrolase with the option ‘-gt 0.6’. For GAG lyase, the trimmed alignment contained 4 proteins with no sequence data, which we discarded. We built phylogenetic trees using RAxML version 8.2.0 (Stamatakis, 2014), with the options ‘-m PROTGAMMALGF -f a -N 100’, and removed branches with less than 50% bootstrap support.

### GAG hydrolase (hyaluronidase) treatment of choanoflagellates

We performed an initial screen for qualitative effects of exogenous GAG hydrolase treatment at concentrations up to 500 *μ*g/ml (hyaluronidase type IV-S from bovine testes, Sigma-Aldrich) on four species of choanoflagellates with all four possible combinations of expressing/not expressing GAG hydrolase and forming/not forming rosette colonies. We tested *C. hollandica* (lacks GAG hydrolase, does not form rosettes), *M. fluctuans* (expresses GAG hydrolase, does not form rosettes), *S. rosetta* (lacks GAG hydrolase, forms rosettes) and *S. helianthica* (expresses GAG hydrolase, forms rosettes). We observed an effect of increased cells per colony in *S. helianthica* at GAG hydrolase concentrations ≥ 300 *μ*g/ml, but no effects in the other three species. Therefore, we continued with quantitative experiments on *S. helianthica*, with *S. rosetta* as a negative control.

We split actively growing cultures of *S. helianthica* and *S. rosetta* into new medium at a concentration of 10^5^ cells/ml. After 24 hours of growth at 25 °C, we treated cultures with concentrations of GAG hydrolase from 10 to 500 *μ*g/ml and incubated for 48 additional hours at 25 °C. We counted cells per rosette or cells per milliliter using gluteraldehyde-fixed (*S. helianthica*) or formaldehyde-fixed (*S. rosetta*) aliquots on a Bright-Line Hemacytometer (Hausser Scientific, Horsham, Pennsylvania, United States). We performed all counts blind to the identity of each sample. We counted 30 colonies per treatment.

We also tested two additional commercially available GAG cleavage enzymes (all from Sigma-Aldrich). We found that bovine hyaluronidase type VIII, a GAG hydrolase, had no effect on *S. helianthica*, but that it also formed visible precipitates at the concentrations we used for type IV-S (≥ 300 *μ*g/ml), which likely prevented its enzymatic activity. We also found no effects in *S. helianthica* for *Streptomyces hyalurolyticus* hyaluronidase, a GAG lyase, which is thought to have no enzymatic activity on chondroitin sulfate (Ohya and Kaneko, 1970; Park et al., 1997).

We performed scanning electron microscopy on *S. helianthica* colonies spun down and fixed onto silanized silica wafers as previously described (Dayel et al., 2011).

### RNAi machinery in choanoflagellates

To define ‘strong’ evidence for the presence of Argonaute in our data set, we used the conserved Pfam domain architecture DUF1785, PAZ and Piwi (Supp. Table 6). To detect ‘strong’ evidence for Dicer, we searched for the architecture Dicer_dimer, PAZ and Ribonuclease_3 (we considered the Dicer_dimer domain alone to be ‘moderate’ evidence). We did not detect Piwi in choanoflagellates, although it is present in most other eukaryotic lineages. Piwi is thought to repress transposable elements (Aravin et al., 2001, 2007), and *M. brevicollis*, the only choanoflagellate species that has been investigated for transposable elements, appears to have very few (Carr et al., 2008b).

Argonaute and Dicer were each lost five times independently in choanoflagellates (Supp. Figure 9), although these could both be overestimates if the genes were not expressed in our culture conditions. A similar pattern of repeated parallel loss has also been previously observed in kinetoplastids, a group of single-celled eukaryotes (Matveyev et al., 2017). Curiously, in contrast to the case for kinetoplastids, where Argonaute and Dicer were generally lost together, we detect five choanoflagellate species in which Argonaute is present and Dicer is absent or vice versa. These absences could reflect the presence of non-canonical RNAi genes or Dicer-independent RNAi pathways in choanoflagellates, as previously reported in *Saccharomyces cerevisiae* and in *M. musculus*, respectively (Drinnenberg et al., 2009; Cheloufi et al., 2010).

### Cell adhesion: Flamingo, Protocadherins, and Integrins

Diverse cadherins have previously been identified in choanoflagellates (Abedin and King, 2008; Nichols et al., 2012), but two classes of cadherins involved in animal planar cell polarity and development were thought to be animal-specific: Flamingo and Protocadherins.

The first described Flamingo cadherin (also known as starry night) is involved in the Frizzled-mediated establishment of planar polarity in the *D. melanogaster* wing (Chae et al., 1999; Usui et al., 1999). In animals, Flamingo cadherins are distinguished by the combination of three diagnostic Pfam domains, the presence of which constituted ‘strong’ evidence for Flamingo in our data set (Supp. Table 6): Cadherin, GPS and 7tm_2, a seven-pass transmembrane domain. The transcriptome of *C. hollandica* is predicted to encode a protein with all three diagnostic domains (Supp. Figure 10). In addition to these three domains, many animal Flamingo cadherins also contain HRM, EGF, and Laminin domains. Nine further choanoflagellate species express proteins with GPS, 7tm_2, and one of these additional domains (‘moderate’ evidence), and all choanoflagellate species express proteins with GPS and 7tm_2 domains (‘weak’ evidence).

Protocadherins are a large and diverse family of genes involved in animal development and cell adhesion (Frank and Kemler, 2002), and most, but not all, family members possess a diagnostic transmembrane Protocadherin domain paired with one or more extracellular Cadherin domains. Five species of choanoflagellates expressed proteins with a Protocadherin domain (‘strong’ evidence for conservation, Supp. Figure 10), although none of these also contained a Cadherin domain. The *Choanoeca perplexa* Protocadherin domain-containing protein also contains a transmembrane domain and an intracellular SH2 phosphorylated tyrosine binding domain, a combination not found in any other organism. Tyrosine kinase signaling networks greatly expanded in the Urchoanozoan (Manning et al., 2008; Pincus et al., 2008), and the Protocadherin domain in *C.perplexa* may have been co-opted for tyrosine kinase signaling function.

Integrins are thought to have been present in the Urchoanozoan, since homologs of both components of the animal integrin plasma membrane α/β heterodimer (Hynes, 2002) have previously been identified in *C. owczarzaki* (Sebé-Pedrós et al., 2010). We found ‘strong’ evidence for integrin β (in the form of the diagnostic Integrin_beta Pfam domain) in only a single species of choanoflagellate, *D. costata* (Supp. Figure 10). In contrast, we detected the integrin β binding protein ICAP-1 (Chang et al., 1997), which has been proposed to act as a competitive inhibitor (Bouvard et al., 2003), in all choanoflagellates except *M. brevicollis* and *S. rosetta*.

## Acknowledgements

We would like to thank Monika Abedin, Rosie Alegado, Hartmut Arndt, Roman Barbalat, Greg Barton, Cédric Berney, David Booth, Lauren Booth, Bastien Boussau, Candace Britton, Thibaut Brunet, Pawel Burkhardt, Minyong Chung, Matthew Davis, Mark Dayel, Stephen Fairclough, Xavier Grau-Bové, Nicolas Henry, Ella Ireland, Tommy Kaplan, Barry Leadbeater, Tera Levin, Matthew MacManes, Kent McDonald, Alan Marron, Scott Nichols, Frank Nitsche, Ryan Null, Mathilde Paris, Iñaki Ruiz-Trillo, Devin Scannell, Monika Sigg, Jason Stajich, Alberto Stolfi, Russell Vance, Jacqueline Villalta, Holli Weld, Jody Westbrook, Melanie Worley, Arielle Woznica, and Susan Young for assistance with laboratory techniques, bioinformatic and statistical analysis methods, helpful discussions, and comments on the manuscript. D.J.R. was supported by a National Defense Science and Engineering Graduate fellowship from the United States Department of Defense, a National Science Foundation Central Europe Summer Research Institute Fellowship, a Chang-Lin Tien Fellowship in Environmental Sciences and Biodiversity, a postdoctoral fellowship from the Conseil Régional de Bretagne, and the French Government “Investissements d’Avenir” program OCEANOMICS (ANR-11-BTBR-0008). P.F. was supported by NSF/EDEN IOS Grant Number 0955517. This work was supported in part by National Institute of General Medical Sciences grant R01 GM089977 to N.K. and by a grant of computer time from the DoD High Performance Computing Modernization Program at the ERDC DoD Supercomputing Resource Center. We thank the Analytical Lab at the Marine Science Institute of the University of California, Santa Barbara for performing nutrient analysis on choanoflagellate growth media. We are grateful to the Anheuser-Busch Coastal Research Center of the University of Virginia for hosting us during collection trips for choanoflagellate isolation. This study used the Vincent J. Coates Genomics Sequencing Laboratory at the University of California, Berkeley, supported by NIH S10 Instrumentation Grants S10RR029668 and S10RR027303.

## Supplementary Files

**Supplementary Figure 1.**
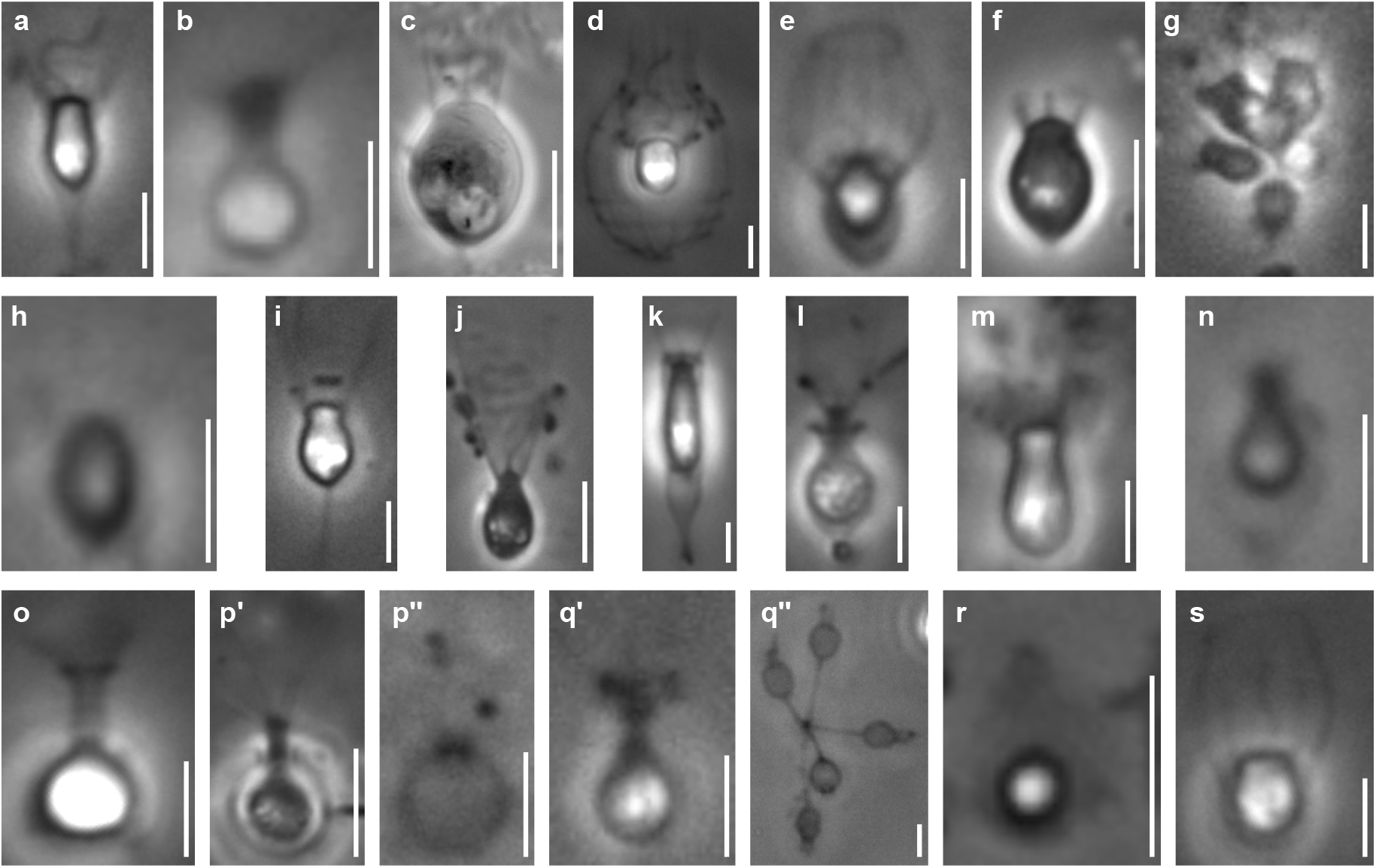
Phase contrast images of species sequenced in this study. All scale bars are 5 *μ*m. (a) *Acanthoeca spectabilis*, a nudiform loricate. (b) *Choanoeca perplexa*, a thecate craspedid. (c) *Codosiga hollandica*, a naked stalked craspedid. (d) *Diaphanoeca grandis*, a tectiform loricate. (e) *Didymoeca costata*, a tectiform loricate. (f) *Hartaetosiga balthica*, a naked stalked craspedid. (g) *Hartaetosiga gracilis*, a naked stalked craspedid; the image shows a colony of cells attached to the same stalk. (h) *Helgoeca nana*, a nudiform loricate. (i) *Microstomoeca roanoka*, a thecate craspedid. (j) *Mylnosiga fluctuans*, a naked craspedid. (k) *Salpingoeca dolichothecata*, a thecate craspedid. (l) *Salpingoeca helianthica*, a thecate craspedid. (m) *Salpingoeca infusionum*, a thecate craspedid. (n) *Salpingoeca kvevrii*, a thecate craspedid. (o) *Salpingoeca macrocollata*, a thecate craspedid. (p) *Salpingoecapunica*, a thecate craspedid; p’ represents a live cell within its theca, and p’’ an empty theca. (q) *Salpingoeca urceolata*, a thecate craspedid; q’ represents a live cell within its theca, and q’’ a group of empty thecae joined at the bases of their stalks. (r) *Savillea parva*, a nudiform loricate. (s) *Stephanoeca diplocostata*, a tectiform loricate. See (Leadbeater, 2015; Richter and Nitsche, 2016) for descriptions of choanoflagellate groups and their extracellular structures.

**Supplementary Figure 2.**
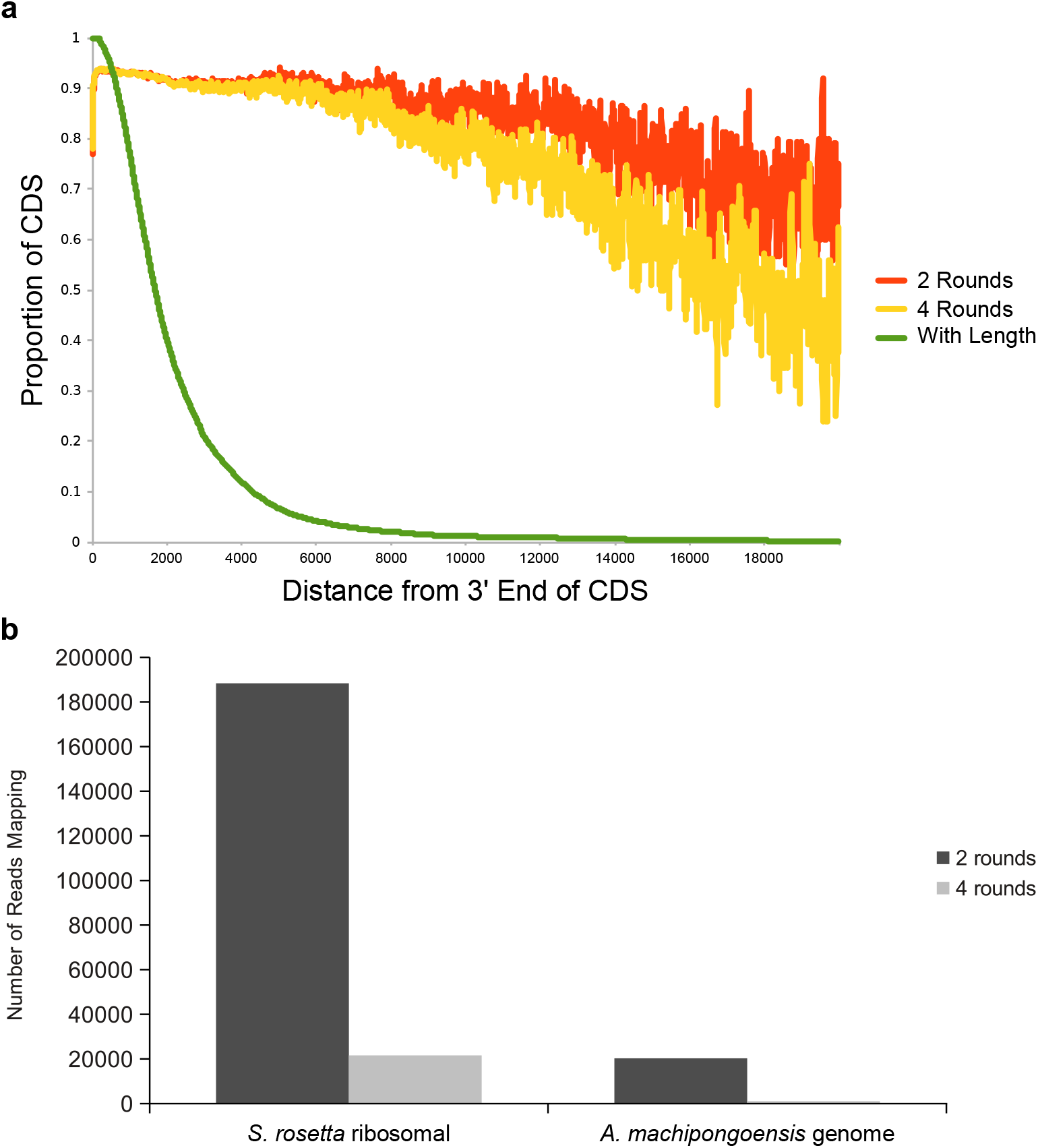
Tests of two versus four rounds of polyA selection. (a) An additional two rounds of polyA selection does not result in significant loss of read coverage at the 5’ ends of transcripts. The proportion of transcripts with at least one sequence read mapping at each given distance from their 3’ ends is plotted for two rounds (orange) and four rounds (yellow) of polyA+ selection. Also shown is the proportion of total transcripts in the *S. rosetta* genome at each length (green). There is a difference between the two round and four round coverage beginning at approximately 10,000 bases from the 3’ end of transcripts, but this represents only a very small fraction of total transcripts encoded in the *S. rosetta* genome. (b) The number of *S. rosetta* reads mapping either to the *S. rosetta* ribosomal locus or to the genome of its prey bacterium, *A. machipongoensis* (neither of which produces polyadenylated transcripts), after either two rounds or for rounds of polyA selection. Four rounds of polyA selection removes roughly an order of magnitude more non-polyadenylated RNA.

**Supplementary Figure 3.**
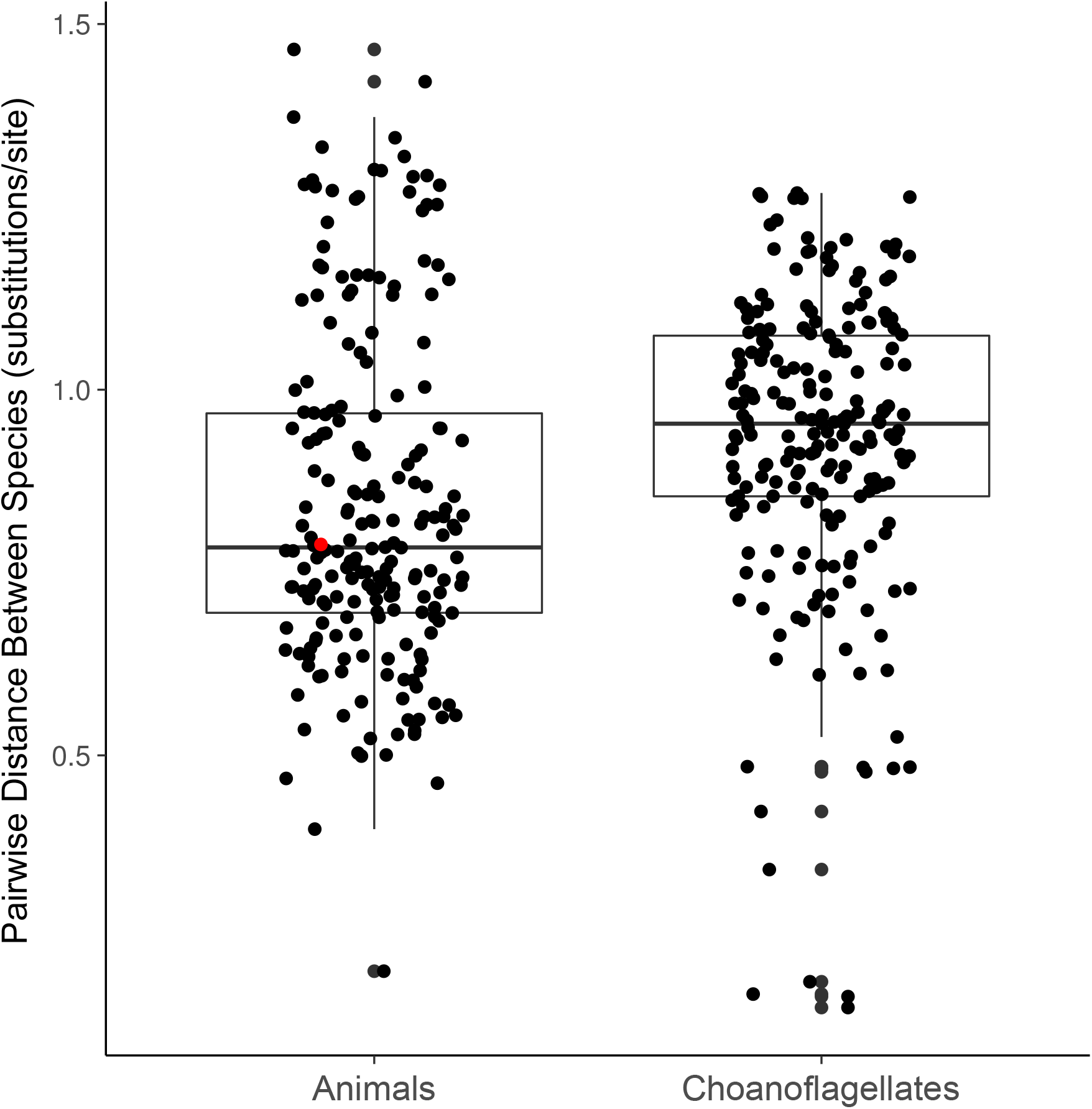
Distributions of phylogenetic diversity within choanoflagellates and within animals in our data set. Phylogenetic diversity is calculated as the average cophenetic distance between each pair of species from 49 separate phylogenetic trees constructed for conserved genes (see Methods). The red dot marks the cophenetic distance between the mouse *Mus musculus* and the sponge *Amphimedon queenslandica*.

**Supplementary Figure 4.**
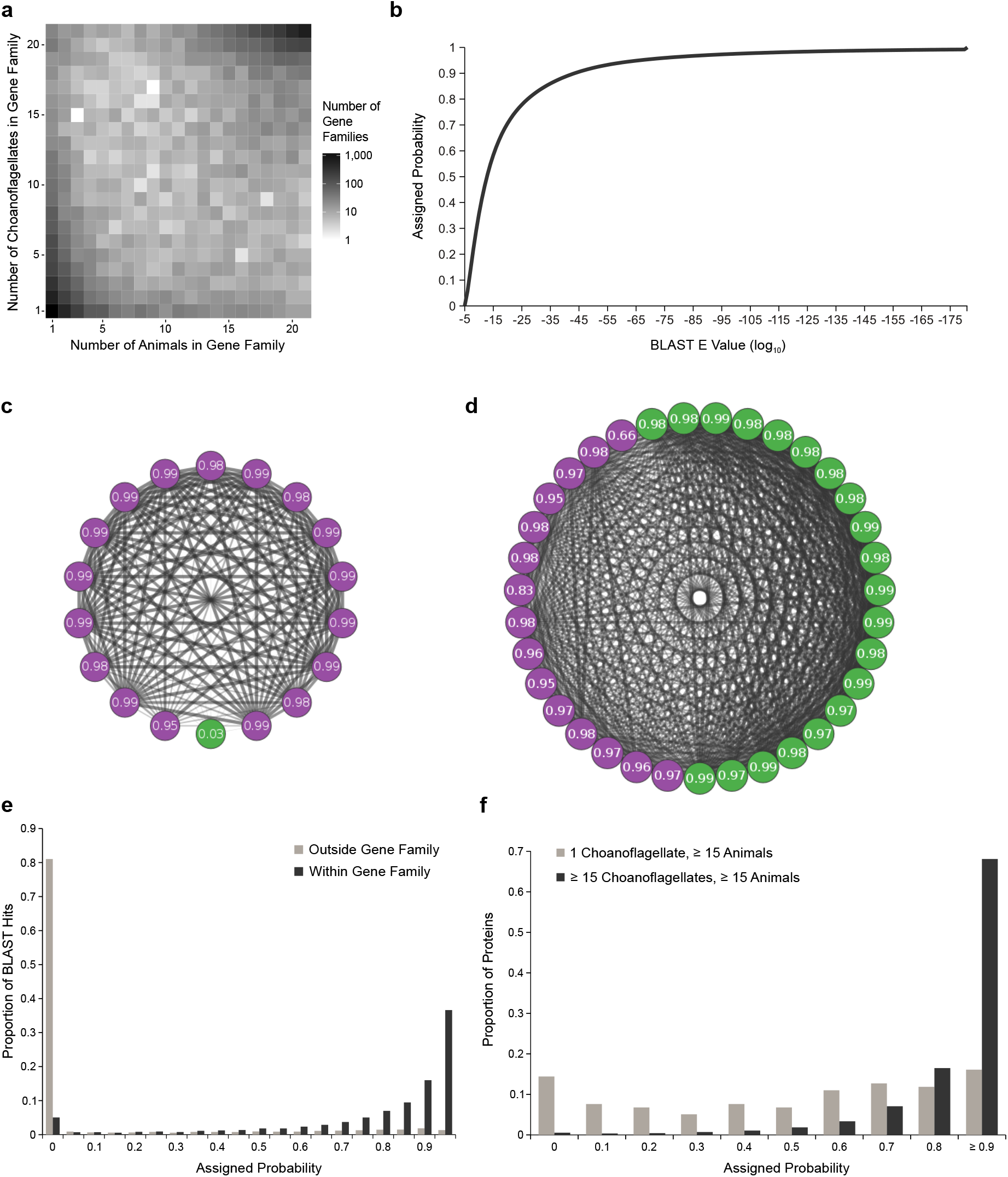
Gene family presence probabilities. (a) Joint distribution of the number of choanoflagellates and the number of animals in all gene families containing at least one animal and at least one choanoflagellate representative. Most gene families contain a similar number of animals and choanoflagellates, but there are also some gene families with many animals and few choanoflagellates or vice versa; these are candidates to contain false orthologs. (b) The empirical cumulative distribution function (van der Vaart, 1998, p. 265) of all BLAST E values. The x axis represents ordered BLAST E values (in log_10_, from highest to lowest), and the y axis represents the probability that they were assigned (from 0 to 1). (c-d) Network diagrams of BLAST hits among choanoflagellates (green) and animals (purple) and their assigned presence probabilities for two example gene families. Edge width and opacity is inversely proportional to BLAST E value. (c) Gene family 9066, which contains one choanoflagellate and 17 animals. The choanoflagellate protein sequence has high E value hits to only a subset of the other sequences in the cluster, and is assigned a low presence probability. The animal sequences are all connected by low E value hits, resulting in high probabilities. (d) Gene family 5870, which contains 20 choanoflagellates and 15 animals. (e) Histogram of the probabilities assigned to all BLAST hits, comparing hits between proteins in the same gene families, which are mostly composed of high probability hits with a small proportion of low probability hits that are likely spurious, versus hits to different gene families, which are less likely to represent orthologous relationships and are assigned low probabilities by our procedure. (f) Histogram comparing the probabilities assigned to all BLAST hits within gene families that contain at least 15 choanoflagellates and at least 15 animals, which are likely to be enriched for true orthologs, versus those present in 1 choanoflagellate and at least 15 animals, which are likely to be enriched for artifactual orthologs. For the first category, probabilities for both choanoflagellate and animal proteins are shown, whereas for the second only choanoflagellate probabilities are plotted.

**Supplementary Figure 5.**
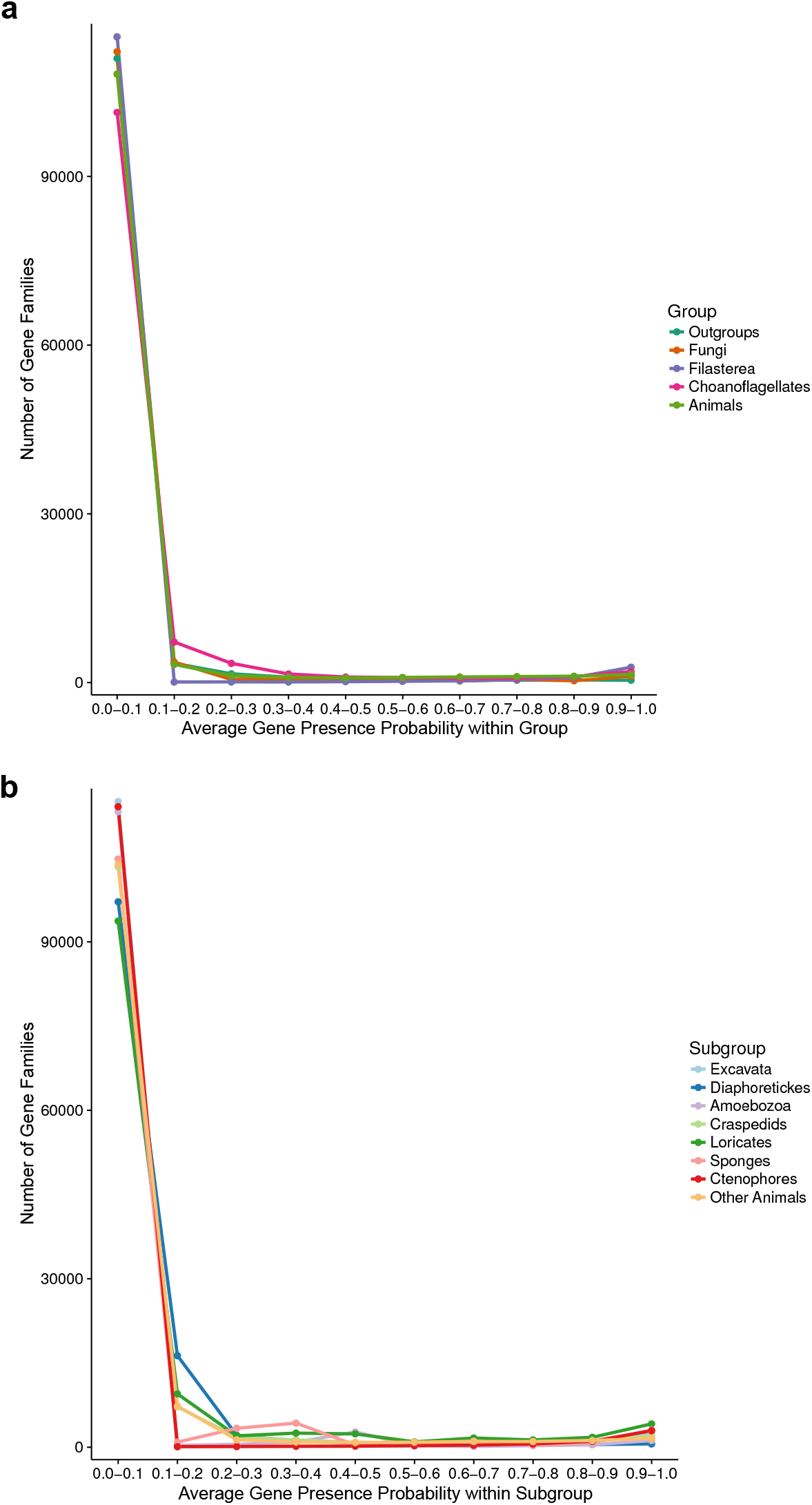
Distributions used to determine the 0.1 average probability threshold for inclusion in gene family analyses. (a) The average gene presence probability within each group of species. Filasterea contains only one representative, *Capsaspora owczarzaki*. (b) The average gene presence probability within each subgroup, used to determine whether a gene family only found in one group was present at the common ancestor of that group; such gene families must pass the 0.1 average probability threshold for at least two subgroups. Outgroups (subgroups: Excavata, Diaphoretickes, Amoebozoa), Choanoflagellates (subgroups: Loricates, Craspedids), Animals (subgroups: Sponges, Ctenophores, Other Animals).

**Supplementary Figure 6.**
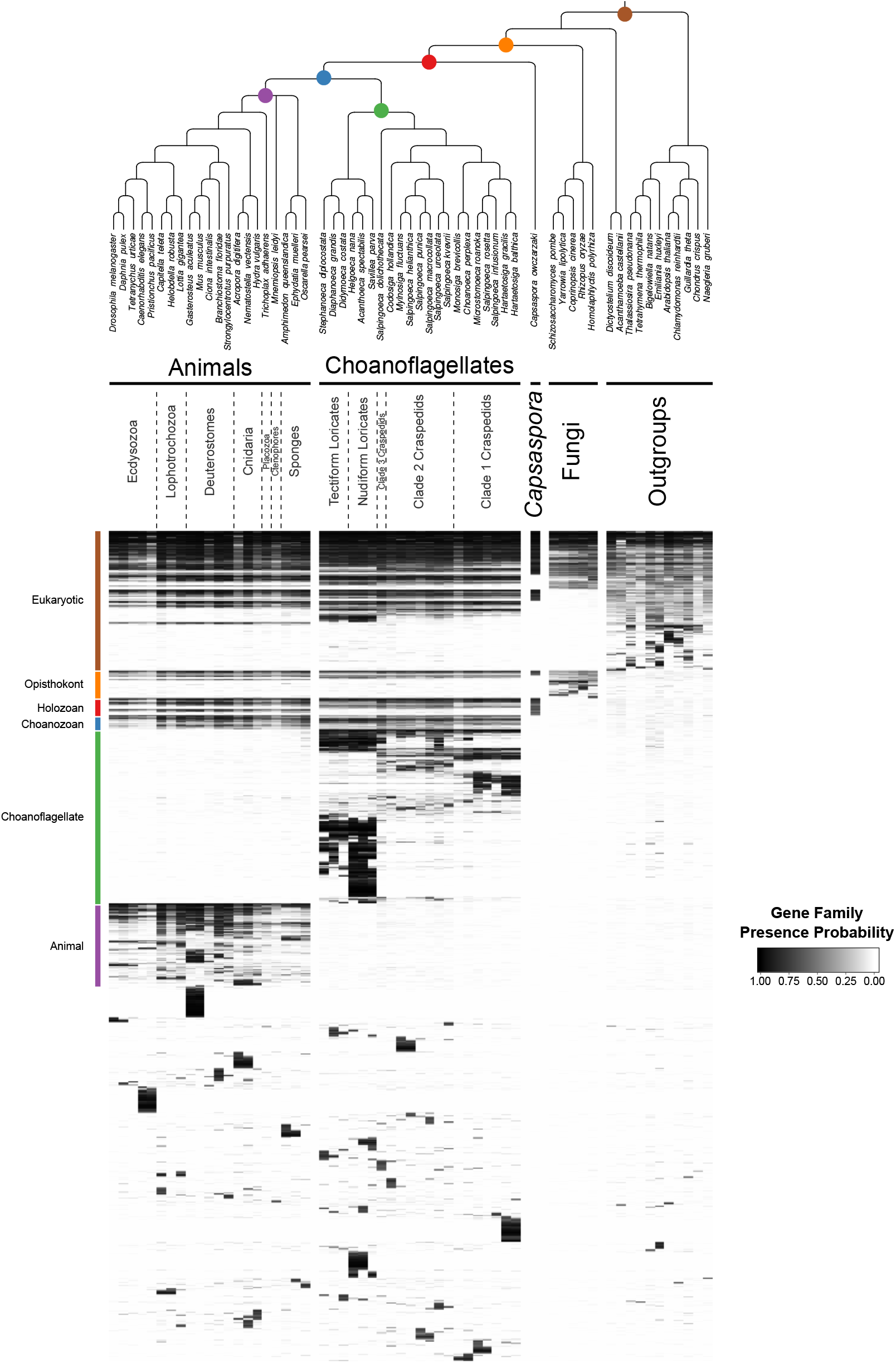
Full heat map of orthologous gene families present (with probability ≥ 0.1) in at least two species. The representation is the same as in Figure 2, with one major difference: gene family origin, indicated on the left, indicates that the average gene family probability within the group is at least 0.1, but does not imply that it was present in the last common ancestor of the group (as is the case in Figure 2).

**Supplementary Figure 7.**
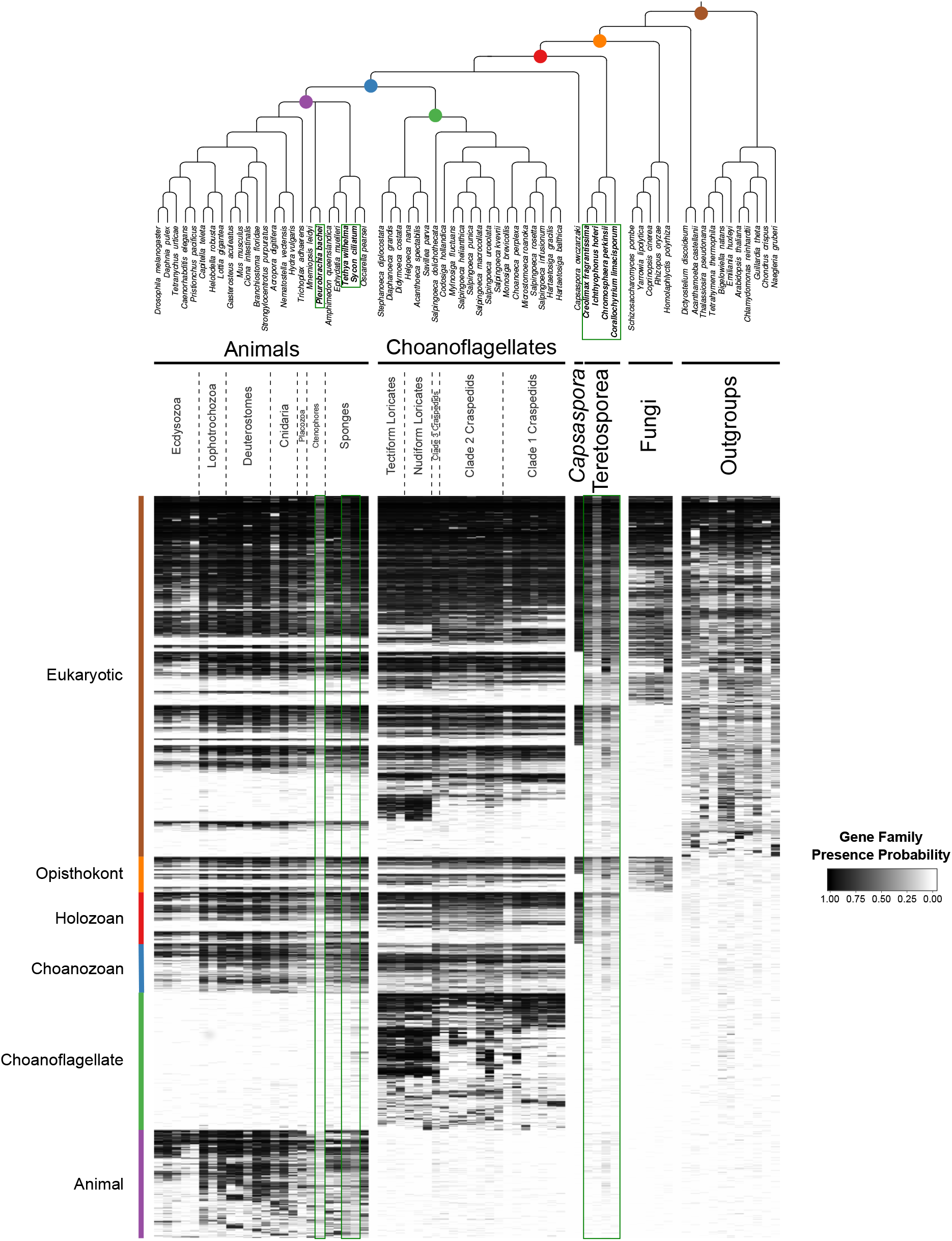
Heat maps with the inclusion of additional species. There are four additional teretosporeans (*Corallochytrium limacisporum, Chromosphaera perkinsii, Ichthyophonus hoferi* and *Creolimax fragrantissima*), two additional sponges (*Sycon ciliatum* and *Tethya wilhelma*) and one additional ctenophore (*Pleurobrachia bachei*). Additional species are in bold and enclosed by dark green boxes. We used the same clustering as in Fig. 2 (additional species were given no weight).

**Supplementary Figure 8.**
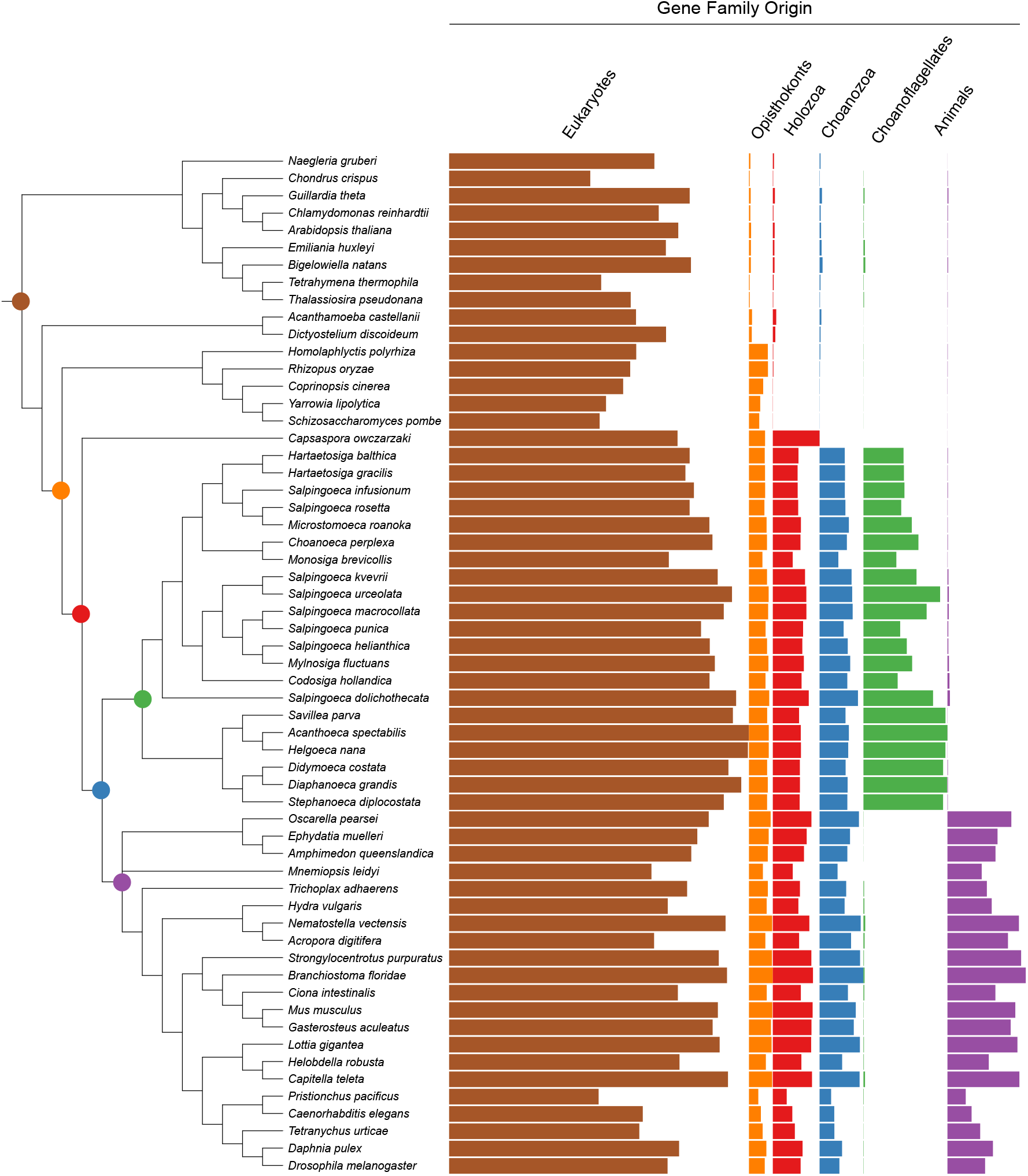
Phylogenetic tree with gene family retention for all species in this study. Gene families are divided by their origin in the last common ancestor in different groups: eukaryotic, opisthokont, holozoan, choanozoan, choanoflagellate, or animal. Colors correspond to nodes indicated in the phylogenetic tree. Bars represent the sum of presence probabilities for gene families with each origin. [Note that, for example, a small sum of probability is assigned to choanoflagellates for animal-specific gene families, and vice versa. This represents the sum of probabilities for the gene families that did not meet the 0.1 average presence probability threshold, but still had some residual probability between 0 and 0.1 (see Methods)].

**Supplementary Figure 9.**
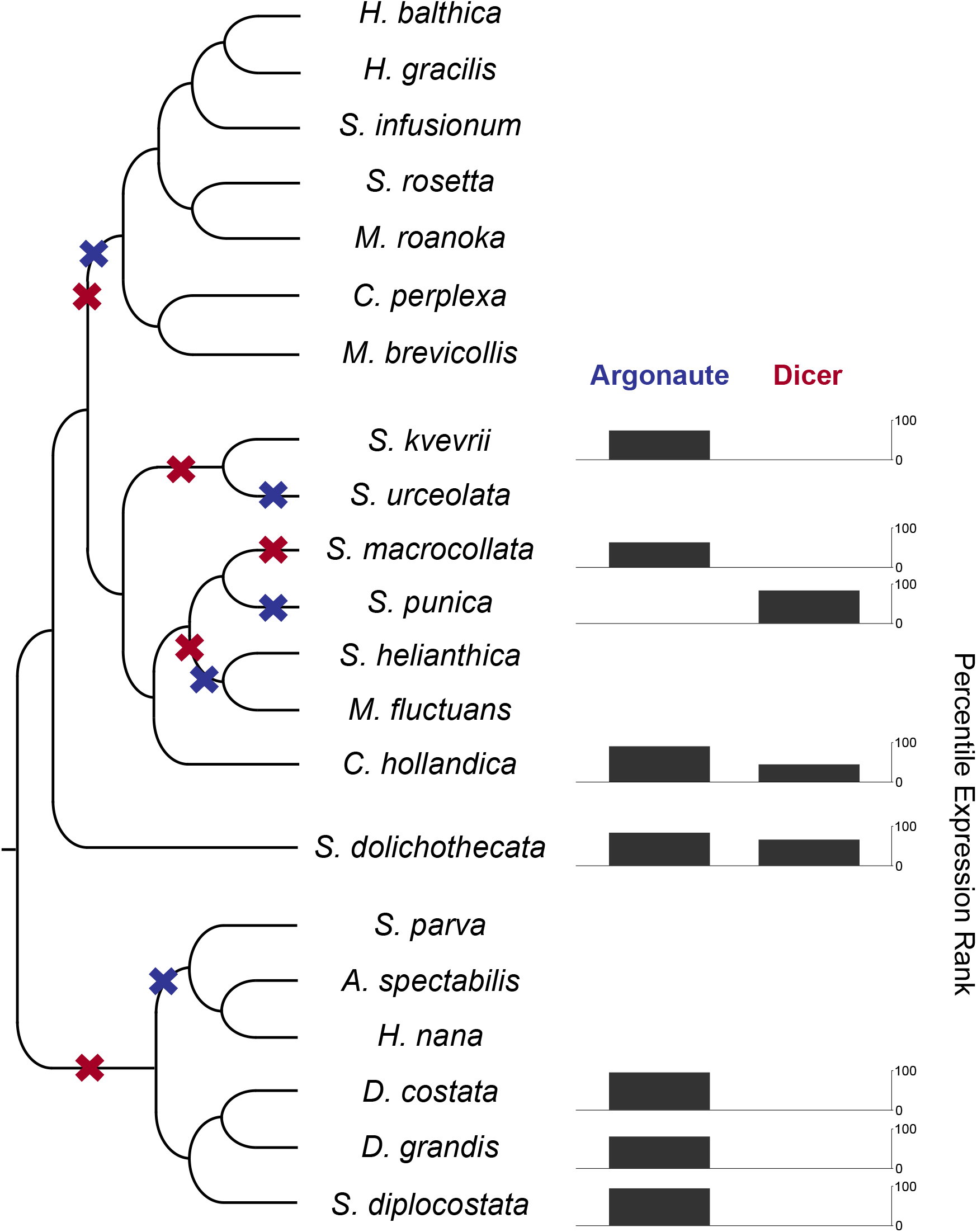
RNAi components in are present in choanoflagellates, but have been lost multiple times in different lineages. A phylogenetic tree of choanoflagellates with inferred losses of Argonaute (blue) and Dicer (red). For each species with moderate evidence for either Argonaute or Dicer, the expression level of that gene is shown as a percentile FPKM rank within each species. Species lacking a gene have no expression value. Overall, there have been multiple losses of both Argonaute and Dicer, and RNAi machinery appears to be entirely absent in the clade containing the two previously sequenced species, *Monosiga brevicollis* and *Salpingoeca rosetta*. In the species with Argonaute, it is highly expressed, and Dicer always shows a lower relative expression level.

**Supplementary Figure 10.**
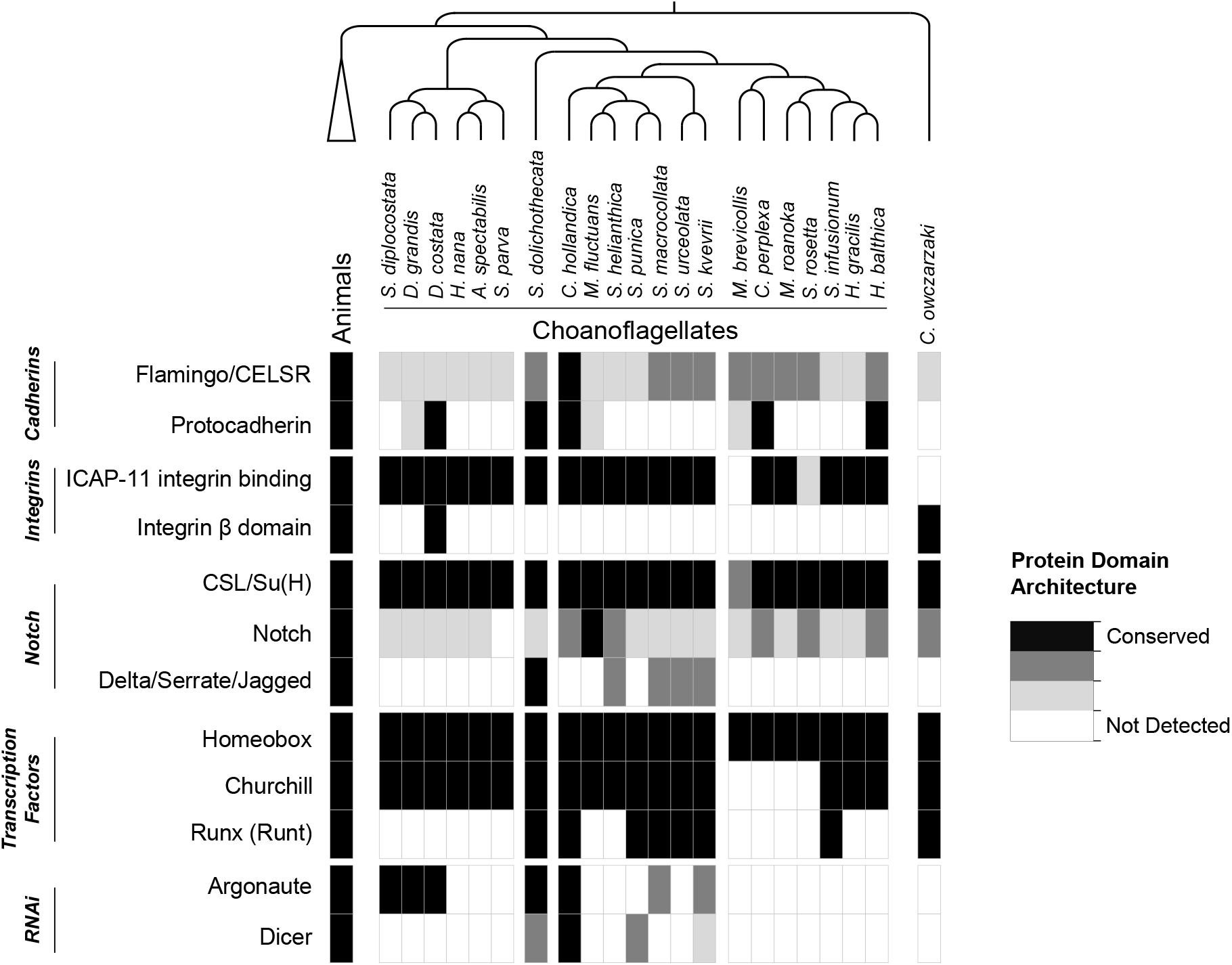
Presences of selected gene families in holozoans. Groups of gene families implicated in animal cell adhesion, cell-cell communication, and transcription regulation, in addition to RNAi components, are shown with the evidence for their presence in animals, choanoflagellates and *Capsaspora owczarzaki*. Criteria for gene family presences are based on diagnostic protein domain architectures (Supp. Table 6). CSL/Su(H): CBF-1/RBPJ-κ/Suppressor of Hairless/Lag-1.

**Supplementary Figure 11.**
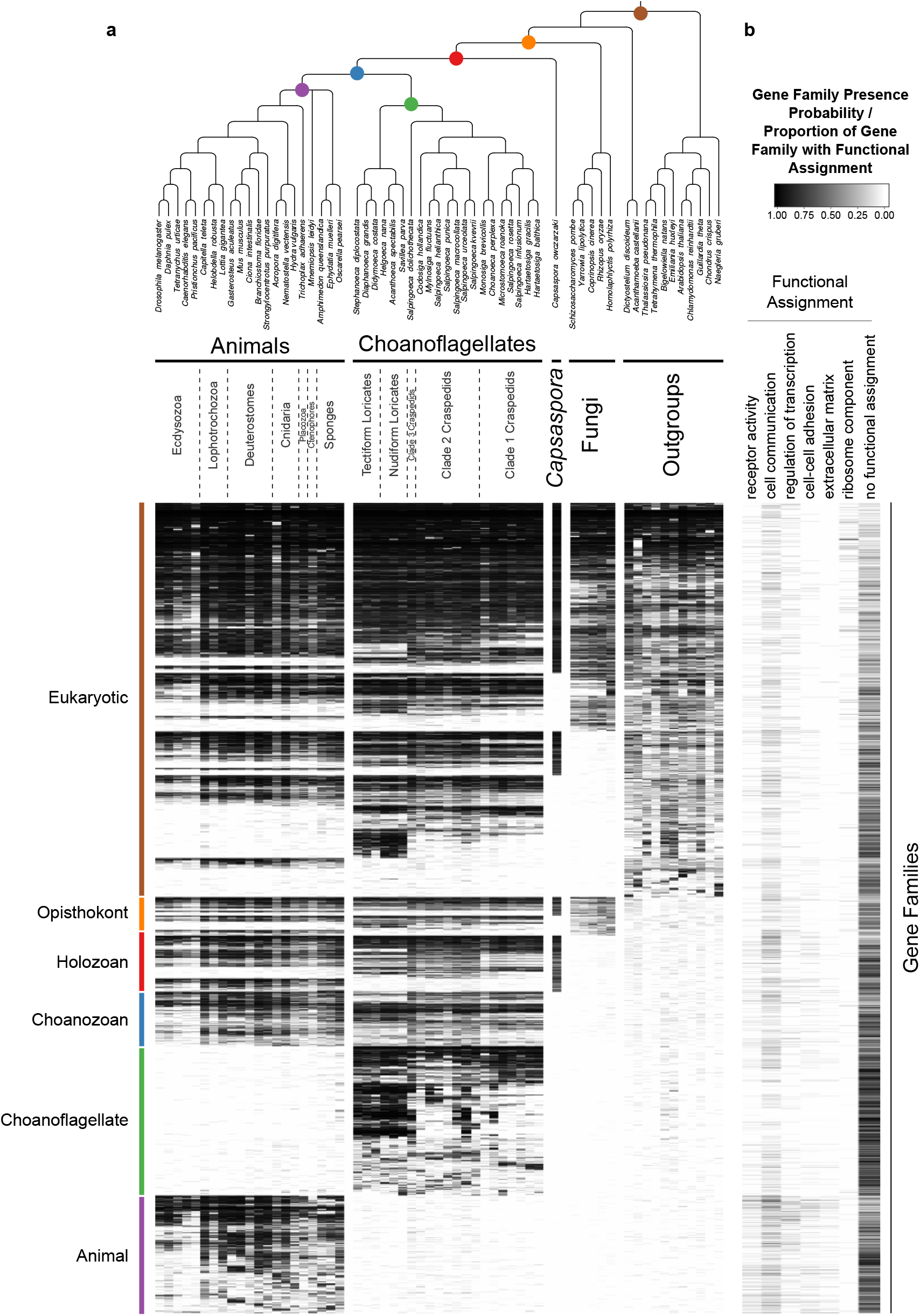
Functional assignments for gene families found in animals, choanoflagellates and their eukaryotic relatives. (a) The heat map shown in Figure 2. (b) Gene Ontology term assignment for gene families represented in the heat map for selected terms (receptor activity GO:0004872, cell communication GO:0007154, regulation of gene expression GO:0010468, cell adhesion GO:0007155, extracellular matrix GO:0031012, structural constituent of ribosome GO:0003735).

**Supplementary Figure 12.**
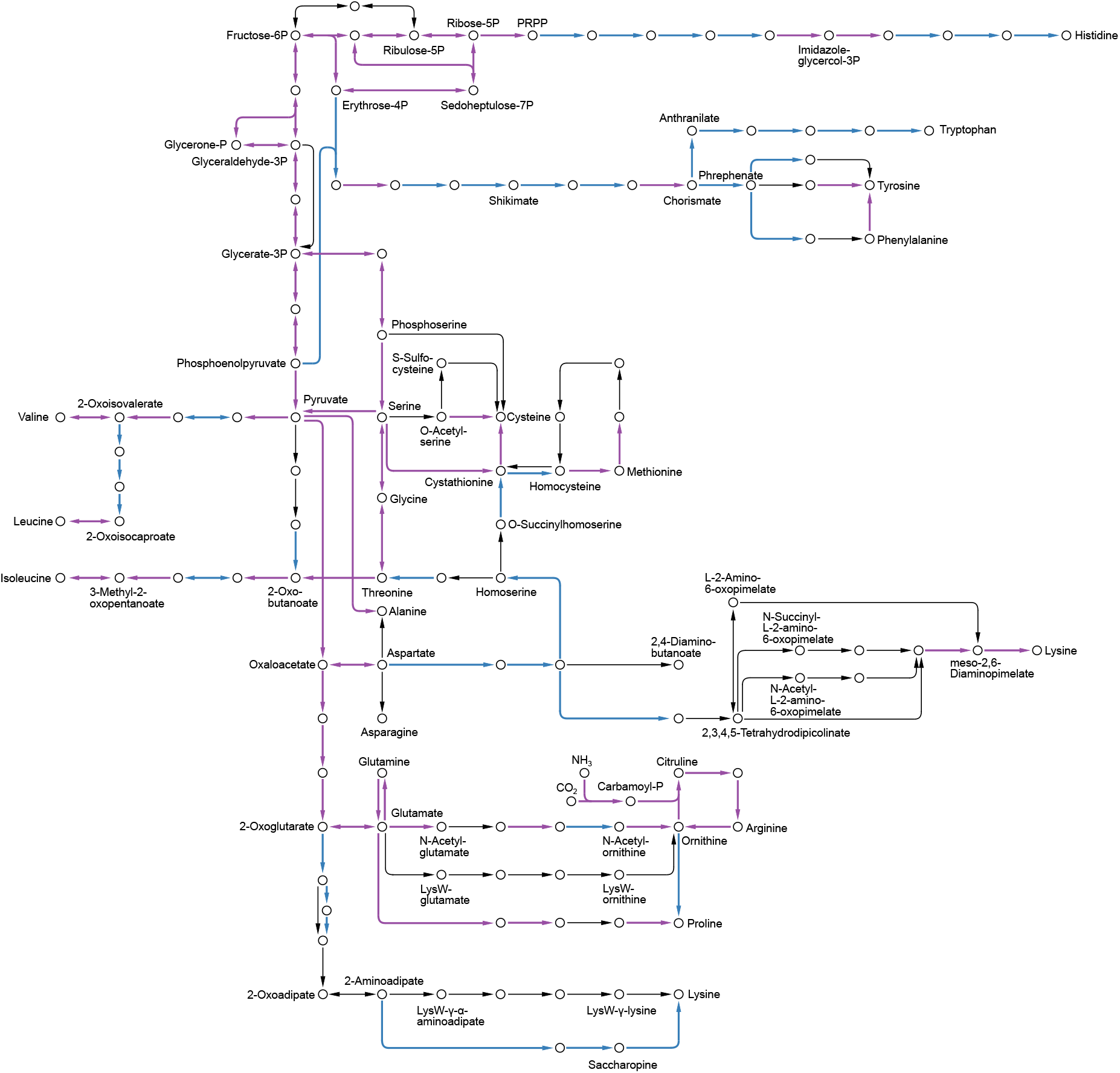
Pathway components necessary for the synthesis of the essential amino acids were lost in animals. These include the pathways for leucine, valine, isoleucine, histidine, lysine, threonine, tryptophan, phenylalanine and methionine, which were all more complete in the Urchoanozoan than in the Urmetazoan. Purple: pathway components present in the Urchoanozoan and the Urmetazoan. Blue: pathway components present in the Urchoanozoan but absent from the Urmetazoan.

**Supplementary Figure 13.**
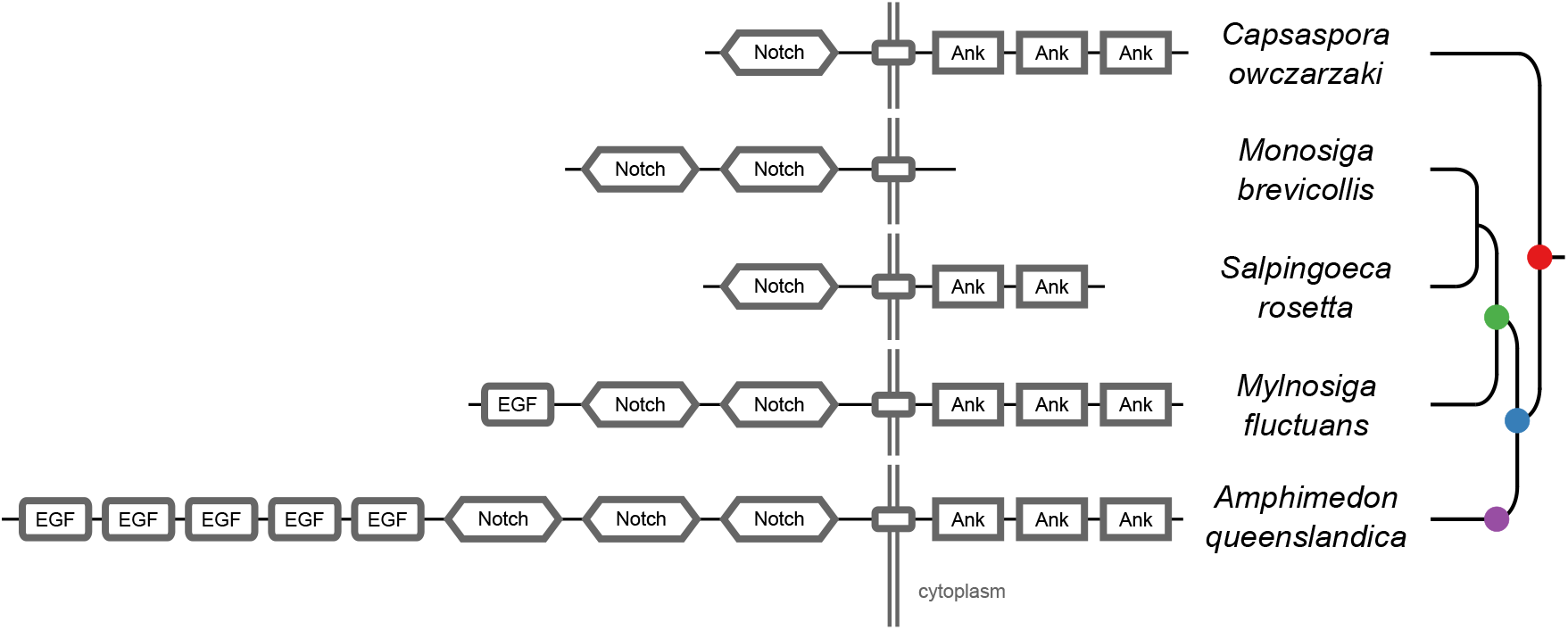
Protein domain architectures of Notch-domain containing proteins. Architectures are shown for the proteins within each species that most closely match the animal Notch protein: *C. owczarzaki* (CAOG_00333), three choanoflagellates, *M. brevicollis* (26647), *S. rosetta* (PTSG_03468), *M. fluctuans* (m.79158), and the sponge *A. queenslandica* (Aqu1.224719). In the *M. brevicollis* protein, Stealth (DUF3184) domains are not depicted. In the *A. queenslandica* protein, the EGF domain variants hEGF and EGF_CA are shown as EGF. Positions of domains in proteins are relative; they reflect ordering, but are not to scale. All protein domains are shown in grey, as they are all of ancient origin. Transmembrane domains are smaller rectangles not labeled with text.

**Supplementary Figure 14.**
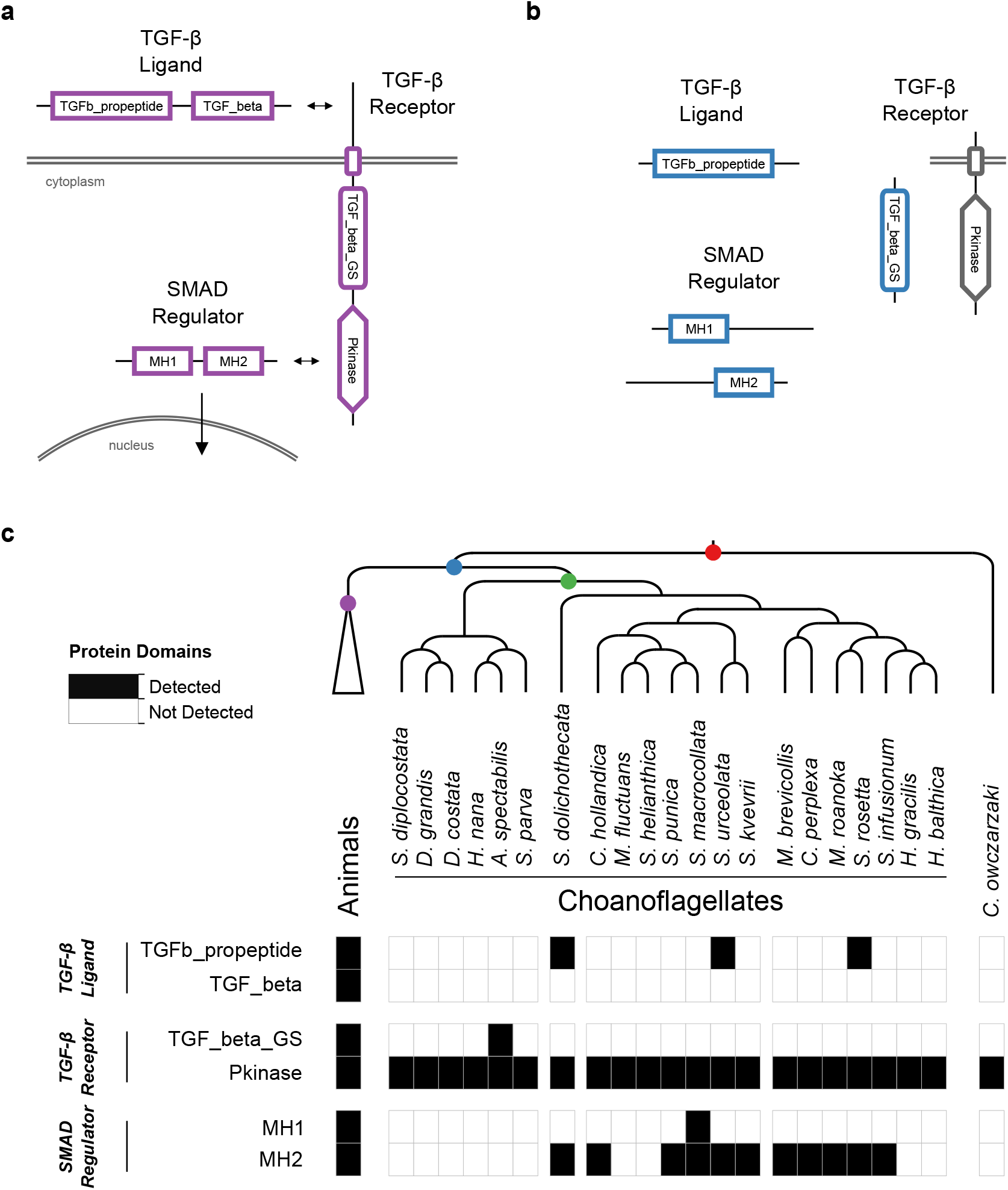
The evolution of the TGF-β signaling pathway in choanozoans. (a) In the canonical animal TGF-β signaling pathway (in purple), a TGF-β ligand (e.g. BMP, Activin) is cleaved from its TGFb_propeptide (LAP) domain, freeing it to interact with TGF-β receptors. TGF-β receptors form a heterocomplex that leads to the phosphorylation of TGF_beta_GS-containing receptors on their TGF_beta_GS domain. They subsequently phosphorylate SMADs, which translocate to the nucleus in their own oligomeric complexes. (b) Several TGF-β pathway constituent domains evolved on the Urchoanozoan stem lineage (in blue), but none of those domains are found in domain architectures of the animal pathway. (c) Presence of TGF-β constituent domains in animals, choanoflagellates and *Capsaspora owczarzaki*. The TGFb_propeptide, TGF_beta_GS, MH1 and MH2 domains are found only in choanozoans, and the TGF_beta domain is found only in animals.

**Supplementary Figure 15.**
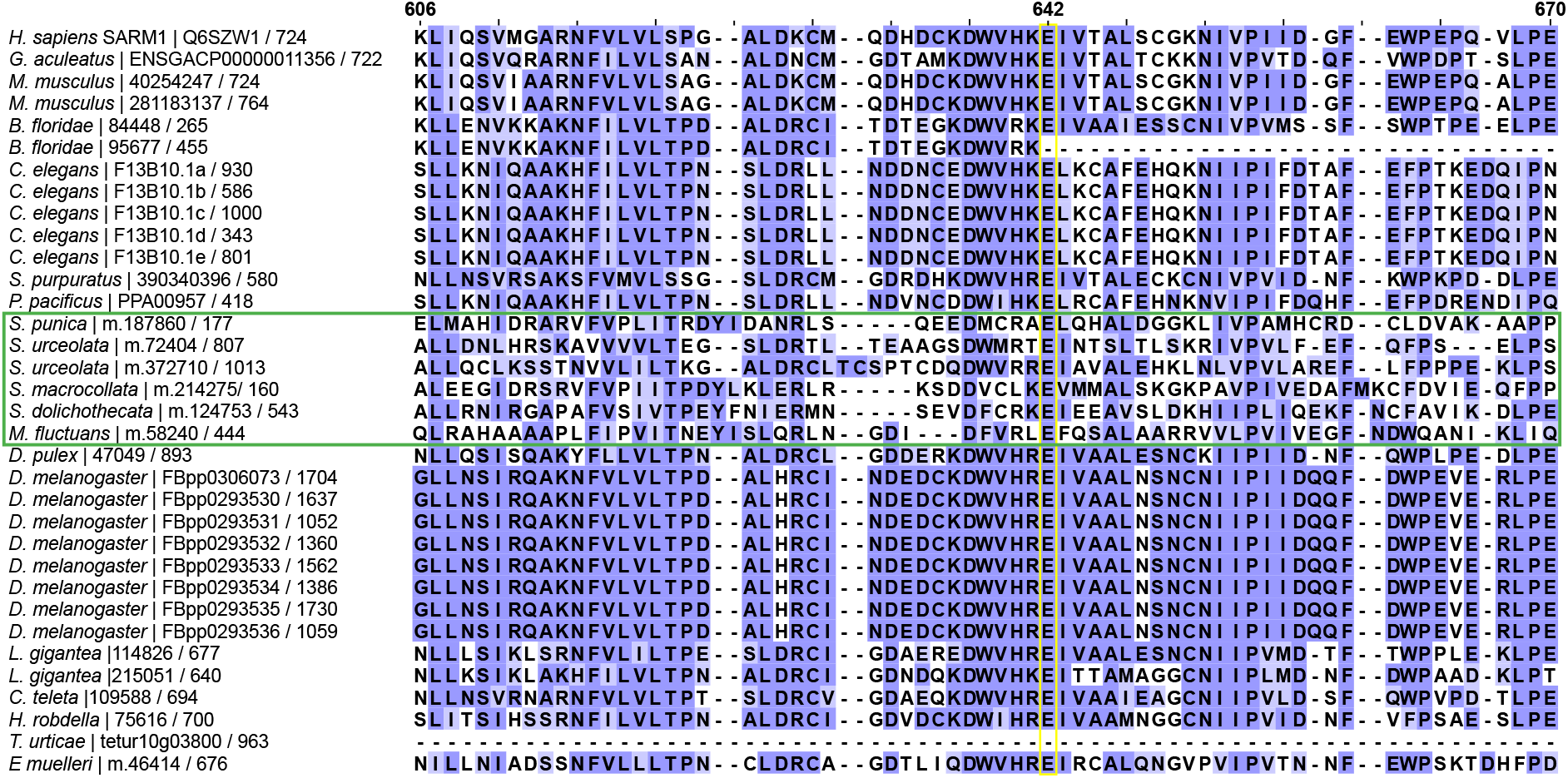
Alignment of gene family 6840, which contains animal SARM1 proteins. Only a portion of the alignment, surrounding the glutamic acid residue that is necessary but not sufficient for SARM1 function (yellow box), is depicted. The *Homo sapiens* SARM1 protein, which was not part of our data set of gene families, but which we included in the alignment, is shown at the top. Choanoflagellate sequences are surrounded by a green box. Positions at the top are given with respect to the human SARM1 protein. Each species is listed with its identifier followed by the length of the protein that was used to build the full alignment.

**Supplementary Figure 16.**
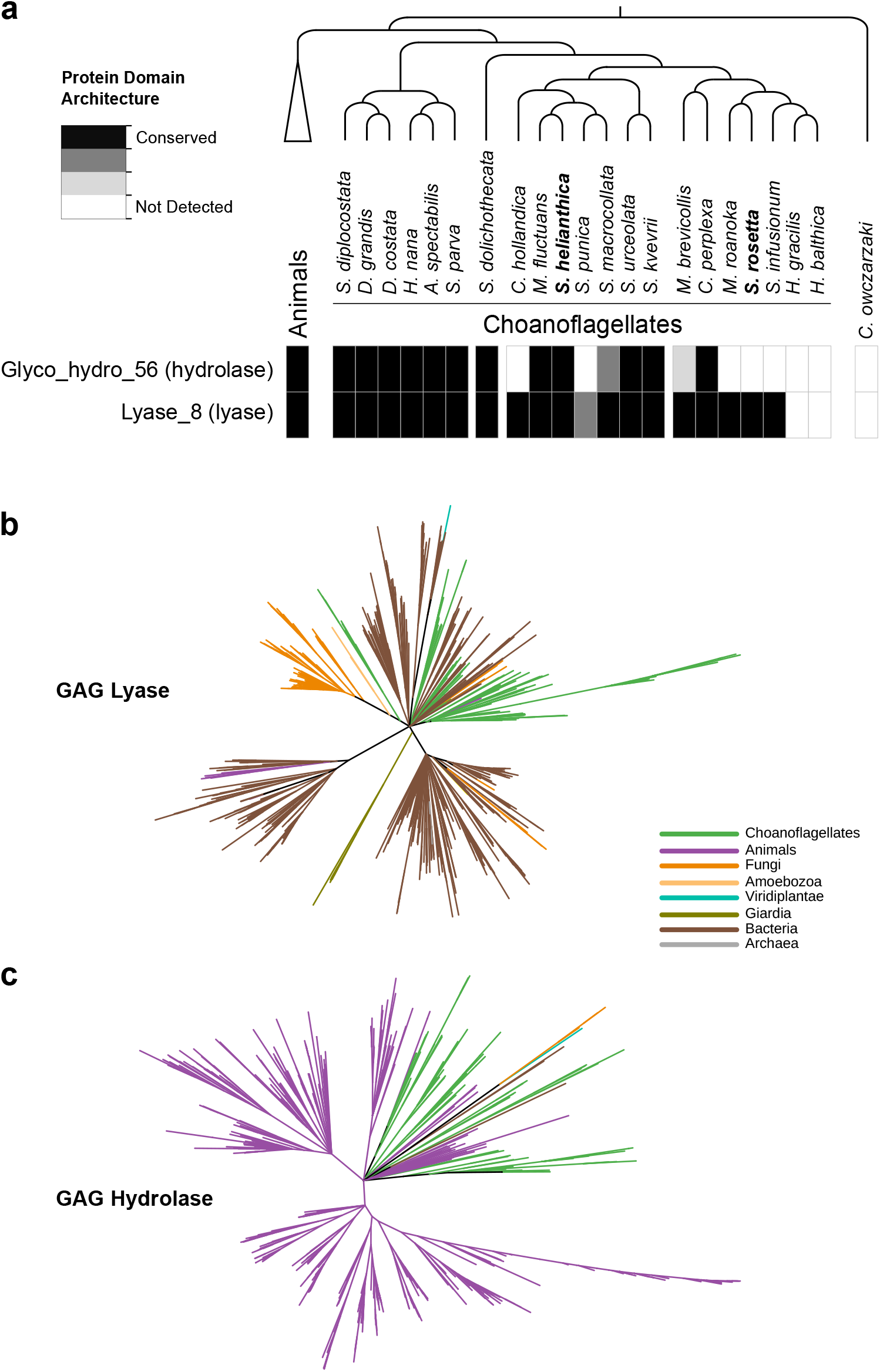
GAG cleavage genes in eukaryotes. (a) Presence of GAG hydrolase and GAG lyase in animals, choanoflagellates, and *Capsaspora owczarzaki*. Criteria are based on the presence of diagnostic protein domains (Supp. Table 6). (b) Phylogenetic tree of GAG lyases identified by the Pfam domain Lyase_8, with branches below bootstrap support of 50% removed. The animal sequences in the tree come only from sponges and cnidarians. (c) Phylogenetic tree of GAG hydrolases identified by the Pfam domain Glyco_hydro_56, with branches below bootstrap support of 50% removed. The animal sequences in the tree come from across animal diversity.

**Supplementary Figure 17.**
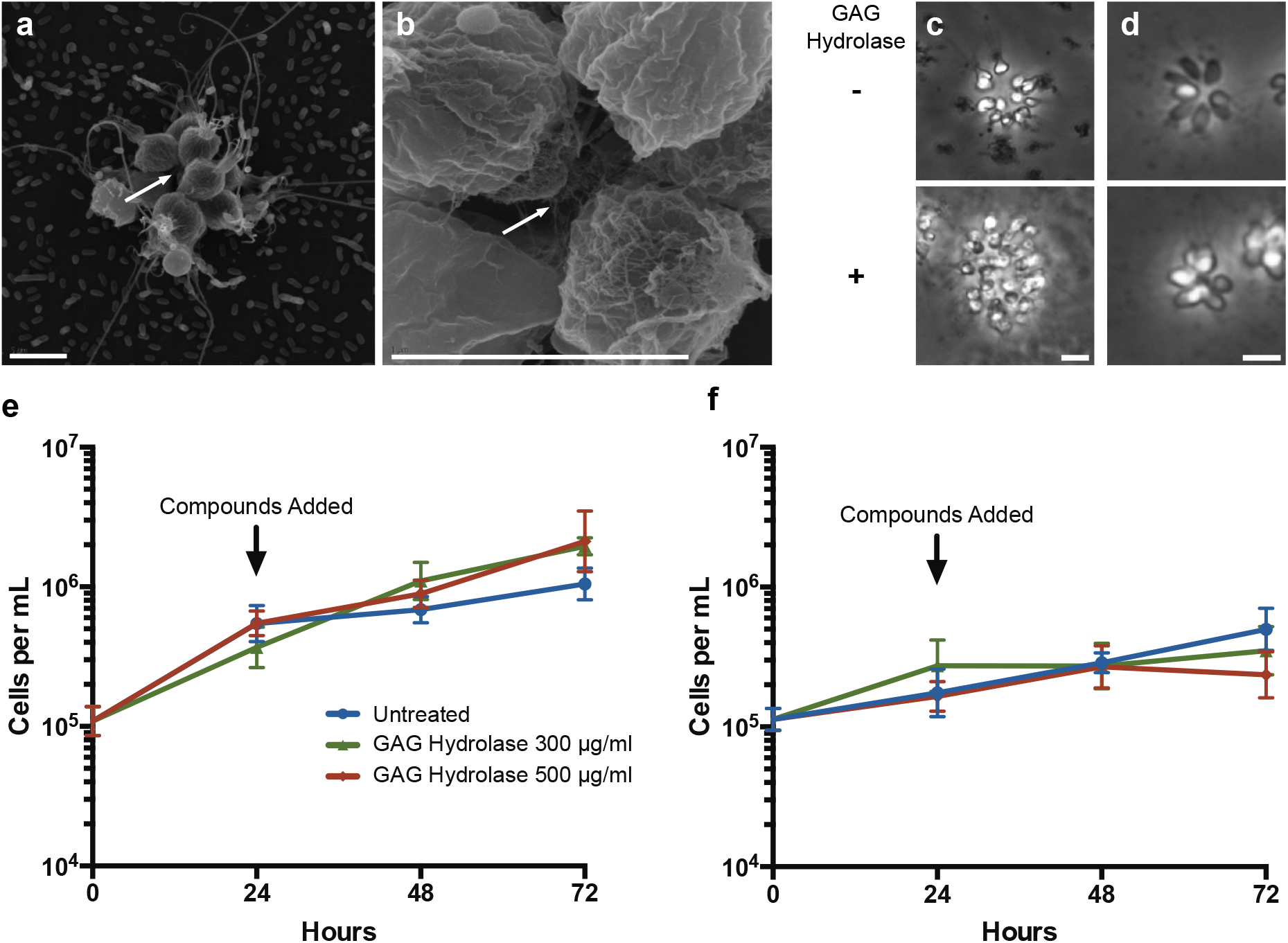
Glycosaminoglycan (GAG) biology in choanoflagellates. (a, b) SEM images of two different *S. helianthica* colonies, showing extracellular matrix between cells within the colony (white arrows). Scale bars in (b) and (c) = 5 *μ*m. (c) Phase contrast images of *S. helianthica* after no treatment or treatment with 500 *μ*g/ml bovine GAG hydrolase demonstrate that GAG hydrolase treatment leads to an increase in colony size. (d) *S. rosetta* does not show a similar response to the same treatments. Scale bars in (c) and (d) = 10 *μ*m. (e, f) Cell proliferation following treatment with GAG hydrolase in (e) *S. helianthica* and (f) *S. rosetta*. N = 8 counts per data point. Error bars represent standard deviation. p values for *S. helianthica* treatments different from untreated, Welch’s unequal variances t-test: 300 *μ*g/ul, 48 hours, p = 0.02; 300 *μ*g/ul, 72 hours, p < 0.0001; 500 *μ*g/ul, 48 hours, p = 0.14; 500 *μ*g/ul, 72 hours, p = 0.07.

**Supplementary Figure 18.**
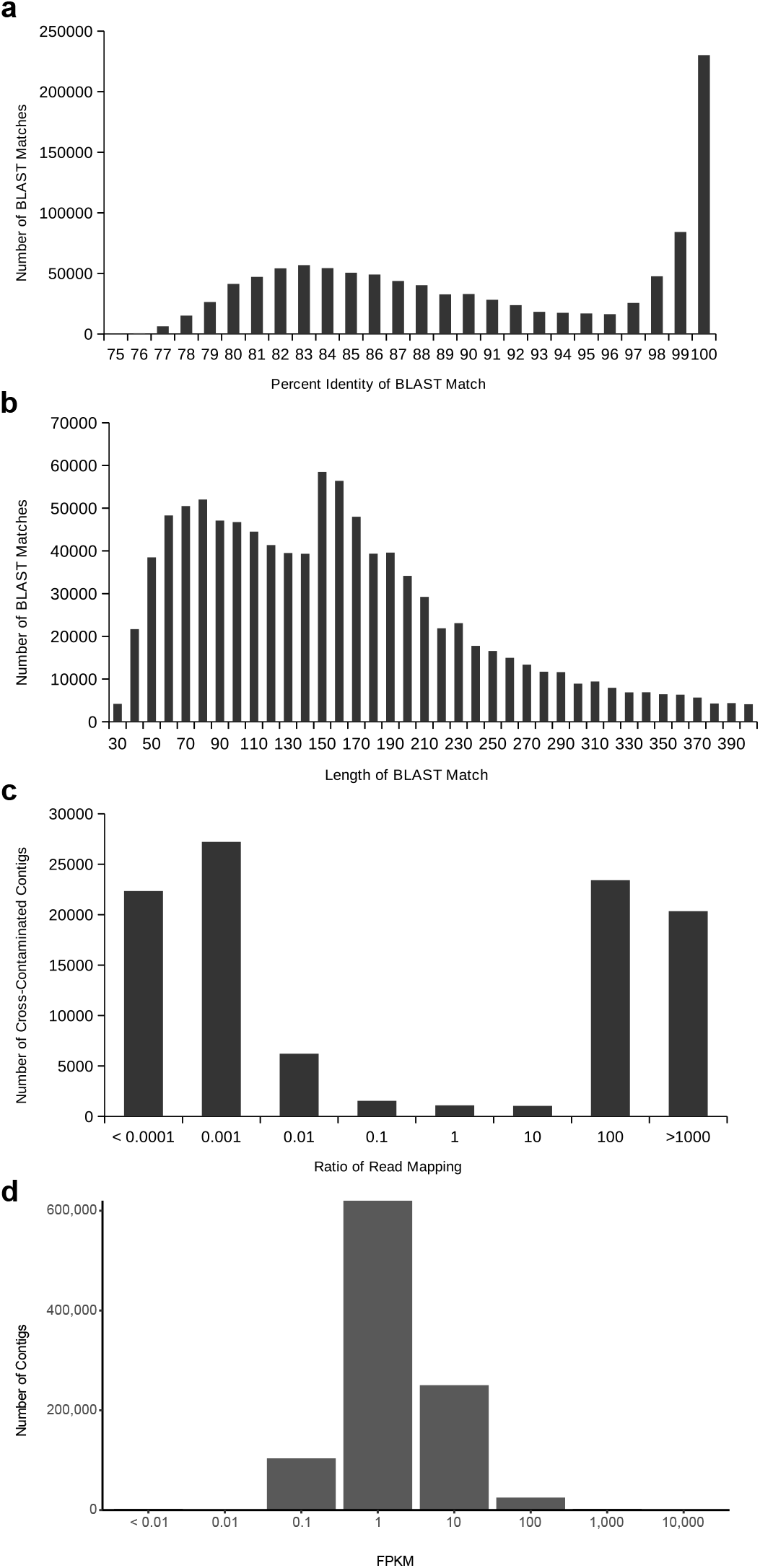
Distributions to establish thresholds within the cross-contamination removal process (a-c) and to eliminate noise contigs by FPKM (d). (a) The distribution of BLAST percent identities between contigs of different species. On the right, towards 100%, are cross-contaminants occurring on the flow cell due to library index misreading. On the left, around roughly 83%, a broader distribution from conserved homologous sequences. (b) The distribution of BLAST match lengths between contigs of different species. The peak at 150 is due to the minimum contig length of transcriptome assembly. (c) The distribution of read mapping ratios between pairs of contigs that meet the BLAST criteria to be considered putatively cross-contaminated, on a log_10_ scale. The vast majority of contigs can be easily identified as coming from one species of the pair based on read mapping ratio. (d) The distribution of FPKM values for all contigs in all transcriptome assemblies (after removal of cross-contamination), on a log_10_ scale.

**Supplementary Figure 19.**
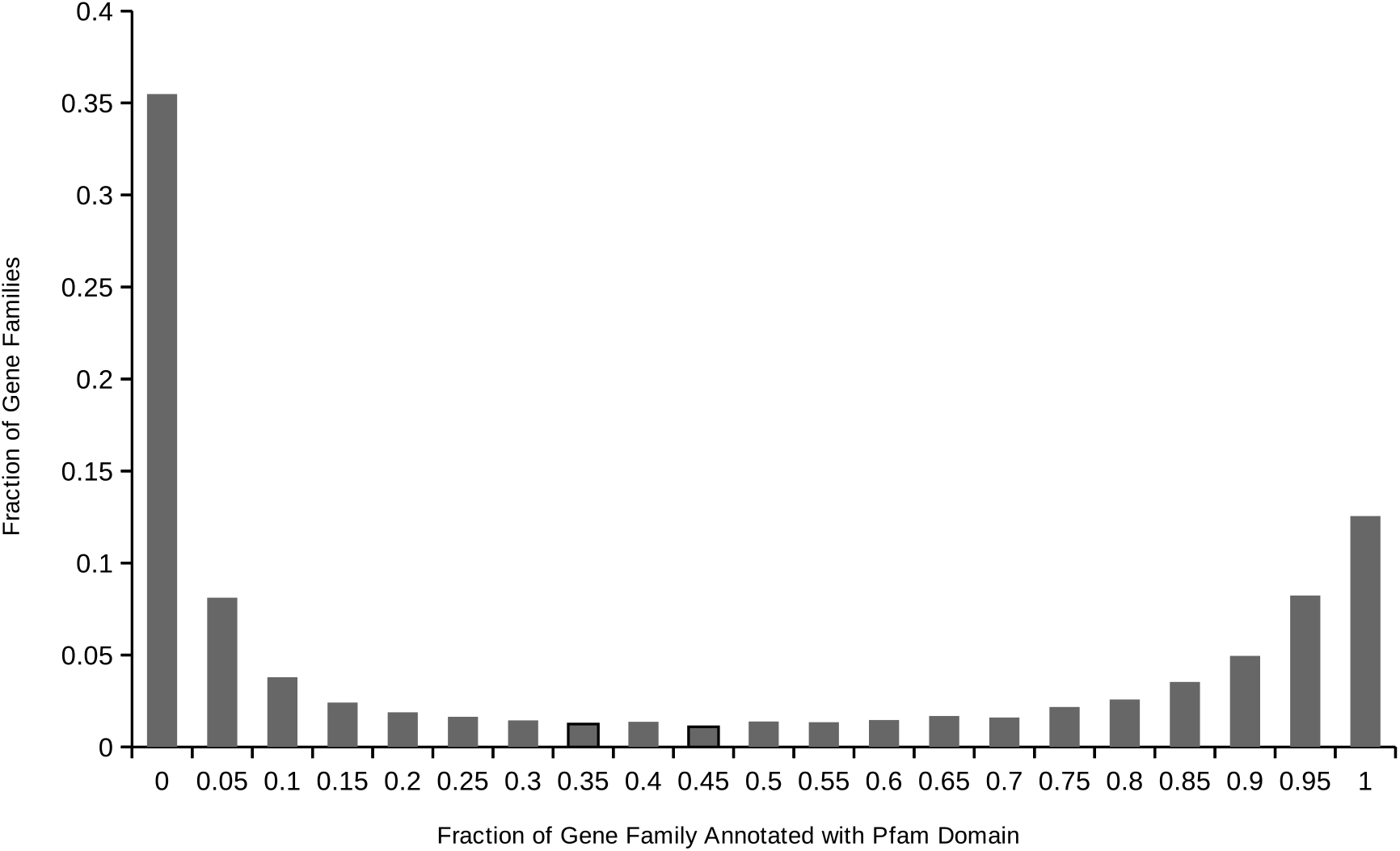
Distribution of gene family Pfam domain annotations within animals. For each gene family containing one or more proteins annotated with at least one domain, the distribution of the proportion of the proteins in the gene family annotated with each domain. We used this distribution to produce a threshold for a true vs. “noise” domain annotation of the family.

### Supplementary Table Legends

**Supplementary Table 1.** Information on each culture sequenced in this study, divided into sections by topic. See Methods for details on each topic. Note that we sequenced and assembled two strains of *Stephanoeca diplocostata*. We determined their protein catalogs to be equivalent for the purposes of gene family construction, and so we used only one of the two cultures to represent the species (7.2, ATCC 50456 isolated from France). **Strain information**: information about the cultures used, including American Type Culture Collection (ATCC) number for each culture and NCBI Taxonomy ID for each species. Previous names for cultures indicate prior names used as labels in culture collections or publications, but which were subsequently determined to have been incorrectly applied [some of these species names are no longer valid *(Codosiga gracilis, Acanthoecopsis unguiculata, Diplotheca costata, Savillea micropora*, *Monosiga gracilis* and *Monosiga ovata*), whereas others are still valid at the time of publication, but their descriptions did not match the cultures *(Salpingoeca amphoridium, Salpingoeca minuta, Salpingoeca pyxidium, Salpingoeca gracilis* and *Salpingoeca napiformis)]*. Previous names for species were never applied to the cultures used, but are considered to be invalid names that were previously used for the species and have been subsequently replaced. **Origin of cultures**: source, isolation location and year, if known. **Growth media, antibiotic treatments and cell types**: the sequence of antibiotic treatments used to obtain a culture for high-volume growth, the growth medium and temperature, and the estimated proportion of each cell type in sequenced cultures. **Culture conditions in large batches for harvesting**: information on how each culture was grown and harvested for mRNA sequencing. **Amount of total RNA used for each sample and groupings for preparation and sequencing**: the amount of total RNA used for each culture, based on the estimated proportion of choanoflagellate RNA versus bacterial RNA in each total RNA extraction, with the goal of beginning each sequencing preparation with 2 *μ*g of choanoflagellate total RNA. Library prep. group and Seq. flow cell group are arbitrary labels to indicate which sets of samples were prepared for sequencing at the same time or sequenced on the same flow cell. **Read counts and quality trimming**: results of sequencing before quality trimming and after quality trimming. **Read error correction**: parameters for error correction by Reptile, adapted to each set of reads independently. **Assembly read counts and N50s**: the counts of contigs and predicted proteins at each step of the assembly process. N50 is the length at which 50% of the nucleotides/amino acids in the assembly are contained in contigs/proteins greater than or equal to that length. **NCBI Short Read Archive**: identifiers to retrieve the raw, unprocessed reads for each library at the NCBI Short Read Archive.

**Supplementary Table 2.** Results of running BUSCO to search for conserved eukaryotic genes in each species’ protein catalog. Each value represents a percentage of genes in the BUSCO eukaryotic gene set (eukaryota_odb9).

**Supplementary Table 3.** Species used for gene family construction and their data sources. Note that we sequenced and assembled two strains of *Stephanoeca diplocostata*. We determined their protein catalogs to be equivalent for the purposes of gene family construction, and so we used only one of the two cultures to represent the species (ATCC 50456 isolated from France).

**Supplementary Table 4.** Parsimony-based rules used to determine the presence of gene families in last common ancestors of interest and whether they represent gains on that stem lineage. Gene family presences in the five major groups (Outgroups, Fungi, Filasterea (represented by *Capsaspora owczarzaki)*, Choanoflagellates and Animals) are each based on the 10% average probability threshold, as described in the Methods. The same rules are presented in two alternative formats. (a) Condensed explanation, describing the criteria for presence and gain in each last common ancestor. (b) Expanded explanation, in which a 0 represents absence and a 1 represents presence according to the 10% average probability threshold. Empty cells represent 0 and are omitted for clarity.

**Supplementary Table 5.** Counts of gene family presence, gain and loss in last common ancestors of interest. Gains and losses are not shown for the Ureukaryote, as our data set only contained eukaryotic species and was thus not appropriate to quantify changes occurring on the eukaryotic stem lineage.

**Supplementary Table 6.** Evidence levels used to determine the presence of gene families of interest. Evidence takes the form either of a protein domain architecture, the presence of a representative protein in a gene family, or a combination of the two. Protein domain names are from Pfam (transmembrane domains are from Phobius). When a domain is listed without a suffix consisting of an underscore followed by a digit, then any possible digit is acceptable (e.g., if LRR is listed, LRR_1, LRR_2, LRR_3, etc. are all acceptable). Commas indicate domain combinations, in order. Brackets are used to group combinations in order to avoid ambiguity when multiple possibilities constitute evidence for presence. Gene family IDs are from our OrthoMCL analysis and when listed, presence of a protein within that gene family is considered evidence (with a brief description of the gene family given in parentheses). *: we did not detect a canonical Toll-like receptor in *Nematostella vectensis* based on our analysis of Pfam domains in the predicted proteins from the genome assembly, but the presence of a canonical TLR has been reported in two other analyses of based on different data sets (Miller et al., 2007; Sullivan et al., 2007). **: we did not detect NF-*x*B in *Capsaspora owczarzaki* based on our analysis of Pfam domains in the predicted proteins from the genome assembly, but it has been previously reported to be present (Sebé-Pedrós et al., 2011).

**Supplementary Table 7.** Gene ontology (GO) terms enriched in the set of animal-specific gene families relative to those present in the Ureukaryote, ordered by ascending p value. The p values listed are following a Bonferroni correction for multiple testing. Only terms with an adjusted p value ≤ 0.05 are shown.

**Supplementary Table 8.** List of the 57 choanoflagellate-specific gene families that are present in all choanoflagellates in this study. Each feature (Panther, Pfam, Transmembrane, Signal Peptide, GO terms) is preceded by the fraction of proteins in the gene family annotated with the feature. Multiple annotations are separated by semicolons. The column “Pfam Kinase / Phosphatase / SH2 / SH3” indicates gene families that contain Pfam domains related to kinase signaling; the value shown represents the kinase-related Pfam domain with the maximum fraction annotated within the gene family.

**Supplementary Table 9.** Results of a MAPLE analysis comparing the gene family content of the Urchoanozoan to the Urmetazoan, to determine gains and losses on the animal stem lineage. Only those differing in completeness by at least 25% are shown. Classifications and names are from MAPLE.

**Supplementary Table 10.** Results of a MAPLE analysis comparing the gene family content of the Urholozoan to the Urchoanozoan, to determine gains and losses on the choanozoan stem lineage. Only those differing in completeness by at least 25% are shown. Classifications and names are from MAPLE.

**Supplementary Table 11.** An updated table from Woznica *et al*. (2017), with columns added for the choanoflagellate *Salpingoeca helianthica*, and for the mouse *M. musculus*, the ctenophore *M. leidyi* and the sponge *O. carmela*. In addition, there are two columns for the *Homo sapiens* genes used as queries to search the genomes of other species. The Woznica *et al*. identifier represents the one originally used in their publication, and the standardized identifier represents the identical set of underlying genes, but with all identifiers matching those found in the NCBI release GRCh38 of the *H. sapiens* genome.

**Supplementary Table 12.** Details on antibiotics tested to reduce bacterial diversity and abundance in different choanoflagellate cultures.

**Supplementary Table 13.** Characteristics of growth media used. n.d.: not determined.

**Supplementary Table 14.** Representative signaling genes in the animal TLR-NF*x*B pathway with their representatives in mouse *(Mus musculus)*. Functions and roles are derived from the UniProt database. Pfam domain architectures are separated by tildes (~) and are listed in order. When two architectures are possible, they are separated by “or”. Pfam domain architecture phylogenetic distribution represents the phylogenetic group containing all of the species with that architecture in the Pfam database. Similarly, gene family phylogenetic distribution represents the phylogenetic group containing all of the species with representative proteins in the gene family. Notes contains either a description of the species/genes present in the gene family, or a brief summary of the blastp hits in the NCBI nr database for proteins encoded by select species in the gene family, which we used to verify the assignment of the protein to the gene family.

### Supplementary File Legends

Supplementary files are available on FigShare: https://dx.doi.org/10.6084/m9.figshare.5686984

**Supplementary File 1. Final sets of contigs from choanoflagellate transcriptome assemblies.** There is one FASTA file per sequenced choanoflagellate. We assembled contigs *de novo* with Trinity, followed by removal of cross-contamination that occurred within multiplexed Illumina sequencing lanes, removal of contigs encoding strictly redundant protein sequences, and elimination of noise contigs with extremely low (PFKM < 0.01) expression levels.

**Supplementary File 2. Final sets of proteins from choanoflagellate transcriptome assemblies.** There is one FASTA file per sequenced choanoflagellate. We assembled contigs *de novo* with Trinity, followed by removal of cross-contamination that occurred within multiplexed Illumina sequencing lanes, removal of strictly redundant protein sequences, and elimination of proteins encoded on noise contigs with extremely low (PFKM < 0.01) expression levels.

**Supplementary File 3. Expression levels of assembled choanoflagellate contigs.** Expression levels are shown in FPKM, as calculated by eXpress. Percentile expression rank is calculated separately for each choanoflagellate.

**Supplementary File 4. Protein sequences for all members of each gene family.** This includes sequences from all species within the data set (i.e., it is not limited to the choanoflagellates we sequenced).

**Supplementary File 5. Gene families, group presences, and species probabilities.** For each gene family, the protein members are listed. Subsequent columns contain inferred gene family presences in different groups of species, followed by probabilities of presence in individual species in the data set.

**Supplementary File 6. List of gene families present, gained and lost in last common ancestors of interest.** A value of 1 indicates that the gene family was present, gained or lost; a value of 0 indicates that it was not. The six last common ancestors are: Ureukaryote, Uropisthokont, Urholozoan, Urchoanozoan, Urchoanoflagellate and Urmetazoan. Gains and losses are not shown for the Ureukaryote, as our data set only contained eukaryote species and was thus not appropriate to quantify changes occurring on the eukaryotic stem lineage.

**Supplementary File 7. Pfam, transmembrane, signal peptide, PANTHER and Gene Ontology annotations for all proteins.** Annotations are listed for all proteins in the data set, including those not part of any gene family. Pfam domains are delimited by a tilde (~) and Gene Ontology terms by a semicolon (;). Transmembrane domains and signal peptides are indicated by the number present in the protein, followed by their coordinates in the protein sequence.

**Supplementary File 8. Pfam, transmembrane, signal peptide, PANTHER and Gene Ontology annotations aggregated by gene family.** The proportion of proteins within the gene family that were assigned an annotation is followed by the name of the annotation. Multiple annotations are delimited by a semicolon (;).

## References

Abedin, M., and King, N. (2008). The premetazoan ancestry of cadherins. Science, 319(5865), 946–948. https://doi.org/10.1126/science.1151084

Abedin, M., and King, N. (2010). Diverse evolutionary paths to cell adhesion. Trends in Cell Biology, 20(12), 734–742. https://doi.org/10.1016/j.tcb.2010.08.002

Adams, J. C., and Watt, F. M. (1993). Regulation of development and differentiation by the extracellular matrix. Development, 117(4), 1183–98. http://www.ncbi.nlm.nih.gov/pubmed/8404525

Adamska, M., Matus, D. Q., Adamski, M., Green, K., Rokhsar, D. S., Martindale, M. Q., and Degnan, B. M. (2007). The evolutionary origin of hedgehog proteins. Current Biology, 17(19), R836–R837. https://doi.org/10.1016/j.cub.2007.08.010

Albalat, R., and Cañestro, C. (2016). Evolution by gene loss. Nature Reviews Genetics, 17(7), 379–391. https://doi.org/10.1038/nrg.2016.39

Alegado, R. A., Brown, L. W., Cao, S., Dermenjian, R. K., Zuzow, R., Fairclough, S. R.,…King, N. (2012). A bacterial sulfonolipid triggers multicellular development in the closest living relatives of animals. eLife, 1, e00013. https://doi.org/10.7554/eLife.00013

Alegado, R. A., Ferriera, S., Nusbaum, C., Young, S. K., Zeng, Q., Imamovic, A.,…King, N. (2011). Complete Genome Sequence of Algoriphagus sp. PR1, Bacterial Prey of a Colony-Forming Choanoflagellate. Journal of Bacteriology, 193(6), 1485–1486. https://doi.org/10.1128/JB.01421-10

Alilain, W. J., Horn, K. P., Hu, H., Dick, T. E., and Silver, J. (2011). Functional regeneration of respiratory pathways after spinal cord injury. Nature, 475(7355), 196–200. https://doi.org/10.1038/nature10199

Altenhoff, A. M., Boeckmann, B., Capella-Gutierrez, S., Dalquen, D. A., DeLuca, T., Forslund, K.,…Dessimoz, C. (2016). Standardized benchmarking in the quest for orthologs. Nature Methods, 13(5), 425–30. https://doi.org/10.1038/nmeth.3830

Altschul, S. (1997). Gapped BLAST and PSI-BLAST: a new generation of protein database search programs. Nucleic Acids Research, 25(17), 3389–3402. https://doi.org/10.1093/nar/25.17.3389

Aravin, A. A., Hannon, G. J., and Brennecke, J. (2007). The Piwi-piRNA pathway provides an adaptive defense in the transposon arms race. Science, 318(5851), 761–4. https://doi.org/10.1126/science.1146484

Aravin, A. A., Naumova, N. M., Tulin, A. V, Vagin, V. V, Rozovsky, Y. M., and Gvozdev, V. A. (2001). Double-stranded RNA-mediated silencing of genomic tandem repeats and transposable elements in the D. melanogaster germline. Current Biology, 11(13), 1017–1027. https://doi.org/10.1016/S0960-9822(01)00299-8

Artavanis-Tsakonas, S. (1999). Notch Signaling: Cell Fate Control and Signal Integration in Development. Science, 284(5415), 770–776. https://doi.org/10.1126/science.284.5415.770

Ashburner, M., Ball, C. A., Blake, J. A., Botstein, D., Butler, H., Cherry, J. M.,…Sherlock, G. (2000). Gene ontology: tool for the unification of biology. The Gene Ontology Consortium. Nature Genetics, 25(1), 25–29. https://doi.org/10.1038/75556

Aspöck, G., Kagoshima, H., Niklaus, G., and Bürglin, T. R. (1999). Caenorhabditis elegans Has Scores of hedgehog Related Genes: Sequence and Expression Analysis. Genome Research, 9(10), 909–923. https://doi.org/10.1101/gr.9.10.909

Ausubel, F. M. (2005). Are innate immune signaling pathways in plants and animals conserved? Nature Immunology, 6(10), 973–979. https://doi.org/10.1038/ni1253

Babonis, L. S., and Martindale, M. Q. (2017). Phylogenetic evidence for the modular evolution of metazoan signalling pathways. Philosophical Transactions of the Royal Society B: Biological Sciences, 372(1713), 20150477. https://doi.org/10.1098/rstb.2015.0477

Benson, G. (1999). Tandem repeats finder: a program to analyze DNA sequences. Nucleic Acids Research, 27(2), 573–580. https://doi.org/10.1093/nar/27.2.573

Bonnans, C., Chou, J., and Werb, Z. (2014). Remodelling the extracellular matrix in development and disease. Nature Reviews Molecular Cell Biology, 15(12), 786–801. https://doi.org/10.1038/nrm3904

Bonner, J. T. (1998). The origins of multicellularity. Integrative Biology: Issues, News, and Reviews, 1(1), 27–36. https://doi.org/10.1002/(SICI)1520-6602(1998)1:1<27::AID-INBI4>3.0.CO;2-6

Bouvard, D., Vignoud, L., Dupé-Manet, S., Abed, N., Fournier, H.-N., Vincent-Monegat, C.,…Block, M. R. (2003). Disruption of Focal Adhesions by Integrin Cytoplasmic Domain-associated Protein-1α. Journal of Biological Chemistry, 278(8), 6567–6574. https://doi.org/10.1074/jbc.M211258200

Brennan, J. J., Messerschmidt, J. L., Williams, L. M., Matthews, B. J., Reynoso, M., and Gilmore, T. D. (2017). Sea anemone model has a single Toll-like receptor that can function in pathogen detection, NF-*x*B signal transduction, and development. Proceedings of the National Academy of Sciences, 114(47), E10122–E10131. https://doi.org/10.1073/pnas.1711530114

Brunet, T., and King, N. (2017). The Origin of Animal Multicellularity and Cell Differentiation. Developmental Cell, 43(2), 124–140. https://doi.org/10.1016/j.devcel.2017.09.016

Burki, F., Kaplan, M., Tikhonenkov, D. V., Zlatogursky, V., Minh, B. Q., Radaykina, L. V.,…Keeling, P. J. (2016). Untangling the early diversification of eukaryotes: a phylogenomic study of the evolutionary origins of Centrohelida, Haptophyta and Cryptista. Proceedings of the Royal Society B: Biological Sciences, 283(1823), 20152802. https://doi.org/10.1098/rspb.2015.2802

Busch, S. A., and Silver, J. (2007). The role of extracellular matrix in CNS regeneration. Current Opinion in Neurobiology, 17(1), 120–127. https://doi.org/10.1016/j.conb.2006.09.004

Buss, L. W. (1987). The evolution of individuality. Princeton, New Jersey: Princeton University Press.

C. elegans Sequencing Consortium, T. (1998). Genome Sequence of the Nematode C. elegans: A Platform for Investigating Biology. Science, 282(5396), 2012–2018. https://doi.org/10.1126/science.282.5396.2012

Capella-Gutierrez, S., Silla-Martinez, J. M., and Gabaldon, T. (2009). trimAl: a tool for automated alignment trimming in large-scale phylogenetic analyses. Bioinformatics, 25(15), 1972–1973. https://doi.org/10.1093/bioinformatics/btp348

Carr, M., Leadbeater, B. S. C., Hassan, R., Nelson, M., and Baldauf, S. L. (2008a). Molecular phylogeny of choanoflagellates, the sister group to Metazoa. Proceedings of the National Academy of Sciences, 105(43), 16641–16646. https://doi.org/10.1073/pnas.0801667105

Carr, M., Nelson, M., Leadbeater, B. S. C., and Baldauf, S. L. (2008b). Three families of LTR retrotransposons are present in the genome of the choanoflagellate Monosiga brevicollis. Protist, 159(4), 579–590. https://doi.org/10.1016/j.protis.2008.05.001

Carr, M., Richter, D. J., Fozouni, P., Smith, T. J., Jeuck, A., Leadbeater, B. S. C., and Nitsche, F. (2017). A six-gene phylogeny provides new insights into choanoflagellate evolution. Molecular Phylogenetics and Evolution, 107, 166–178. https://doi.org/10.1016/j.ympev.2016.10.011

Chae, J., Kim, M. J., Goo, J. H., Collier, S., Gubb, D., Charlton, J.,…Park, W. J. (1999). The Drosophila tissue polarity gene starry night encodes a member of the protocadherin family. Development, 126(23), 5421–5429. http://www.ncbi.nlm.nih.gov/pubmed/10556066

Chang, D. D., Wong, C., Smith, H., and Liu, J. (1997). ICAP-1, a Novel β 1 Integrin Cytoplasmic Domain–associated Protein, Binds to a Conserved and Functionally Important NPXY Sequence Motif of β 1 Integrin. The Journal of Cell Biology, 138(5), 1149–1157. https://doi.org/10.1083/jcb.138.5.1149

Chang, E. S., Neuhof, M., Rubinstein, N. D., Diamant, A., Philippe, H., Huchon, D., and Cartwright, P. (2015). Genomic insights into the evolutionary origin of Myxozoa within Cnidaria. Proceedings of the National Academy of Sciences, 112(48), 14912–14917. https://doi.org/10.1073/pnas.1511468112

Cheloufi, S., Dos Santos, C. O., Chong, M. M. W., and Hannon, G. J. (2010). A dicer-independent miRNA biogenesis pathway that requires Ago catalysis. Nature, 465(7298), 584–589. https://doi.org/10.1038/nature09092

Cherr, G. N., Yudin, A. I., and Overstreet, J. W. (2001). The dual functions of GPI-anchored PH-20: hyaluronidase and intracellular signaling. Matrix Biology, 20(8), 515–525. https://doi.org/10.1016/S0945-053X(01)00171-8

Cock, J. M., Sterck, L., Rouzé, P., Scornet, D., Allen, A. E., Amoutzias, G.,…Wincker, P. (2010). The Ectocarpus genome and the independent evolution of multicellularity in brown algae. Nature, 465(7298), 617–621. https://doi.org/10.1038/nature09016

Couillault, C., Pujol, N., Reboul, J., Sabatier, L., Guichou, J.-F., Kohara, Y., and Ewbank, J. J. (2004). TLR-independent control of innate immunity in Caenorhabditis elegans by the TIR domain adaptor protein TIR-1, an ortholog of human SARM. Nature Immunology, 5(5), 488–494. https://doi.org/10.1038/ni1060

Dayel, M. J., Alegado, R. A., Fairclough, S. R., Levin, T. C., Nichols, S. A., McDonald, K., and King, N. (2011). Cell differentiation and morphogenesis in the colony-forming choanoflagellate Salpingoeca rosetta. Developmental Biology, 357(1), 73–82. https://doi.org/10.1016/j.ydbio.2011.06.003

de Hoon, M. J. L., Imoto, S., Nolan, J., and Miyano, S. (2004). Open source clustering software. Bioinformatics, 20(9), 1453–1454. https://doi.org/10.1093/bioinformatics/bth078

de Mendoza, A., Sebé-Pedrós, A., Sestak, M. S., Matejcic, M., Torruella, G., Domazet-Loso, T., and Ruiz-Trillo, I. (2013). Transcription factor evolution in eukaryotes and the assembly of the regulatory toolkit in multicellular lineages. Proceedings of the National Academy of Sciences, 110(50), E4858–E4866. https://doi.org/10.1073/pnas.1311818110

de Mendoza, A., Suga, H., Permanyer, J., Irimia, M., and Ruiz-Trillo, I. (2015). Complex transcriptional regulation and independent evolution of fungal-like traits in a relative of animals. eLife, 4. https://doi.org/10.7554/eLife.08904

DeAndrade, M. P., Zhang, L., Doroodchi, A., Yokoi, F., Cheetham, C. C., Chen, H.-X.,…Li, Y. (2012). Enhanced Hippocampal Long-Term Potentiation and Fear Memory in Btbd9 Mutant Mice. PLoS ONE, 7(4), e35518. https://doi.org/10.1371/journal.pone.0035518

Douzery, E. J. P., Snell, E. A., Bapteste, E., Delsuc, F., and Philippe, H. (2004). The timing of eukaryotic evolution: Does a relaxed molecular clock reconcile proteins and fossils? Proceedings of the National Academy of Sciences, 101(43), 15386–15391. https://doi.org/10.1073/pnas.0403984101

Drinnenberg, I. A., Weinberg, D. E., Xie, K. T., Mower, J. P., Wolfe, K. H., Fink, G. R., and Bartel, D. P. (2009). RNAi in budding yeast. Science, 326(5952), 544–550. https://doi.org/10.1126/science.1176945

Dujardin, F. (1841). Histoire naturelle des zoophytes. Infusoires: comprenant la physiologie et la classification de ces animaux et la manière de les étudier à l’aide du microscope. Paris, France: Librairie Encyclopédique de Roret.

Eddy, S. R. (2011). Accelerated Profile HMM Searches. PLoS Computational Biology, 7(10), e1002195. https://doi.org/10.1371/journal.pcbi.1002195

Ekman, D., Björklund, Å. K., and Elofsson, A. (2007). Quantification of the Elevated Rate of Domain Rearrangements in Metazoa. Journal of Molecular Biology, 372(5), 1337–1348. https://doi.org/10.1016/j.jmb.2007.06.022

Enright, A. J. (2002). An efficient algorithm for large-scale detection of protein families. Nucleic Acids Research, 30(7), 1575–1584. https://doi.org/10.1093/nar/30.7.1575

Erives, A. J., and Fassler, J. S. (2015). Metabolic and Chaperone Gene Loss Marks the Origin of Animals: Evidence for Hsp104 and Hsp78 Chaperones Sharing Mitochondrial Enzymes as Clients. PLOS ONE, 10(2), e0117192. https://doi.org/10.1371/journal.pone.0117192

Essuman, K., Summers, D. W., Sasaki, Y., Mao, X., DiAntonio, A., and Milbrandt, J. (2017). The SARM1 Toll/Interleukin-1 Receptor Domain Possesses Intrinsic NAD + Cleavage Activity that Promotes Pathological Axonal Degeneration. Neuron, 93(6), 1334–1343.e5. https://doi.org/10.1016/j.neuron.2017.02.022

Fairclough, S. R., Chen, Z., Kramer, E., Zeng, Q., Young, S., Robertson, H. M.,…King, N. (2013). Premetazoan genome evolution and the regulation of cell differentiation in the choanoflagellate Salpingoeca rosetta. Genome Biology, 14(2), R15. https://doi.org/10.1186/gb-2013-14-2-r15

Fairclough, S. R., Dayel, M. J., and King, N. (2010). Multicellular development in a choanoflagellate. Current Biology, 20(20), R875–6. https://doi.org/10.1016/j.cub.2010.09.014

Ferrer-Bonet, M., and Ruiz-Trillo, I. (2017). Capsaspora owczarzaki. Current Biology, 27(17), R829–R830. https://doi.org/10.1016/j.cub.2017.05.074

Finn, R. D., Coggill, P., Eberhardt, R. Y., Eddy, S. R., Mistry, J., Mitchell, A. L.,…Bateman, A. (2016). The Pfam protein families database: towards a more sustainable future. Nucleic Acids Research, 44(D1), D279–D285. https://doi.org/10.1093/nar/gkv1344

Fitzgerald, L. M., and Szmant, A. M. (1997). Biosynthesis of “essential” amino acids by scleractinian corals. Biochemical Journal, 322(1), 213–221. https://doi.org/10.1042/bj3220213

Fortunato, S. A. V., Adamski, M., Ramos, O. M., Leininger, S., Liu, J., Ferrier, D. E. K., and Adamska, M. (2014). Calcisponges have a ParaHox gene and dynamic expression of dispersed NK homeobox genes. Nature, 514(7524), 620–623. https://doi.org/10.1038/nature13881

Francis, W. R., Eitel, M., Vargas R., S., Adamski, M., Haddock, S. H. D., Krebs, S.,…Wörheide, G. (2017). The Genome Of The Contractile Demosponge *Tethya wilhelma* And The Evolution Of Metazoan Neural Signalling Pathways. bioRxiv. https://doi.org/10.1101/120998

Frank, M., and Kemler, R. (2002). Protocadherins. Current Opinion in Cell Biology, 14(5), 557–562. https://doi.org/10.1016/S0955-0674(02)00365-4

Gauthier, M. E. A., Du Pasquier, L., and Degnan, B. M. (2010). The genome of the sponge Amphimedon queenslandica provides new perspectives into the origin of Toll-like and interleukin 1 receptor pathways. Evolution & Development, 12(5), 519–533. https://doi.org/10.1111/j.1525-142X.2010.00436.x

Gazave, E., Lapébie, P., Richards, G. S., Brunet, F., Ereskovsky, A. V, Degnan, B. M.,…Renard, E. (2009). Origin and evolution of the Notch signalling pathway: an overview from eukaryotic genomes. BMC Evolutionary Biology, 9(1), 249. https://doi.org/10.1186/1471-2148-9-249

Giger, R. J., Hollis, E. R., and Tuszynski, M. H. (2010). Guidance Molecules in Axon Regeneration. Cold Spring Harbor Perspectives in Biology, 2(7), a001867–a001867. https://doi.org/10.1101/cshperspect.a001867

Gilmore, T. D. (1999). The Rel/NF-*x*B signal transduction pathway: introduction. Oncogene, 18(49), 6842–6844. https://doi.org/10.1038/sj.onc.1203237

Gilmore, T. D., and Wolenski, F. S. (2012). NF-*x*B: where did it come from and why? Immunological Reviews, 246(1), 14–35. https://doi.org/10.1111/j.1600-065X.2012.01096.x

Girish, K. S., and Kemparaju, K. (2007). The magic glue hyaluronan and its eraser hyaluronidase: A biological overview. Life Sciences, 80(21), 1921–1943. https://doi.org/10.1016/j.lfs.2007.02.037

Glöckner, G., Lawal, H. M., Felder, M., Singh, R., Singer, G., Weijer, C. J., and Schaap, P. (2016). The multicellularity genes of dictyostelid social amoebas. Nature Communications, 7, 12085. https://doi.org/10.1038/ncomms12085

Grabherr, M. G., Haas, B. J., Yassour, M., Levin, J. Z., Thompson, D. A., Amit, I.,…Regev, A. (2011). Full-length transcriptome assembly from RNA-Seq data without a reference genome. Nature Biotechnology, 29(7), 644–652. https://doi.org/10.1038/nbt.1883

Gramates, L. S., Marygold, S. J., Santos, G. dos, Urbano, J.-M., Antonazzo, G., Matthews, B. B.,…Zhou, P. (2017). FlyBase at 25: looking to the future. Nucleic Acids Research, 45(D1), D663–D671. https://doi.org/10.1093/nar/gkw1016

Grau-Bové, X., Sebé-Pedrós, A., and Ruiz-Trillo, I. (2015). The Eukaryotic Ancestor Had a Complex Ubiquitin Signaling System of Archaeal Origin. Molecular Biology and Evolution, 32(3), 726–739. https://doi.org/10.1093/molbev/msu334

Grau-Bové, X., Torruella, G., Donachie, S., Suga, H., Leonard, G., Richards, T. A., and Ruiz-Trillo, I. (2017). Dynamics of genomic innovation in the unicellular ancestry of animals. eLife, 6. https://doi.org/10.7554/eLife.26036

Grossmann, S., Bauer, S., Robinson, P. N., and Vingron, M. (2007). Improved detection of overrepresentation of Gene-Ontology annotations with parent child analysis. Bioinformatics, 23(22), 3024–3031. https://doi.org/10.1093/bioinformatics/btm440

Gu, Z., David, L., Petrov, D., Jones, T., Davis, R. W., and Steinmetz, L. M. (2005). Elevated evolutionary rates in the laboratory strain of Saccharomyces cerevisiae. Proceedings of the National Academy of Sciences, 102(4), 1092–1097. https://doi.org/10.1073/pnas.0409159102

Guedes, R., Prosdocimi, F., Fernandes, G., Moura, L., Ribeiro, H., and Ortega, J. (2011). Amino acids biosynthesis and nitrogen assimilation pathways: a great genomic deletion during eukaryotes evolution. BMC Genomics, 12(Suppl 4), S2. https://doi.org/10.1186/1471-2164-12-S4-S2

Haas, B. J., Papanicolaou, A., Yassour, M., Grabherr, M., Blood, P. D., Bowden, J.,…Regev, A. (2013). De novo transcript sequence reconstruction from RNA-seq using the Trinity platform for reference generation and analysis. Nature Protocols, 8(8), 1494–1512. https://doi.org/10.1038/nprot.2013.084

Haeckel, E. (1869). Uber den Organismus der Schwamme und ihre Verwandtschaft mit der Corallen. Jenaische Zeitschrift, 5, 207–254.

Haeckel, E. (1873). On the Calciospongiae, their position in the animal kingdom and their relation to the theory of descendence. The Annals and Magazine of Natural History, 4, 241–262–431.

Haeckel, E. (1874). The gastrea-theory, the phylogenetic classification of the animal kingdom and the homology of the germ-lamellae. Quarterly Journal of Microscopical Science, 14, 142–165.

Han, M. V, and Zmasek, C. M. (2009). phyloXML: XML for evolutionary biology and comparative genomics. BMC Bioinformatics, 10(1), 356. https://doi.org/10.1186/1471-2105-10-356

Harrower, M., and Brewer, C. A. (2003). ColorBrewer.org: An Online Tool for Selecting Colour Schemes for Maps. The Cartographic Journal, 40(1), 27–37. https://doi.org/10.1179/000870403235002042

Hausmann, G., von Mering, C., and Basler, K. (2009). The Hedgehog Signaling Pathway: Where Did It Come From? PLoS Biology, 7(6), e1000146. https://doi.org/10.1371/journal.pbio.1000146

Hay, E. D. (Ed.). (1991). Cell Biology of Extracellular Matrix. Boston, MA: Springer US. https://doi.org/10.1007/978-1-4615-3770-0

Hedges, S. B., Blair, J. E., Venturi, M. L., and Shoe, J. L. (2004). A molecular timescale of eukaryote evolution and the rise of complex multicellular life. BMC Evolutionary Biology, 4, 2. https://doi.org/10.1186/1471-2148-4-2

Heldin, C.-H., Miyazono, K., and ten Dijke, P. (1997). TGF-β signalling from cell membrane to nucleus through SMAD proteins. Nature, 390(6659), 465–471. https://doi.org/10.1038/37284

Hynes, R. O. (2002). Integrins: bidirectional, allosteric signaling machines. Cell, 110(6), 673–687. http://www.ncbi.nlm.nih.gov/pubmed/12297042

Hynes, W. L., and Walton, S. L. (2000). Hyaluronidases of Gram-positive bacteria. FEMS Microbiology Letters, 183(2), 201–207. https://doi.org/10.1111/j.1574-6968.2000.tb08958.x

Imler, J.-L. (2003). Biology of Toll receptors: lessons from insects and mammals. Journal of Leukocyte Biology, 75(1), 18–26. https://doi.org/10.1189/jlb.0403160

James-Clark, H. (1867). Conclusive proofs on the animality of the ciliate sponges, and their affinities with the Infusoria Flagellata. The Annals and Magazine of Natural History, 19(109), 13–18. https://doi.org/10.1080/00222936708679703

Janeway, C. A., and Medzhitov, R. (2002). Innate Immune Recognition. Annual Review of Immunology, 20(1), 197–216. https://doi.org/10.1146/annurev.immunol.20.083001.084359

Jinek, M., and Doudna, J. A. (2009). A three-dimensional view of the molecular machinery of RNA interference. Nature, 457(7228), 405–412. https://doi.org/10.1038/nature07755

Käll, L., Krogh, A., and Sonnhammer, E. L. (2004). A Combined Transmembrane Topology and Signal Peptide Prediction Method. Journal of Molecular Biology, 338(5), 1027–1036. https://doi.org/10.1016/j.jmb.2004.03.016

Kanehisa, M., Goto, S., Sato, Y., Furumichi, M., and Tanabe, M. (2012). KEGG for integration and interpretation of large-scale molecular data sets. Nucleic Acids Research, 40(D1), D109–D114. https://doi.org/10.1093/nar/gkr988

Kaneiwa, T., Yamada, S., Mizumoto, S., Montaño, A. M., Mitani, S., and Sugahara, K. (2008). Identification of a Novel Chondroitin Hydrolase in Caenorhabditis elegans. Journal of Biological Chemistry, 283(22), 14971–14979. https://doi.org/10.1074/jbc.M709236200

Katoh, K., and Standley, D. M. (2013). MAFFT Multiple Sequence Alignment Software Version 7: Improvements in Performance and Usability. Molecular Biology and Evolution, 30(4), 772–780. https://doi.org/10.1093/molbev/mst010

Kent, W. J. (2002). BLAT---The BLAST-Like Alignment Tool. Genome Research, 12(4), 656–664. https://doi.org/10.1101/gr.229202

Kent, W. S. (1880). A Manual of the Infusoria. London: David Bogue.

King, N. (2004). The Unicellular Ancestry of Animal Development. Developmental Cell, 7(3), 313–325. https://doi.org/10.1016/j.devcel.2004.08.010

King, N., and Rokas, A. (2017). Embracing Uncertainty in Reconstructing Early Animal Evolution. Current Biology, 27(19), R1081–R1088. https://doi.org/10.1016/j.cub.2017.08.054

King, N., Westbrook, M. J., Young, S. L., Kuo, A., Abedin, M., Chapman, J.,…Rokhsar, D. (2008). The genome of the choanoflagellate Monosiga brevicollis and the origin of metazoans. Nature, 451(7180), 783–788. https://doi.org/10.1038/nature06617

Kingston, R. E. (2001). Preparation of Poly(A) + RNA. In Current Protocols in Molecular Biology (Vol. Chapter 4, p. Unit4.5). Hoboken, NJ, USA: John Wiley & Sons, Inc. https://doi.org/10.1002/0471142727.mb0405s21

Kircher, M., Sawyer, S., and Meyer, M. (2012). Double indexing overcomes inaccuracies in multiplex sequencing on the Illumina platform. Nucleic Acids Research, 40(1), e3–e3. https://doi.org/10.1093/nar/gkr771

Kjellén, L., and Lindahl, U. (1991). Proteoglycans: Structures and Interactions. Annual Review of Biochemistry, 60(1), 443–475. https://doi.org/10.1146/annurev.bi.60.070191.002303

Knoll, A. H. (2011). The Multiple Origins of Complex Multicellularity. Annual Review of Earth and Planetary Sciences, 39(1), 217–239. https://doi.org/10.1146/annurev.earth.031208.100209

Leadbeater, B. S. C. (2015). The choanoflagellates: evolution, biology, and ecology. Cambridge, United Kingdom: Cambridge University Press.

Leadbeater, B. S. C., and McCready, S. M. M. (2000). The flagellates: Historical perspectives. In B. S. C. Leadbeater & J. C. Green (Eds.), The Flagellates: Unity, Diversity and Evolution (pp. 1–26). London: Taylor & Francis Limited.

Lee, R. Y. N., Howe, K. L., Harris, T. W., Arnaboldi, V., Cain, S., Chan, J.,…Sternberg, P. W. (2017). WormBase 2017: molting into a new stage. Nucleic Acids Research. https://doi.org/10.1093/nar/gkx998

Letunic, I., and Bork, P. (2016). Interactive tree of life (iTOL) v3: an online tool for the display and annotation of phylogenetic and other trees. Nucleic Acids Research, 44(W 1), W242–W245. https://doi.org/10.1093/nar/gkw290

Leulier, F., and Lemaitre, B. (2008). Toll-like receptors — taking an evolutionary approach. Nature Reviews Genetics, 9(3), 165–178. https://doi.org/10.1038/nrg2303

Levin, T. C., Greaney, A. J., Wetzel, L., and King, N. (2014). The rosetteless gene controls development in the choanoflagellate S. rosetta. eLife, 3. https://doi.org/10.7554/eLife.04070

Li, H., and Durbin, R. (2009). Fast and accurate short read alignment with Burrows-Wheeler transform. Bioinformatics, 25(14), 1754–1760. https://doi.org/10.1093/bioinformatics/btp324

Li, H., Handsaker, B., Wysoker, A., Fennell, T., Ruan, J., Homer, N.,…Durbin, R. (2009). The Sequence Alignment/Map format and SAMtools. Bioinformatics, 25(16), 2078–2079. https://doi.org/10.1093/bioinformatics/btp352

Li, L. (2003). OrthoMCL: Identification of Ortholog Groups for Eukaryotic Genomes. Genome Research, 13(9), 2178–2189. https://doi.org/10.1101/gr.1224503

Li, W., and Godzik, A. (2006). Cd-hit: a fast program for clustering and comparing large sets of protein or nucleotide sequences. Bioinformatics, 22(13), 1658–1659. https://doi.org/10.1093/bioinformatics/btl158

Liu, H.-Y., Chen, C.-Y., and Hsueh, Y.-P. (2014). Innate immune responses regulate morphogenesis and degeneration: roles of Toll-like receptors and Sarm1 in neurons. Neuroscience Bulletin, 30(4), 645–654. https://doi.org/10.1007/s12264-014-1445-5

Lohse, M., Bolger, A. M., Nagel, A., Fernie, A. R., Lunn, J. E., Stitt, M., and Usadel, B. (2012). RobiNA: a user-friendly, integrated software solution for RNA-Seq-based transcriptomics. Nucleic Acids Research, 40(W1), W622–W627. https://doi.org/10.1093/nar/gks540

Louis, A., Roest Crollius, H., and Robinson-Rechavi, M. (2012). How much does the amphioxus genome represent the ancestor of chordates? Briefings in Functional Genomics, 11(2), 89–95. https://doi.org/10.1093/bfgp/els003

Malik, S., and Roeder, R. G. (2010). The metazoan Mediator co-activator complex as an integrative hub for transcriptional regulation. Nature Reviews Genetics, 11(11), 761–772. https://doi.org/10.1038/nrg2901

Manning, G., Young, S. L., Miller, W. T., and Zhai, Y. (2008). The protist, Monosiga brevicollis, has a tyrosine kinase signaling network more elaborate and diverse than found in any known metazoan. Proceedings of the National Academy of Sciences, 105(28), 9674–9679. https://doi.org/10.1073/pnas.0801314105

Matveyev, A. V., Alves, J. M. P., Serrano, M. G., Lee, V., Lara, A. M., Barton, W. A.,…Buck, G. A. (2017). The Evolutionary Loss of RNAi Key Determinants in Kinetoplastids as a Multiple Sporadic Phenomenon. Journal of Molecular Evolution, 84(2–3), 104–115. https://doi.org/10.1007/s00239-017-9780-1

McFall-Ngai, M., Hadfield, M. G., Bosch, T. C. G., Carey, H. V, Domazet-Lošo, T., Douglas, A. E.,…Wernegreen, J. J. (2013). Animals in a bacterial world, a new imperative for the life sciences. Proceedings of the National Academy of Sciences, 110(9), 3229–3236. https://doi.org/10.1073/pnas.1218525110

Mi, H., Dong, Q., Muruganujan, A., Gaudet, P., Lewis, S., and Thomas, P. D. (2010). PANTHER version 7: improved phylogenetic trees, orthologs and collaboration with the Gene Ontology Consortium. Nucleic Acids Research, 38(Database issue), D204–10. https://doi.org/10.1093/nar/gkp1019

Miller, D. J., Hemmrich, G., Ball, E. E., Hayward, D. C., Khalturin, K., Funayama, N.,…Bosch, T. C. (2007). The innate immune repertoire in Cnidaria - ancestral complexity and stochastic gene loss. Genome Biology, 8(4), R59. https://doi.org/10.1186/gb-2007-8-4-r59

Mizumoto, S., Ikegawa, S., and Sugahara, K. (2013). Human Genetic Disorders Caused by Mutations in Genes Encoding Biosynthetic Enzymes for Sulfated Glycosaminoglycans. Journal of Biological Chemistry, 288(16), 10953–10961. https://doi.org/10.1074/jbc.R112.437038

Moroz, L. L., Kocot, K. M., Citarella, M. R., Dosung, S., Norekian, T. P., Povolotskaya, I. S.,…Kohn, A. B. (2014). The ctenophore genome and the evolutionary origins of neural systems. Nature, 510(7503), 109–114. https://doi.org/10.1038/nature13400

Munger, J. S., Harpel, J. G., Gleizes, P.-E., Mazzieri, R., Nunes, I., and Rifkin, D. B. (1997). Latent transforming growth factor-β: Structural features and mechanisms of activation. Kidney International, 51(5), 1376–1382. https://doi.org/10.1038/ki.1997.188

Narbonne, G. M. (2005). The Ediacara Biota: Neoproterozoic Origin of Animals and Their Ecosystems. Annual Review of Earth and Planetary Sciences, 33(1), 421–442. https://doi.org/10.1146/annurev.earth.33.092203.122519

NCBI Resource Coordinators. (2017). Database Resources of the National Center for Biotechnology Information. Nucleic Acids Research, 45(D1), D12–D17. https://doi.org/10.1093/nar/gkw1071

Nguyen, T. A., Cissé, O. H., Yun Wong, J., Zheng, P., Hewitt, D., Nowrousian, M.,…Jedd, G. (2017). Innovation and constraint leading to complex multicellularity in the Ascomycota. Nature Communications, 8, 14444. https://doi.org/10.1038/ncomms14444

Nichols, S. A., Roberts, B. W., Richter, D. J., Fairclough, S. R., and King, N. (2012). Origin of metazoan cadherin diversity and the antiquity of the classical cadherin/ -catenin complex. Proceedings of the National Academy of Sciences, 109(32), 13046–13051. https://doi.org/10.1073/pnas.1120685109

O’Malley, M. A., Wideman, J. G., and Ruiz-Trillo, I. (2016). Losing Complexity: The Role of Simplification in Macroevolution. Trends in Ecology & Evolution, 31(8), 608–621. https://doi.org/10.1016/j.tree.2016.04.004

O’Neill, L. A. J. (2008). The interleukin-1 receptor/Toll-like receptor superfamily: 10 years of progress. Immunological Reviews, 226(1), 10–18. https://doi.org/10.1111/j.1600-065X.2008.00701.x

Ohya, T., and Kaneko, Y. (1970). Novel hyaluronidase from streptomyces. Biochimica et Biophysica Acta (BBA) - Enzymology, 198(3), 607–609. https://doi.org/10.1016/0005-2744(70)90139-7

Paradis, E., Claude, J., and Strimmer, K. (2004). APE: Analyses of Phylogenetics and Evolution in R language. Bioinformatics, 20(2), 289–290. https://doi.org/10.1093/bioinformatics/btg412

Park, Y., Cho, S., and Linhardt, R. J. (1997). Exploration of the action pattern of Streptomyces hyaluronate lyase using high-resolution capillary electrophoresis. Biochimica et Biophysica Acta (BBA) - Protein Structure and Molecular Enzymology, 1337(2), 217–226. https://doi.org/10.1016/S0167-4838(96)00167-7

Payne, S. H., and Loomis, W. F. (2006). Retention and Loss of Amino Acid Biosynthetic Pathways Based on Analysis of Whole-Genome Sequences. Eukaryotic Cell, 5(2), 272–276. https://doi.org/10.1128/EC.5.2.272-276.2006

Peña, J. F., Alié, A., Richter, D. J., Wang, L., Funayama, N., and Nichols, S. A. (2016). Conserved expression of vertebrate microvillar gene homologs in choanocytes of freshwater sponges. EvoDevo, 7(1), 13. https://doi.org/10.1186/s13227-016-0050-x

Peterson, K. J., Lyons, J. B., Nowak, K. S., Takacs, C. M., Wargo, M. J., and McPeek, M. A. (2004). Estimating metazoan divergence times with a molecular clock. Proceedings of the National Academy of Sciences, 101(17), 6536–6541. https://doi.org/10.1073/pnas.0401670101

Philippe, H., Derelle, R., Lopez, P., Pick, K., Borchiellini, C., Boury-Esnault, N.,…Manuel, M. (2009). Phylogenomics Revives Traditional Views on Deep Animal Relationships. Current Biology, 19(8), 706–712. https://doi.org/10.1016/j.cub.2009.02.052

Pincus, D., Letunic, I., Bork, P., and Lim, W. A. (2008). Evolution of the phospho-tyrosine signaling machinery in premetazoan lineages. Proceedings of the National Academy of Sciences, 105(28), 9680–9684. https://doi.org/10.1073/pnas.0803161105

Posas, F., Wurgler-Murphy, S. M., Maeda, T., Witten, E. A., Thai, T. C., and Saito, H. (1996). Yeast HOG1 MAP Kinase Cascade Is Regulated by a Multistep Phosphorelay Mechanism in the SLN1-YPD1–SSK1 “Two-Component” Osmosensor. Cell, 86(6), 865–875. https://doi.org/10.1016/S0092-8674(00)80162-2

Prochnik, S. E., Umen, J., Nedelcu, A. M., Hallmann, A., Miller, S. M., Nishii, I.,…Rokhsar, D. S. (2010). Genomic Analysis of Organismal Complexity in the Multicellular Green Alga Volvox carteri. Science, 329(5988), 223–226. https://doi.org/10.1126/science.1188800

Punta, M., Coggill, P. C., Eberhardt, R. Y., Mistry, J., Tate, J., Boursnell, C.,…Finn, R. D. (2012). The Pfam protein families database. Nucleic Acids Research, 40(D1), D290–D301. https://doi.org/10.1093/nar/gkr1065

Putnam, N. H., Srivastava, M., Hellsten, U., Dirks, B., Chapman, J., Salamov, A.,…Rokhsar, D. S. (2007). Sea Anemone Genome Reveals Ancestral Eumetazoan Gene Repertoire and Genomic Organization. Science, 317(5834), 86–94. https://doi.org/10.1126/science.1139158

R Core Team, T. (2017). R: A language and environment for statistical computing. Vienna, Austria: R Foundation for Statistical Computing. https://www.r-project.org/

Richards, G. S., and Degnan, B. M. (2009). The Dawn of Developmental Signaling in the Metazoa. Cold Spring Harbor Symposia on Quantitative Biology, 74, 81–90. https://doi.org/10.1101/sqb.2009.74.028

Richards, G. S., Simionato, E., Perron, M., Adamska, M., Vervoort, M., and Degnan, B. M. (2008). Sponge Genes Provide New Insight into the Evolutionary Origin of the Neurogenic Circuit. Current Biology, 18(15), 1156–1161. https://doi.org/10.1016/j.cub.2008.06.074

Richter, D. J. (2013). The gene content of diverse choanoflagellates illuminates animal origins. University of California, Berkeley. https://escholarship.org/uc/item/7xc2p94p

Richter, D. J., and King, N. (2013). The Genomic and Cellular Foundations of Animal Origins. Annual Review of Genetics. https://doi.org/10.1146/annurev-genet-111212-133456

Richter, D. J., and Nitsche, F. (2016). Choanoflagellatea. In Handbook of the Protists (pp. 1–19). Cham: Springer International Publishing. https://doi.org/10.1007/978-3-319-32669-6_5-1

Riesgo, A., Farrar, N., Windsor, P. J., Giribet, G., and Leys, S. P. (2014). The Analysis of Eight Transcriptomes from All Poriferan Classes Reveals Surprising Genetic Complexity in Sponges. Molecular Biology and Evolution, 31(5), 1102–1120. https://doi.org/10.1093/molbev/msu057

Roberts, A., and Pachter, L. (2012). Streaming fragment assignment for real-time analysis of sequencing experiments. Nature Methods, 10(1), 71–73. https://doi.org/10.1038/nmeth.2251

Robinson, P. N., Wollstein, A., Böhme, U., and Beattie, B. (2004). Ontologizing gene-expression microarray data: characterizing clusters with Gene Ontology. Bioinformatics, 20(6), 979–981. https://doi.org/10.1093/bioinformatics/bth040

Rokas, A. (2008). The Origins of Multicellularity and the Early History of the Genetic Toolkit For Animal Development. Annual Review of Genetics, 42(1), 235–251. https://doi.org/10.1146/annurev.genet.42.110807.091513

RStudio Team, T. (2016). RStudio: Integrated Development for R. Boston, MA: RStudio, Inc. http://www.rstudio.com/

Rubin, G. M., Yandell, M. D., Wortman, J. R., Gabor Miklos, G. L., Nelson, C. R., Hariharan, I. K.,…Lewis, S. (2000). Comparative genomics of the eukaryotes. Science, 287(5461), 2204–2215. https://doi.org/10.1126/science.287.5461.2204

Saldanha, A. J. (2004). Java Treeview--extensible visualization of microarray data. Bioinformatics, 20(17), 3246–3248. https://doi.org/10.1093/bioinformatics/bth349

Schenkelaars, Q., Pratlong, M., Kodjabachian, L., Fierro-Constain, L., Vacelet, J., Le Bivic, A.,…Borchiellini, C. (2017). Animal multicellularity and polarity without Wnt signaling. Scientific Reports, 7(1), 15383. https://doi.org/10.1038/s41598-017-15557-5

Schroeder, A., Mueller, O., Stocker, S., Salowsky, R., Leiber, M., Gassmann, M.,…Ragg, T. (2006). The RIN: an RNA integrity number for assigning integrity values to RNA measurements. BMC Molecular Biology, 7(1), 3. https://doi.org/10.1186/1471-2199-7-3

Sebé-Pedrós, A., Ariza-Cosano, A., Weirauch, M. T., Leininger, S., Yang, A., Torruella, G.,…Ruiz-Trillo, I. (2013a). Early evolution of the T-box transcription factor family. Proceedings of the National Academy of Sciences, 110(40), 16050–16055. https://doi.org/10.1073/pnas.1309748110

Sebé-Pedrós, A., Ballaré, C., Parra-Acero, H., Chiva, C., Tena, J. J., Sabidó, E.,…Ruiz-Trillo, I. (2016a). The Dynamic Regulatory Genome of Capsaspora and the Origin of Animal Multicellularity. Cell, 165(5), 1224–1237. https://doi.org/10.1016/j.cell.2016.03.034

Sebé-Pedrós, A., de Mendoza, A., Lang, B. F., Degnan, B. M., Ruiz-Trillo, I., Sebe-Pedros, A.,…Ruiz-Trillo, I. (2011). Unexpected Repertoire of Metazoan Transcription Factors in the Unicellular Holozoan Capsaspora owczarzaki. Molecular Biology and Evolution, 28(3), 1241–1254. https://doi.org/10.1093/molbev/msq309

Sebé-Pedrós, A., Degnan, B. M., and Ruiz-Trillo, I. (2017). The origin of Metazoa: a unicellular perspective. Nature Reviews Genetics, 18(8), 498–512. https://doi.org/10.1038/nrg.2017.21

Sebé-Pedrós, A., Irimia, M., del Campo, J., Parra-Acero, H., Russ, C., Nusbaum, C.,…Ruiz-Trillo, I. (2013b). Regulated aggregative multicellularity in a close unicellular relative of metazoa. eLife, 2. https://doi.org/10.7554/eLife.01287

Sebé-Pedrós, A., Peña, M. I., Capella-Gutiérrez, S., Antó, M., Gabaldón, T., Ruiz-Trillo, I., and Sabidó, E. (2016b). High-Throughput Proteomics Reveals the Unicellular Roots of Animal Phosphosignaling and Cell Differentiation. Developmental Cell, 39(2), 186–197. https://doi.org/10.1016/j.devcel.2016.09.019

Sebé-Pedrós, A., Roger, A. J., Lang, F. B., King, N., and Ruiz-Trillo, I. (2010). Ancient origin of the integrin-mediated adhesion and signaling machinery. Proceedings of the National Academy of Sciences, 107(22), 10142–10147. https://doi.org/10.1073/pnas.1002257107

Sebé-Pedrós, A., and Ruiz-Trillo, I. (2010). Integrin-mediated adhesion complex. Communicative & Integrative Biology, 3(5), 475–477. https://doi.org/10.4161/cib.3.5.12603

Sebé-Pedrós, A., Zheng, Y., Ruiz-Trillo, I., and Pan, D. (2012). Premetazoan Origin of the Hippo Signaling Pathway. Cell Reports, 1(1), 13–20. https://doi.org/10.1016/j.celrep.2011.11.004

Sethman, C. R., and Hawiger, J. (2013). The Innate Immunity Adaptor SARM Translocates to the Nucleus to Stabilize Lamins and Prevent DNA Fragmentation in Response to Pro-Apoptotic Signaling. PLoS ONE, 8(7), e70994. https://doi.org/10.1371/journal.pone.0070994

Shabalina, S., and Koonin, E. (2008). Origins and evolution of eukaryotic RNA interference. Trends in Ecology & Evolution, 23(10), 578–587. https://doi.org/10.1016/j.tree.2008.06.005

Simakov, O., and Kawashima, T. (2017). Independent evolution of genomic characters during major metazoan transitions. Developmental Biology, 427(2), 179–192. https://doi.org/10.1016/j.ydbio.2016.11.012

Simão, F. A., Waterhouse, R. M., Ioannidis, P., Kriventseva, E. V., and Zdobnov, E. M. (2015). BUSCO: assessing genome assembly and annotation completeness with single-copy orthologs. Bioinformatics, 31(19), 3210–3212. https://doi.org/10.1093/bioinformatics/btv351

Snell, E. A., Brooke, N. M., Taylor, W. R., Casane, D., Philippe, H., and Holland, P. W. (2006). An unusual choanoflagellate protein released by Hedgehog autocatalytic processing. Proceedings of the Royal Society B: Biological Sciences, 273(1585), 401–407. https://doi.org/10.1098/rspb.2005.3263

Song, X., Jin, P., Qin, S., Chen, L., and Ma, F. (2012). The Evolution and Origin of Animal Toll-Like Receptor Signaling Pathway Revealed by Network-Level Molecular Evolutionary Analyses. PLoS ONE, 7(12), e51657. https://doi.org/10.1371/journal.pone.0051657

Sorokin, L. (2010). The impact of the extracellular matrix on inflammation. Nature Reviews Immunology, 10(10), 712–723. https://doi.org/10.1038/nri2852

Srivastava, M., Simakov, O., Chapman, J., Fahey, B., Gauthier, M. E. A., Mitros, T.,…Rokhsar, D. S. (2010). The Amphimedon queenslandica genome and the evolution of animal complexity. Nature, 466(7307), 720–726. https://doi.org/10.1038/nature09201

Stamatakis, A. (2014). RAxML version 8: a tool for phylogenetic analysis and post-analysis of large phylogenies. Bioinformatics, 30(9), 1312–1313. https://doi.org/10.1093/bioinformatics/btu033

Starcevic, A., Akthar, S., Dunlap, W. C., Shick, J. M., Hranueli, D., Cullum, J., and Long, P. F. (2008). Enzymes of the shikimic acid pathway encoded in the genome of a basal metazoan, Nematostella vectensis, have microbial origins. Proceedings of the National Academy of Sciences, 105(7), 2533–2537. https://doi.org/10.1073/pnas.0707388105

Stern, R., and Jedrzejas, M. J. (2006). Hyaluronidases: Their Genomics, Structures, and Mechanisms of Action. Chemical Reviews, 106(3), 818–839. https://doi.org/10.1021/cr050247k

Suga, H., Chen, Z., de Mendoza, A., Sebé-Pedrós, A., Brown, M. W., Kramer, E.,…Ruiz-Trillo, I. (2013). The Capsaspora genome reveals a complex unicellular prehistory of animals. Nature Communications, 4. https://doi.org/10.1038/ncomms3325

Sullivan, J. C., and Finnerty, J. R. (2007). A surprising abundance of human disease genes in a simple “basal” animal, the starlet sea anemone (Nematostella vectensis). Genome, 50(7), 689–92. https://doi.org/10.1139/g07-045

Sullivan, J. C., Kalaitzidis, D., Gilmore, T. D., and Finnerty, J. R. (2007). Rel homology domain-containing transcription factors in the cnidarian Nematostella vectensis. Development Genes and Evolution, 217(1), 63–72. https://doi.org/10.1007/s00427-006-0111-6

Takami, H., Taniguchi, T., Arai, W., Takemoto, K., Moriya, Y., and Goto, S. (2016). An automated system for evaluation of the potential functionome: MAPLE version 2.1.0. DNA Research, 23(5), 467–475. https://doi.org/10.1093/dnares/dsw030

Thomas, P. D. (2003). PANTHER: a browsable database of gene products organized by biological function, using curated protein family and subfamily classification. Nucleic Acids Research, 31(1), 334–341. https://doi.org/10.1093/nar/gkg115

Thomas, P. D., Kejariwal, A., Guo, N., Mi, H., Campbell, M. J., Muruganujan, A., and Lazareva-Ulitsky, B. (2006). Applications for protein sequence-function evolution data: mRNA/protein expression analysis and coding SNP scoring tools. Nucleic Acids Research, 34(Web Server), W645–W650. https://doi.org/10.1093/nar/gkl229

Torruella, G., de Mendoza, A., Grau-Bové, X., Antó, M., Chaplin, M. A., del Campo, J.,…Ruiz-Trillo, I. (2015). Phylogenomics Reveals Convergent Evolution of Lifestyles in Close Relatives of Animals and Fungi. Current Biology, 25(18), 2404–2410. https://doi.org/10.1016/j.cub.2015.07.053

Uniprot Consortium, T. (2017). UniProt: the universal protein knowledgebase. Nucleic Acids Research, 45(D1), D158–D169. https://doi.org/10.1093/nar/gkw1099

Urbach, J. M., and Ausubel, F. M. (2017). The NBS-LRR architectures of plant R-proteins and metazoan NLRs evolved in independent events. Proceedings of the National Academy of Sciences, 114(5), 1063–1068. https://doi.org/10.1073/pnas.1619730114

Usui, T., Shima, Y., Shimada, Y., Hirano, S., Burgess, R. W., Schwarz, T. L.,…Uemura, T. (1999). Flamingo, a Seven-Pass Transmembrane Cadherin, Regulates Planar Cell Polarity under the Control of Frizzled. Cell, 98(5), 585–595. https://doi.org/10.1016/S0092-8674(00)80046-X

van der Vaart, A. W. (1998). Asymtotic statistics. New York: Cambridge University Press.

van Dongen, S., and Abreu-Goodger, C. (2012). Using MCL to Extract Clusters from Networks. In Methods Mol. Biol. (Vol. 804, pp. 281–295). https://doi.org/10.1007/978-1-61779-361-5_15

von Moltke, J., Ayres, J. S., Kofoed, E. M., Chavarría-Smith, J., and Vance, R. E. (2013). Recognition of Bacteria by Inflammasomes. Annual Review of Immunology, 31(1), 73–106. https://doi.org/10.1146/annurev-immunol-032712-095944

Waterhouse, A. M., Procter, J. B., Martin, D. M. A., Clamp, M., and Barton, G. J. (2009). Jalview Version 2--a multiple sequence alignment editor and analysis workbench. Bioinformatics, 25(9), 1189–1191. https://doi.org/10.1093/bioinformatics/btp033

Wickham, H. (2009). ggplot2: Elegant Graphics for Data Analysis. New York: Springer-Verlag. http://ggplot2.org

Wiens, M., Korzhev, M., Krasko, A., Thakur, N. L., Perović-Ottstadt, S., Breter, H. J.,…Müller, W. E. G. (2005). Innate Immune Defense of the Sponge Suberites domuncula against Bacteria Involves a MyD88-dependent Signaling Pathway. Journal of Biological Chemistry, 280(30), 27949–27959. https://doi.org/10.1074/jbc.M504049200

Wolf, Y. I., and Koonin, E. V. (2013). Genome reduction as the dominant mode of evolution. BioEssays, 35(9), 829–837. https://doi.org/10.1002/bies.201300037

Woznica, A., Cantley, A. M., Beemelmanns, C., Freinkman, E., Clardy, J., and King, N. (2016). Bacterial lipids activate, synergize, and inhibit a developmental switch in choanoflagellates. Proceedings of the National Academy of Sciences, 113(28), 7894–7899. https://doi.org/10.1073/pnas.1605015113

Woznica, A., Gerdt, J. P., Hulett, R. E., Clardy, J., and King, N. (2017). Mating in the Closest Living Relatives of Animals Is Induced by a Bacterial Chondroitinase. Cell, 170(6), 1175–1183.e11. https://doi.org/10.1016/j.cell.2017.08.005

Wu, X., Wu, F.-H., Wang, X., Wang, L., Siedow, J. N., Zhang, W., and Pei, Z.-M. (2014). Molecular evolutionary and structural analysis of the cytosolic DNA sensor cGAS and STING. Nucleic Acids Research, 42(13), 8243–8257. https://doi.org/10.1093/nar/gku569

Yamagata, T., Saito, H., Habuchi, O., and Suzuki, S. (1968). Purification and properties of bacterial chondroitinases and chondrosulfatases. Journal of Biological Chemistry, 243(7), 1523–35. http://www.ncbi.nlm.nih.gov/pubmed/5647268

Yang, X., Dorman, K. S., and Aluru, S. (2010). Reptile: representative tiling for short read error correction. Bioinformatics, 26(20), 2526–2533. https://doi.org/10.1093/bioinformatics/btq468

Yuen, B., Bayes, J. M., and Degnan, S. M. (2014). The Characterization of Sponge NLRs Provides Insight into the Origin and Evolution of This Innate Immune Gene Family in Animals. Molecular Biology and Evolution, 31(1), 106–120. https://doi.org/10.1093/molbev/mst174

